# Switching of RNA splicing regulators in immature neuroblasts: a key step in adult neurogenesis

**DOI:** 10.1101/2023.03.12.532290

**Authors:** Corentin Bernou, Marc-André Mouthon, Mathieu Daynac, Thierry Kortulewski, Benjamin Demaille, Vilma Barroca, Sébastien Couillard-Despres, Nathalie Dechamps, Véronique Ménard, Léa Bellenger, Christophe Antoniewski, Alexandra Chicheportiche, François D. Boussin

**Affiliations:** Université Paris Cité, Inserm, CEA, Stabilité Génétique Cellules Souches et Radiations, LRP/iRCM, F-92265, Fontenay-aux-Roses, France; Université Paris-Saclay, Inserm, CEA, Stabilité Génétique Cellules Souches et Radiations, LRP/iRCM, F-92265, Fontenay-aux-Roses, France; Inserm, ARTbio Bioinformatics Analysis Facility, Sorbonne Université, CNRS, Institut de Biologie Paris Seine, Paris, France; ARTbio Bioinformatics Analysis Facility, Sorbonne Université, CNRS, Institut de Biologie Paris Seine, Paris, France; Spinal Cord Injury and Tissue Regeneration Center Salzburg (SCI-TReCS), Paracelsus Medical University, 5020 Salzburg, Austria; Institute of Experimental Neuroregeneration, Paracelsus Medical University, 5020 Salzburg, Austria; Austrian Cluster for Tissue Regeneration, 1200 Vienna, Austria

**Keywords:** Subventricular zone, adult neurogenesis, transcriptomic analyses, immature neuroblasts

## Abstract

The lateral wall of the subventricular zone harbors neural stem cells (NSC, B cells) which generate proliferating transient-amplifying progenitors (TAP, C cells) that ultimately give rise to neuroblasts (NB, A cells). Molecular profiling at the single cell level struggles to distinguish these different cell types. Here, we combined transcriptome analyses of FACS-sorted cells and single-cell RNAseq to demonstrate the existence of an abundant, clonogenic and multipotent population of immature neuroblasts (iNB cells) at the transition between TAP and migrating NB (mNB). iNB are reversibly engaged in neuronal differentiation. Indeed, they keep molecular features of both undifferentiated progenitors, plasticity and unexpected regenerative properties. Strikingly, they undergo important progressive molecular switches, including changes in the expression of splicing regulators leading to their differentiation in mNB subdividing them into 2 subtypes, iNB1 and iNB2. Due to their plastic properties, iNB could represent a new target for regenerative therapy of brain damage.

## Introduction

New neurons are constantly produced from neural stem cells (NSC) throughout life in two restricted neurogenic areas of the adult mammalian brain: the sub-granular zone (SGZ) of the hippocampus and the subventricular zone (SVZ) ^1^. The lateral wall (LW) of the SVZ, adjacent to the striatum, is the largest germinal zone of the adult mouse brain, in which NSC (type B1 cells) are produced during the embryonic development and are mostly in a quiescent state thereafter^2,3^. Upon activation, NSC divide and give rise to rapidly dividing transit-amplifying cells (TAP, type C cells), which ultimately produce neuroblasts (NB, type A cells). These neuroblasts migrate out of the SVZ anteriorly along blood vessels and the rostral migratory stream (RMS) to reach the olfactory bulb (OB) where they integrate and differentiate into various subtypes of interneurons required to fine-tune odor discrimination ^1^. The diversity of the OB interneurons depends on the intrinsic regional identity of NSC in the SVZ along the anterior-posterior and medial-lateral axes^1,4^.

Besides their neurogenic fate, NSC and neural progenitors present in the SVZ are also proficient to produce astrocytes and oligodendrocytes that migrate toward the corpus callosum, the RMS and the striatum in the normal^5,6,7^ and injured adult brain^8,9^.

Transcriptomic analyses using different combinations of cell surface markers and transgenic mice models to isolate SVZ cells have provided important initial insights into stem cell quiescence and have highlighted consecutive transitions during neuronal lineage progression ^2,10–13^. Since then, the single cell RNA-sequencing has revolutionized the field and has made it possible to precisely elucidate the transcriptome of SVZ cells present in the LW^14–17^ and in the septal wall which also harbours NSC niches^18^.

We have previously developed a FACS-based method, using a combination of three surface markers (LeX, EGF receptor, CD24), allowing to sort five distinct SVZ cell populations: quiescent NSC (LeX-EGFR-CD24-, hereafter referred to as sorted-qNSC, s-qNSC), activated NSC (LeX+EGFR+CD24-, referred to as sorted, s-aNSC), transit amplifying cells (LeX^−^EGFR^+^CD24^−^, referred to as sorted, s-TAP), immature (LeX^−^ EGFR^+^CD24^low^, referred to as sorted, s-iNB) and migrating neuroblasts (LeX^−^EGFR^−^CD24^low^, referred to as sorted, s-mNB) from the adult mouse brain^19,20^. Importantly, whereas s-iNB presented neuroblast markers, e.g. CD24 and doublecortin (DCX), s-iNB gathered the most abundant population among SVZ cycling progenitors (representing approximately 4.3% of total SVZ cells *vs* 1% s-aNSC and 2.5% s-TAP)^20^.

We have previously shown that following a genotoxic stress (exposure to ionizing radiation) inducing the depletion of cycling progenitors, s-qNSC exit quiescence restoring successively s-aNSC, s-TAP, s-iNB and finally s-mNB^20^.

Transcriptomic studies revealed that age-related SVZ neurogenesis decline is associated with TGFß signaling and specific cell cycle regulation changes in s-aNSC. A progressive lengthening of aNSC G1 phase with age leads to a decrease in the production of s-TAP, s-iNB and s-mNB ^20–22^. However, despite the decrease of neurogenesis with aging the s-iNB remained more abundant than s-aNSC and s-TAP in the aged brain ^21^.

Studies of SVZ neural progenitors have been essentially focused on the characterization of the molecular mechanisms regulating quiescence, activation and the regenerative potential of aNSC. Here, we combined transcriptome analyses using DNA microarrays on FAC-sorted cells and single-cell RNAseq to demonstrate that iNB form a molecularly distinct class of SVZ cells, exhibiting molecular features of both neural progenitors and neuroblasts while keeping unexpected regenerative capacity and plasticity. We identified important and sequential molecular switches occurring in these cells before their final differentiation in migrating neuroblasts. Altogether, these data led us to propose a revision of the current model of adult neurogenesis.

## Results

### Regenerative potential and multipotency of sorted iNB *in vitro and in vivo*

We have previously shown that s-iNB, which expressed the neuroblast markers CD24 and DCX^20^ and markers of neural progenitors such as Mash1^20^, Dlx2 (Fig.S1A) and SOX2 (Fig.S1B) had a clonogenic potential and were able to proliferate *in vitro* for weeks in normoxic conditions (20%O2) unlike s-mNB ^20^. However, s-iNB exhibited reduced clonogenic potential compared to that of s-aNSC and s-TAP (Fig.1A). It has been previously reported that hypoxic conditions promoted SVZ NSC proliferation and neurogenesis^23,24^. Hypoxic conditions (4%O2) had no effects on s-mNB, but increased the proliferation capacities of s-aNSC, s-TAP and s-iNB, which reached similar growth rates, making them undistinguishable based on this endpoint (Fig.1A). Hypoxic conditions also increased the clonogenic potentials of both s-TAP and s-iNB but not that of s-aNSC (Fig. 1B). In these conditions, the clonogenic potential of s-TAP reached similar values as s-aNSC, but that of s-iNB was still lower (Fig.1B).

**Figure 1:**
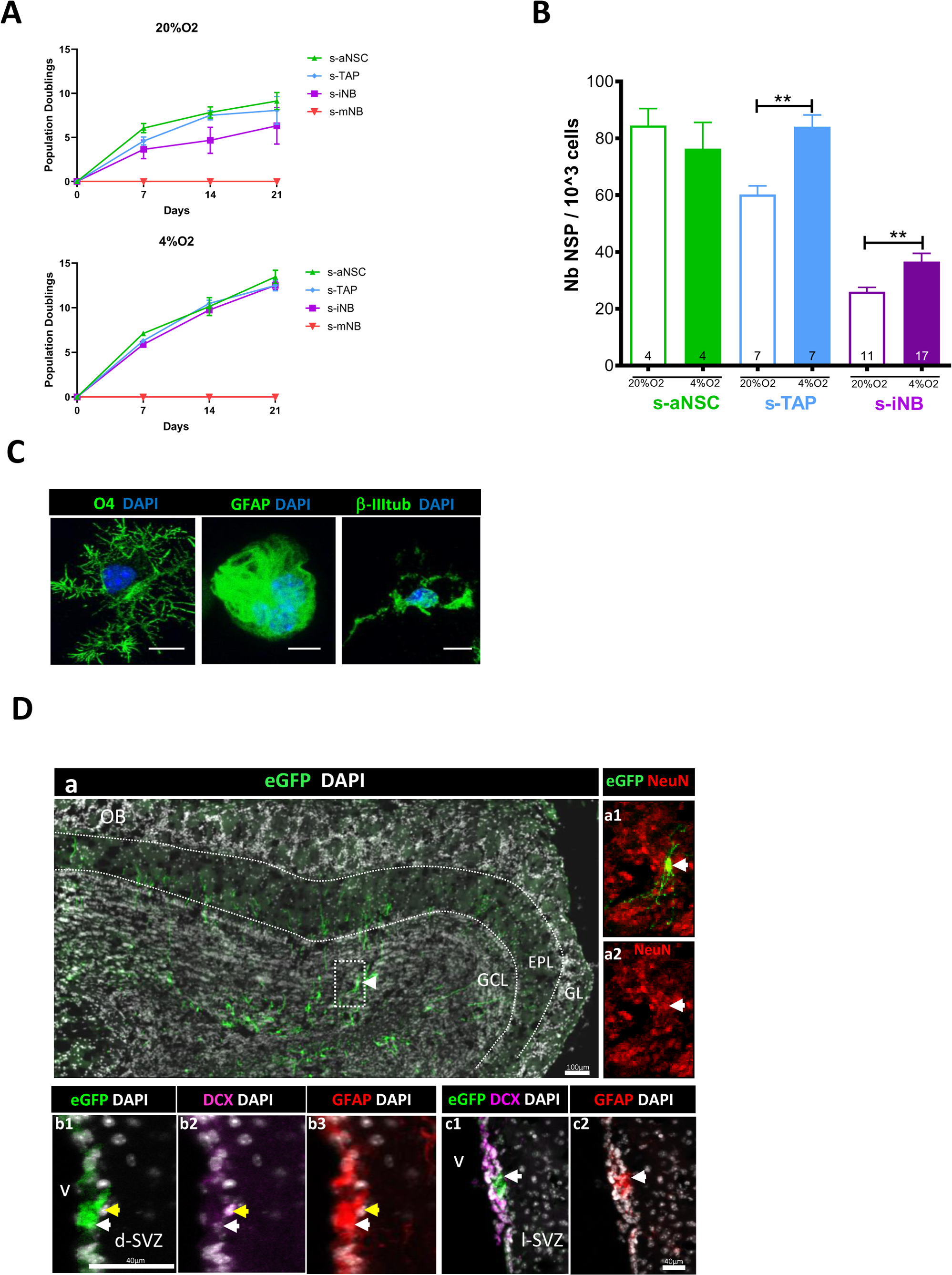
Plasticity of iNB *in vitro* and *in vivo*. s-aNSC, s-TAP, s-iNB, and s-mNB were sorted from the SVZ of 2-month old C57Bl/6 mice. **(A)** Population doublings (PD). Data were obtained from 3 independent experiments (n=8). **(B)** Clonogenicity assay. Results were obtained from 2 independent experiments. The number inside the bars indicated the number of microplate wells analysed (mean ± SEM). **(C)** Differentiation assay. Representative images of immunofluorescence of freshly sorted iNB cells cultured in oligodendrocyte, astrocyte, or neuronal differentiation medium, and stained for NG2 and CNPase, GFAP and CD133 or βIII-Tubulin and doublecortin (DCX) expressions, respectively. *p<0.05, **p<0.01, ***p<0.001. Scale bar: 20µm. **(D)** EGFP-positive s-iNB were isolated from β−actin:eGFP mice and transplanted unilaterally at 3 injection points at proximity of the dSVZ/RMS of recipient C57Bl/6J mice. Transplanted brains were analysed five weeks later by immunostaining. **(a)** Detection of eGFP+ cells in the granule cell layer and the external cell layer of the olfactory bulb of a mouse transplanted. **(a1-a2)** a2 high magnification of the inset (dotted line in a) showing eGFP+ NeuN+ cells (white arrows). **(b-c)** Detection of eGFP^+^ GFAP^+^ DCX^+^ cells (yellow arrows) and eGFP+ GFAP+ (white arrows) in the dorsal (b2) and lateral (c2) SVZ. GCL:granule cell layer, EPL : external plexiform layer, GL: glomerular layer, lSVZ: lateral SVZ, dSVZ: dorsal SVZ, Scale bars= 40µm or 100µm.

Quite surprisingly, freshly isolated s-iNB had the capacity to generate the three neural lineages, neurons, astrocytes and oligodendrocytes, when plated for 5–7 days in the appropriate differentiation media, similarly as s-aNSC and s-TAP (Fig.1C).

Thus, to assess the regenerative potential of these immature neuroblasts *in vivo*, eGFP^+^ iNB were sorted from β−actin:eGFP mice, transplanted into the left striatum of recipient C57Bl/6 mice and compared to eGFP^+^s-mNB and eGFP^+^s-NSC/TAP. Five weeks after transplantation, eGFP^+^ cells were detected in different regions of interest (Table 1) in the brains of mice transplanted with eGFP^+^s-NSC/TAP or eGFP^+^s-iNB, whereas no eGFP+ were observed after transplantation of eGFP^+^s-mNB even at the injection site (IS) (Table 1). Excepted in one out of 3, animals transplanted with eGFP^+^s-iNB presented high levels of eGFP^+^ cells in the OB with levels equivalent or superior to animals transplanted with eGFP^+^s-NSC/TAP or eGFP^+^s-iNB (Fig. 1D). EGFP^+^ cells were detected in the rostral migratory stream (Fig. S1A) as well as in the granular cell layer (GCL) and the external plexiform layer (EPL) of the OB where they differentiated into NeuN^+^ neurons in 2/3 animals transplanted with eGFP^+^s-iNB (Fig.1Da) and in 3/3 animals transplanted with EGFP^+^s-NSC/TAP (Table 1).

**Table 1:**
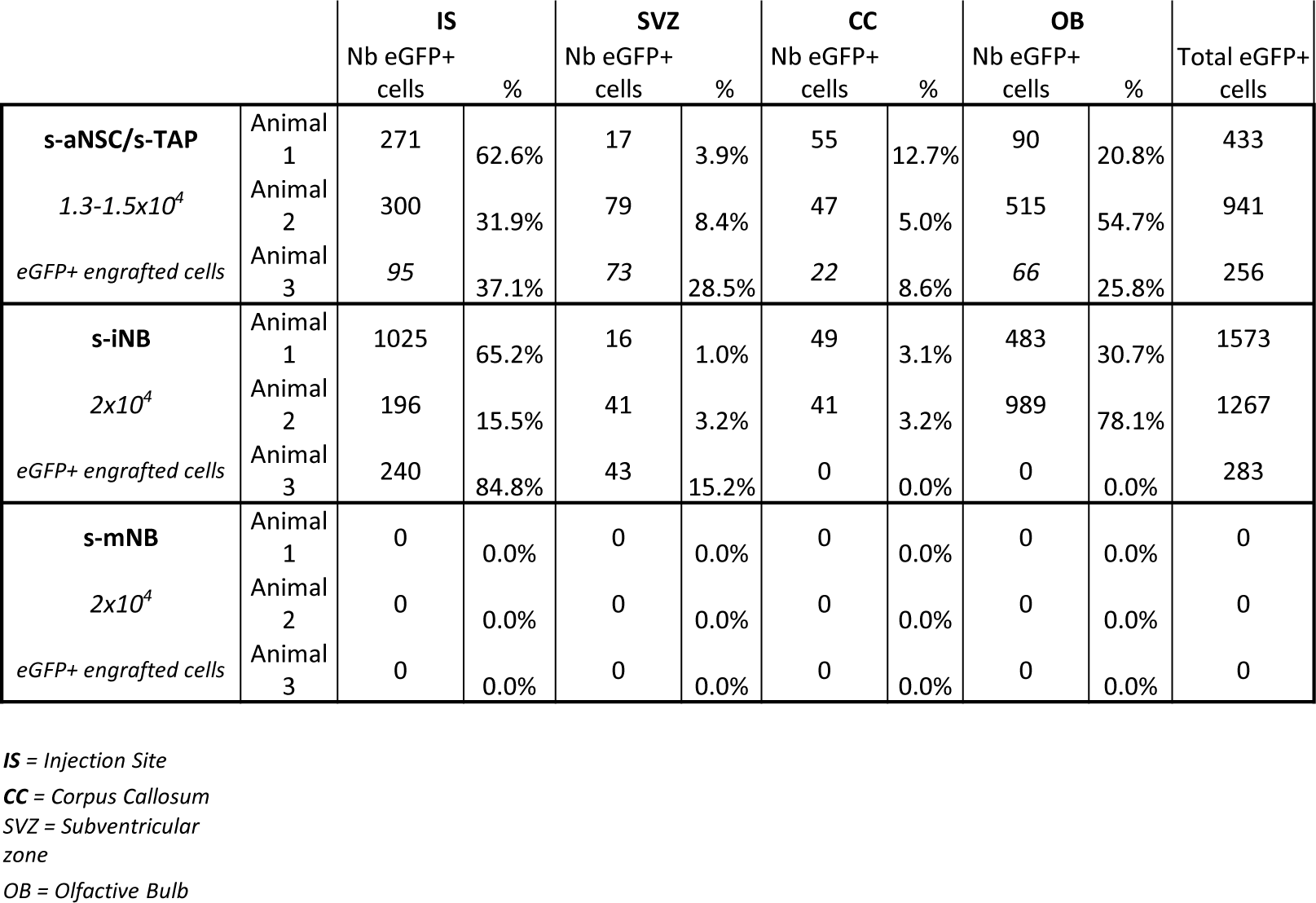
Transplantations of eGFP^+^s-iNB, eGFP^−^s-iNB and eGFP^+^s-mNB freshly isolated from β−actin:eGFP mice model. Table recapitulating the immunohistological analyses of the numbers of eGFP^+^ cells in different regions in recipient C57Bl6/J mice brains, 5 weeks after transplantation.

|  |  | IS |  | SVZ |  | CC |  | OB |  | Total eGFP+ |
| --- | --- | --- | --- | --- | --- | --- | --- | --- | --- | --- |
|  |  | Nb eGFP+<br>cells | % | Nb eGFP+<br>cells | % | Nb eGFP+<br>cells | % | Nb eGFP+<br>cells | % | cells |
| **s-aNSC/s-TAP**<br>$1.3-1.5 \times 10^4$<br>*eGFP+ engrafted cells* | Animal 1 | 271 | 62.6% | 17 | 3.9% | 55 | 12.7% | 90 | 20.8% | 433 |
|  | Animal 2 | 300 | 31.9% | 79 | 8.4% | 47 | 5.0% | 515 | 54.7% | 941 |
|  | Animal 3 | 95 | 37.1% | 73 | 28.5% | 22 | 8.6% | 66 | 25.8% | 256 |
| **s-iNB**<br>$2 \times 10^4$<br>*eGFP+ engrafted cells* | Animal 1 | 1025 | 65.2% | 16 | 1.0% | 49 | 3.1% | 483 | 30.7% | 1573 |
|  | Animal 2 | 196 | 15.5% | 41 | 3.2% | 41 | 3.2% | 989 | 78.1% | 1267 |
|  | Animal 3 | 240 | 84.8% | 43 | 15.2% | 0 | 0.0% | 0 | 0.0% | 283 |
| **s-mNB**<br>$2 \times 10^4$<br>*eGFP+ engrafted cells* | Animal 1 | 0 | 0.0% | 0 | 0.0% | 0 | 0.0% | 0 | 0.0% | 0 |
|  | Animal 2 | 0 | 0.0% | 0 | 0.0% | 0 | 0.0% | 0 | 0.0% | 0 |
|  | Animal 3 | 0 | 0.0% | 0 | 0.0% | 0 | 0.0% | 0 | 0.0% | 0 |

*IS* = Injection Site
*CC* = Corpus Callosum
*SVZ* = Subventricular zone

Importantly, eGFP^+^ cells were present in the SVZ of all the animals transplanted with eGFP^+^s-iNB and eGFP^+^s-NSC/TAP (Fig. 1Db, Fig. 1Dc), some of them expressing GFAP indicating the generation of astrocytes, and therefore possibly NSC (Fig. 1Db3, Fig. 1Dc2). Plasticity of s-iNB was also shown at the IS by the detection of eGFP^+^ cells expressing markers of astrocytes (GFAP) and oligodendrocytes (CNPase) (Fig.S2B and C).

Overall, these results demonstrate the plasticity of s-iNB *in vitro*, making them very similar to s-NSC/TAP, questioning the exact nature of this abundant SVZ cell populations that express neuroblast markers.

### s-iNB are a molecularly distinct class of proliferating SVZ progenitors

We characterized the transcriptomic profiles of each SVZ populations sorted from 2 month-old C57Bl/6 mice using Clariom^TM^ D mouse microarrays that provides intricate transcriptome-wide gene- and exon-level expression profiles (Fig. 2A).

**Figure 2:**
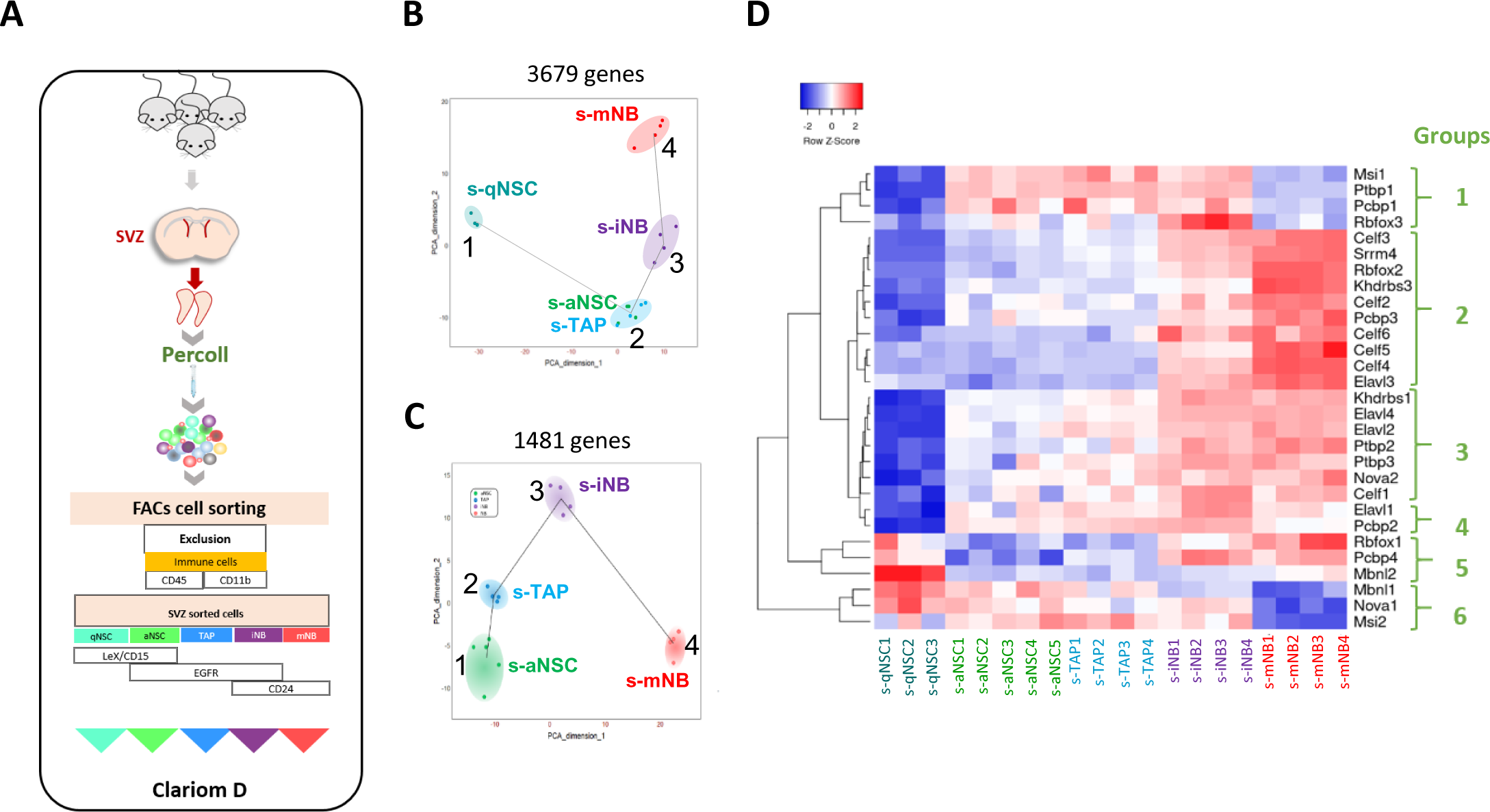
Transcriptomic analysis of SVZ neural progenitor cells. **(A)** Schematic illustration of our strategy of flow cytometry cell sorting of the five major neurogenic populations (s-qNSC, s-aNSC, s-TAP, s-iNB, s-mNB) of SVZ microdissected from adult mouse brains. **(B)** Pseudotime analysis showing the lineage progression of sorted cell populations based on the highly dispersed genes between s-qNSC, s-aNSC, s-TAP, s-iNB and s-mNB. **(C)** Pseudotime analysis after exclusion of s-qNSC based to further discriminate s-aNSC and s-TAP. **(E)** Heatmap representation of RSR genes differentially expressed among s-aNSC, s-TAP, s-iNB, s-mNB (FDR p-value<0.05).

Pseudotime analysis (TSCAN) ^25^ based on 3679 highly dispersed genes between the five sorted cell populations confirmed the lineage progression from s-qNSC (cluster1), then s-aNSC and s-TAP within a single cluster (cluster2), followed by s-iNB (cluster3) and finally s-mNB (cluster4) (Fig. 2B). Another pseudotime analysis excluding s-qNSC, based on 1481 highly dispersed genes, delineated two distinct clusters for s-aNSC and s-TAP (Fig. 2C), highlighting their molecular differences.

Venn Diagram comparison of data sets revealed 2950, 315, 129, 395 and 1693 genes unique to s-qNSC, s-aNSC, s-TAP, s-iNB and s-mNB respectively (Supp_data_1). G:Profiler functional profiling of these transcriptome signatures with KEGG and reactome databases uncovered significant stage-dependent changes (Fig. S3, Supp_data_1). S-aNSC overexpressed genes involved in metabolism of glyoxylate and dicarboxylate metabolism. S-qNSC rather overexpressed genes related to metabolism of lipids and Rho GTPase cycle. S-TAP significantly displayed genes involved in Mitotic, G1 phase and G1/S transition, Activation of the pre-replicative complex. S-mNB specifically overexpressed genes involved in axon guidance and nervous system development. Conversely, s-iNB signatures contained many genes linked to cell cycle, prometaphase, M phase, and cell cycle checkpoints, clearly distinguishing them from the other cell populations including s-mNB.

Alternative splicing (AS) plays various functions in brain development including neural cell differentiation and neuronal migration ^26–30^. Using the splicing index (SI) corresponding to a value assigned to each exon by estimating its abundance with respect to adjacent exons using TAC software (ThermoFisher Scientific), we identified differentially spliced genes (DSG, adjusted p-value ≤ 0.05) unique to s-qNSC, s-aNSC, s-TAP, s-iNB and s-mNB (Supp_data_2). GO annotations of DSG (Fig. S4) clearly revealed that spliced genes in SVZ cells are mainly involved in neuron development and neurogenesis. Interestingly this also showed that qNSC logically differed from the other cell types by splicing concerning genes involved in mitosis and cell cycle, consistently with their quiescent state. More importantly, GO annotations of DSG confirmed that s-TAP and s-iNB have distinct features.

We further analysed particularly DSG at the successive cell transitions along neurogenesis. Results show a very large number of DSG between s-qNSC *vs* s-aNSC (9484) then no DSG at the s-aNSC and s-TAP transition, only 169 between the s-TAP and s-iNB transition, and finally 3759 between s-iNB and s-mNB (Supp_data_3). Functional profiling analysis with g:Profiler linked the DSG between s-TAP and s-iNB to the nervous system development and those between s-iNB *vs* s-mNB to protein binding or cell junction.

We then focused on the differentially expressed genes (adjusted p-value ≤ 0.05) belonging to the nine families of RNA splicing regulators (RSR) that play essential roles in the neurogenic pathway (*Ptbp, Nova, Rbfox, Elavl, Celf, Dbhs, Msi, Pcbp, and Mbnl)* ^26^. Strikingly, a clear down-regulation of most RSR gene expression distinguishes s-qNSCs from the other populations (Supp_data_4). Interestingly another shift discriminates s-aNSC and s-TAP from s-mNB (Supp_data_4). This led us to identify several groups of RSR genes in function of their variation of expression along with neurogenesis: the groups 1 and 6 were overexpressed in s-aNSC and s-TAP and downregulated in s-mNB; the group 2, 3 and 5 were up-regulated in mNB as compared to s-aNSC and s-TAP. All these groups were up-regulated in s-iNB, suggesting that the switch in RSR expression between cycling progenitors and s-mNB occurred in these cells (Fig. 2D). Interestingly, *Ptbp1*, encoding a major splicing repressor ^31^, was downregulated in s-mNB as compared to s-iNB, consistently with the large increase in DSG at the transition between s-iNB and s-mNB.

Altogether, these data confirm at the molecular level that s-iNB comprises cycling progenitors at the transition between s-TAP and s-mNB that undergo important molecular switches, which include major changes in the expression of some RSR genes leading to a major increase in the number of DSG occurring at the final maturation of s-mNB.

### s-iNB correspond to specific SVZ cell clusters identified by single cell RNA sequencing (scRNAseq)

We compared our microarray datasets to published scRNAseq datasets from adult mouse SVZ ^14,17,32^. Violin plots show that s-qNSC and s-mNB perfectly matched with the corresponding populations identified in these studies, which was less obvious for s-aNSC and s-TAP (Fig. S5). s-iNB matched with the clusters termed “Mitosis” (Class VI) in the study of Llorens et al. ^32^ and “Mitotic TAP” in that of Zywitza et al.^17^, and with clusters 8, 10, 17, 16 linked to “Mitosis”, but also with clusters 6 and 15 described as DCX^+^Ki67^+^ neuroblasts in the study by Cebrian_Silla et al.^14^. These comparisons thus confirm that s-iNB comprise cycling SVZ cells exhibiting both TAP and NB features, which illustrates the difficulties to determine the exact nature of cell clusters identified by scRNAseq.

We have previously reported^20^ that after brain exposure to ionizing radiation, proliferating progenitors are highly radiation-sensitive and rapidly eliminated after irradiation. On the contrary qNSC that are radiation resistant repopulate the different SVZ neurogenic cell populations successively the following days after irradiation. We therefore used this model and our transcriptomic signatures of SVZ stem and progenitor cells to gain new insights on the nature and the hierarchical ordering of neural progenitor clusters identified by sc-RNAseq. TenX Genomics droplet-based single-cell transcriptomic analyses were performed on SVZ cells from 2 mice, 5 days after irradiation (4 Gy, SVZ only) and from 2 non-irradiated control mice after removal of myelin and red blood cells (Fig. 3A). Based on a filtering criterion of 1,500 genes/cell to exclude low-quality sequenced cells, 17,343 cells were retained for analysis, divided into 11,529 non-irradiated cells and 5,814 irradiated cells (Fig. S6).

**Figure 3:**
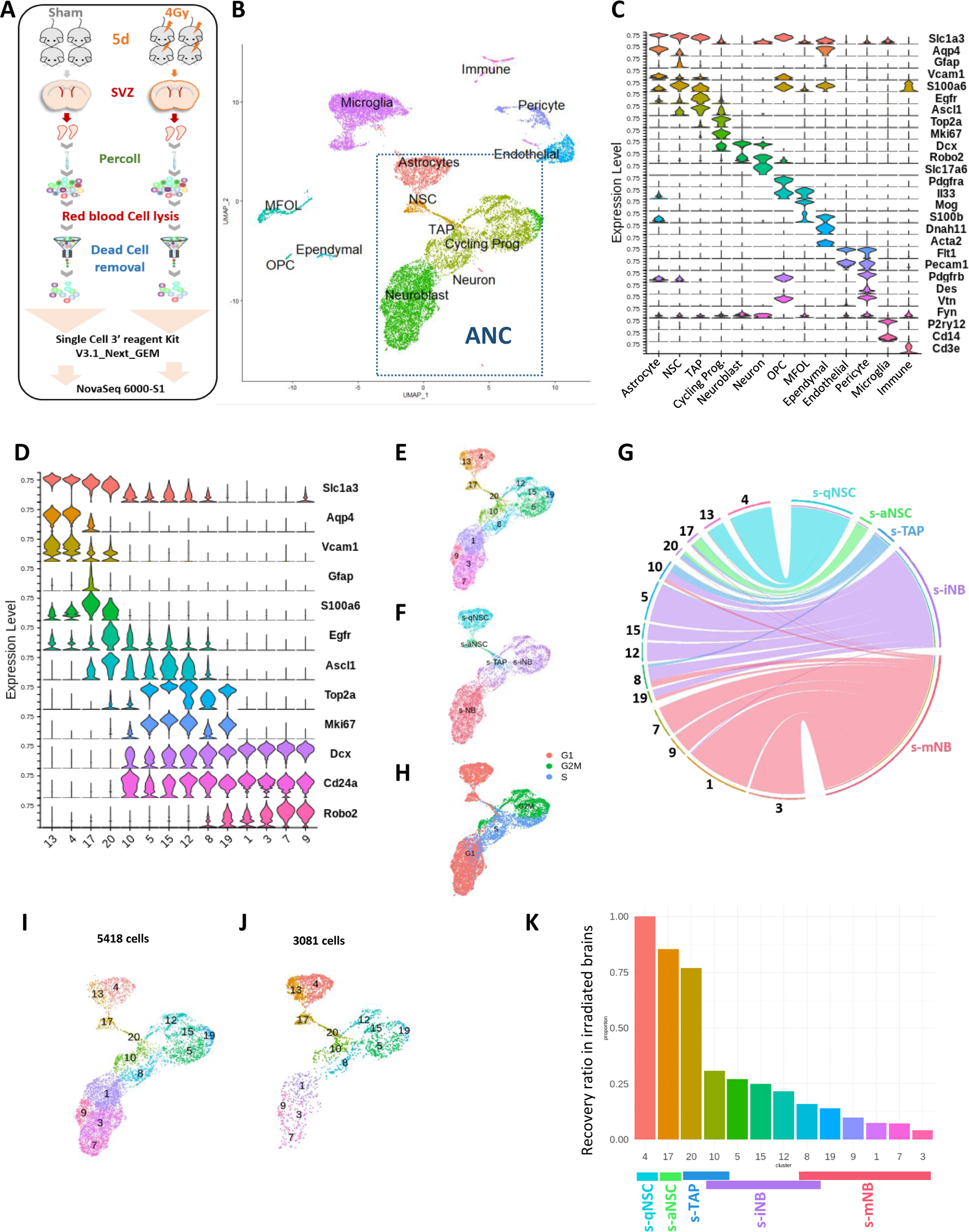
scRNA-seq analysis: whole SVZ. **(A)** Experimental design. **(B)** Uniform Manifold Approximation and Projection (UMAP) of the 17343 high-quality sequenced cells annotated with corresponding cell type, combining scRNA-seq datasets of non-irradiated mice and 4Gy irradiated mice. Clustering analysis at resolution 1.2 segregated cells into 33 clusters that were matched with corresponding cell types determined with marker expression in (**C**). Among them, 14 clusters (dotted lines rectangle) corresponded to the clusters of astrocytes and neurogenic cells (ANC). **(C)** Violin diagrams illustrating the expression level of known cell markers in the 33 clusters according to the literature. **(D)** Violin diagrams representing the expression level of selected markers specific to neurogenic cells and astrocyte from ANC clusters at the resolution 1.2. **(E)** UMAP focusing on the ANC subset. **(F)** UMAP of the top UCell score for the top 100 highly expressed genes of each neurogenic cell sorted population in the ANC clusters (Supp_data_1). **(G)** Chord diagram illustrating the correspondence of cell clusters with cell top score. Cluster 10 matched with both s-TAP and s-iNB and clusters 8 and 19 both to s-iNB and to s-mNB. **(H)** Feature plot of cell cycle scoring **(I, J)** UMAP representation of ANC clusters in the unirradiated (I) and irradiated (J) samples. **(K)** Barplot showing the recovery ratios of each ANC cluster in irradiated brains calculated as the number of cells per clusters from irradiated brains /control brains normalized to the respective numbers of s-qNSC (cluster 4). Taking into account UCell score results, s-iNB clusters overlapped the cluster 10 and 8 containing s-TAP and s-mNB, respectively.

Non-irradiated and irradiated samples were combined and integrated using the SCTransform workflow. After integration, PCA was performed and 50 PCs were used for Uniform-manifold approximation (UMAP). Leiden graph-based clustering, with a resolution of 1.2, identified 33 clusters (Fig. S7A). These clusters were first annotated based on the expression of known cell markers (Fig. 3B and C, Fig. S7B). Nineteen clusters gathering more than half SVZ-residing cells corresponded to non-neurogenic cells (ordered by decreasing abundance): microglia (expressing *P2ry12*^33^), ependymal cells (expressing both *S100β* and *Slc1a3*^34^), endothelial cells (expressing *Flt1*^35^), oligodendrocyte progenitors (OPC, expressing *Pdgfrα*^36^), Myelin forming oligodendrocytes (MFOL, expressing *Mog*^17^) and pericytes (expressing both *Vtn* and *Pcam1* ^7,37^), and dopamine neurons (expressing *Vglut2*^38^). The 14 other clusters corresponded to astrocytes and neurogenic cells (Fig. 3B, C, D). The clusters 4 and 13 that expressed *S100β* were further identified as qNSC (cluster 4) and astrocytes (cluster 13) based on the transcriptomic signatures described by Cebrian-Silla et al., 2021^14^ (Fig S7C, Supp_data_5). Cluster 17, was identified as aNSC based on *Gfap, Thbs4, S100α6*, *Egfr^low^*and *Ascl1* expressions^14^, and cluster 20 as TAP based on *Egfr* and *Ascl1* expressions ^39^. Clusters 10, 5, 15, 12, and 8 were defined as cycling progenitors based on the expression of proliferative markers such as *Top2a*, *Mki67*, *Ascl1.* Clusters 1, 3, 7 and 9 were identified as mNB due to the loss of *Mki67*, *Top2 a* and *Ascl1* expressions and the expression of *Robo2* and *Dcx*. Cluster 19 did not expressed *Ascl1* but *Top2a* and *Mki67* as well as with *Robo2* and *Dcx,* was therefore positioned at the transition between iNB and mNB.

We then applied the UCell workflow to further characterize the neurogenic clusters with the transcriptomic signatures of each of our five sorted SVZ cell populations (Supp_data_1), calculating individual cell scores for each signature based on the Mann– Whitney U statistical analysis, and attributing cells to populations based on the highest score amongst signatures^40^. This score allowed us to unequivocally attribute some clusters to the sorted cell populations (Fig. 3E,F and G): cluster 4 and 13 matched with s-qNSC, cluster 17 with s-aNSC, cluster 20 with s-TAP, clusters 5, 15 and 12 with s-iNB and finally clusters 7, 9, 1 and 3 to s-mNB. Reflecting the continuous nature of the SVZ neurogenic process, the cluster 10 matched both with s-TAP and s-iNB and clusters 8 and 19 both to s-iNB and to s-mNB. Cell-Cycle Scoring^36^ indicated that the clusters related to s-iNB were enriched in cells in S or G/2M phase (Fig. 3H).

As the consequence of radiation-induced cell death, the relative abundances of each cluster were different in controls (Fig. 3I) and irradiated (Fig. 3J) brains and dependent on the progressive repopulation of the different types of progenitors. Normalized to the numbers of s-qNSC (cluster 4), which are supposed not to be affected by irradiation^20^, the ratios of the number of cells per clusters in irradiated brains over that in unirradiated controls gave an index of the progressive reconstitution of each cluster (Fig. 3K). As expected, clusters 17 and 20, corresponding to s-aNSC and s-TAP respectively, were almost entirely regenerated. The decreasing values of the ratios evidence a temporal ordering of the regeneration of each cluster corresponding to s-iNB along with neurogenesis as follows: 10, 5, 15, 12, and 8, the latter being the one at the iNB vs mNB transition as anticipated above.

We next addressed the expression of RSR genes in each cluster (Fig. 4). Not all genes could have been investigated since some of them were below the detection threshold in all the cell populations studied. The expression of RSR genes *(Kdrbs3, Kdrbs1 Elavl3 and Elavl2, Elavl4, and Celf1*) that were shown to increase along with neurogenesis in microarrays, also increased in cluster 10, remained stable in the other clusters related to s-iNB and reached a maximum value in the s-mNB clusters (Fig. 4A). Conversely, the expressions of *Mbnl1* and *Msi2* that were shown to decrease along with neurogenesis in microarrays (Fig. 4B) peaked in the clusters related to s-aNSC (17) and s-TAP (20), decreased in the cluster 10, remained low in the clusters related to s-iNB (5, 15, 12, 8) and were not detected in the clusters related to s-mNB (19, 1, 3, 7, 9). Altogether, these data corroborate our microarray data and show that a major switch in the expression of RSR genes occurs in the clusters related to s-iNB.

**Figure 4:**
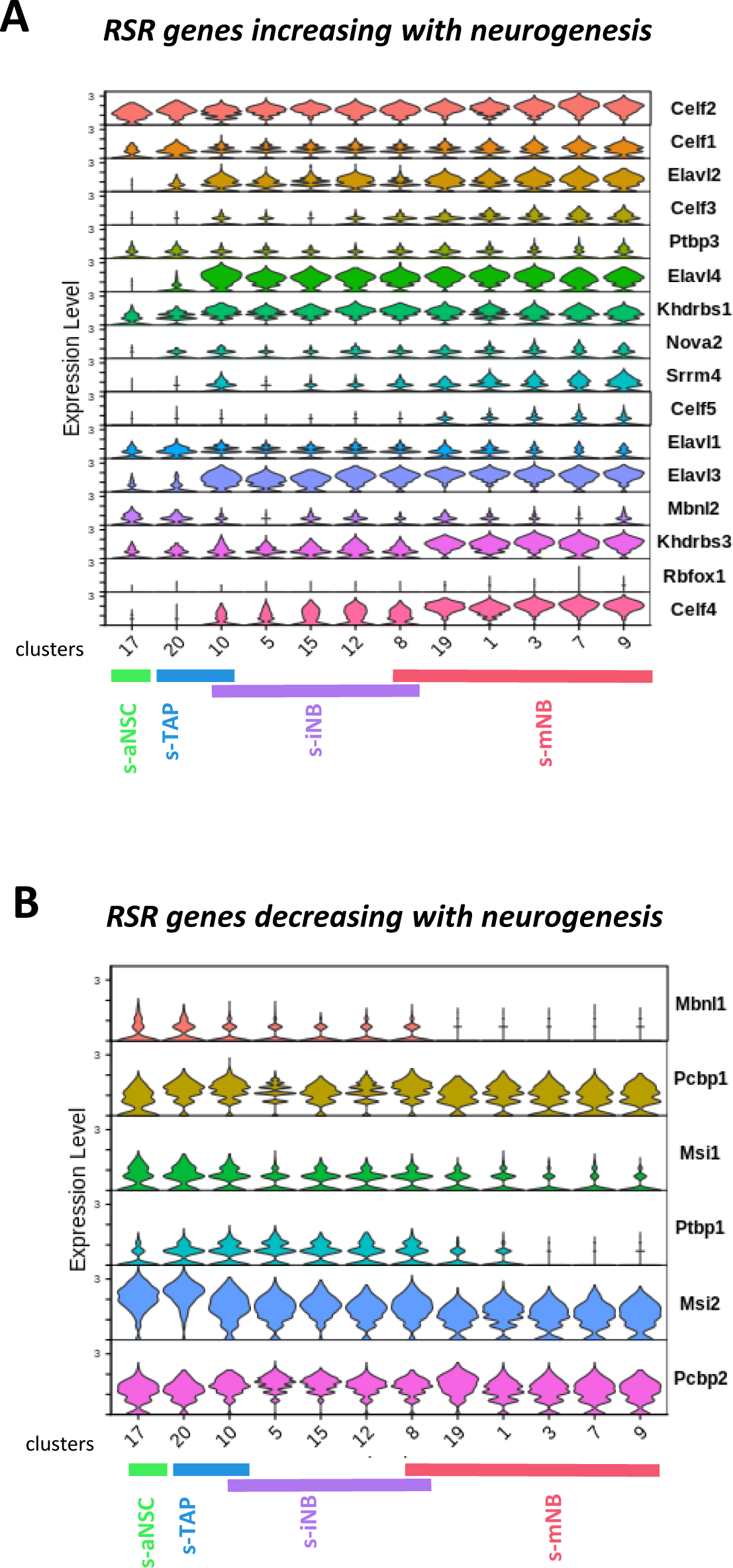
Expression of RSR genes in scRNA-seq neurogenic clusters. Violin diagrams illustrating the expression of the RSR genes increasing **(A)** or decreasing **(B)** with neurogenesis.

Interestingly, the splicing repressor *Ptbp1* was similarly expressed in s-aNSC, s-TAP and s-iNB-related clusters but was downregulated progressively in the successive s-mNB clusters, suggesting that changes in *Ptbp1* expression constitute an important molecular step for the final maturation of mNB, which involves a major increase of DSG.

### The *Dcx* status identifies two molecularly distinct subpopulations in s-iNB

We have previously shown by immunofluorescence that not all the s-iNB were positive for doublecortin (*Dcx*) expression, a common marker of neuroblasts^20^. Violin plot (Fig. 3D) and feature plot (Fig. 5A) show that *Dcx* expression was restrained to s-mNB and to clusters related to s-iNB in which it increased consistently with their hierarchical ordering we defined above. Variations in *Dcx* expression among iNB related clusters led us to analyse separately *Dcx^low^* and *Dcx^high^* cells in these clusters (Fig. S8). The ratios of numbers of cells per cluster indicated a greater recovery of *Dcx^low^* cells as compared to *Dcx^high^*cells in s-iNB related clusters from irradiated brains (Fig. 5B), reflecting their temporal ordering.

**Figure 5:**
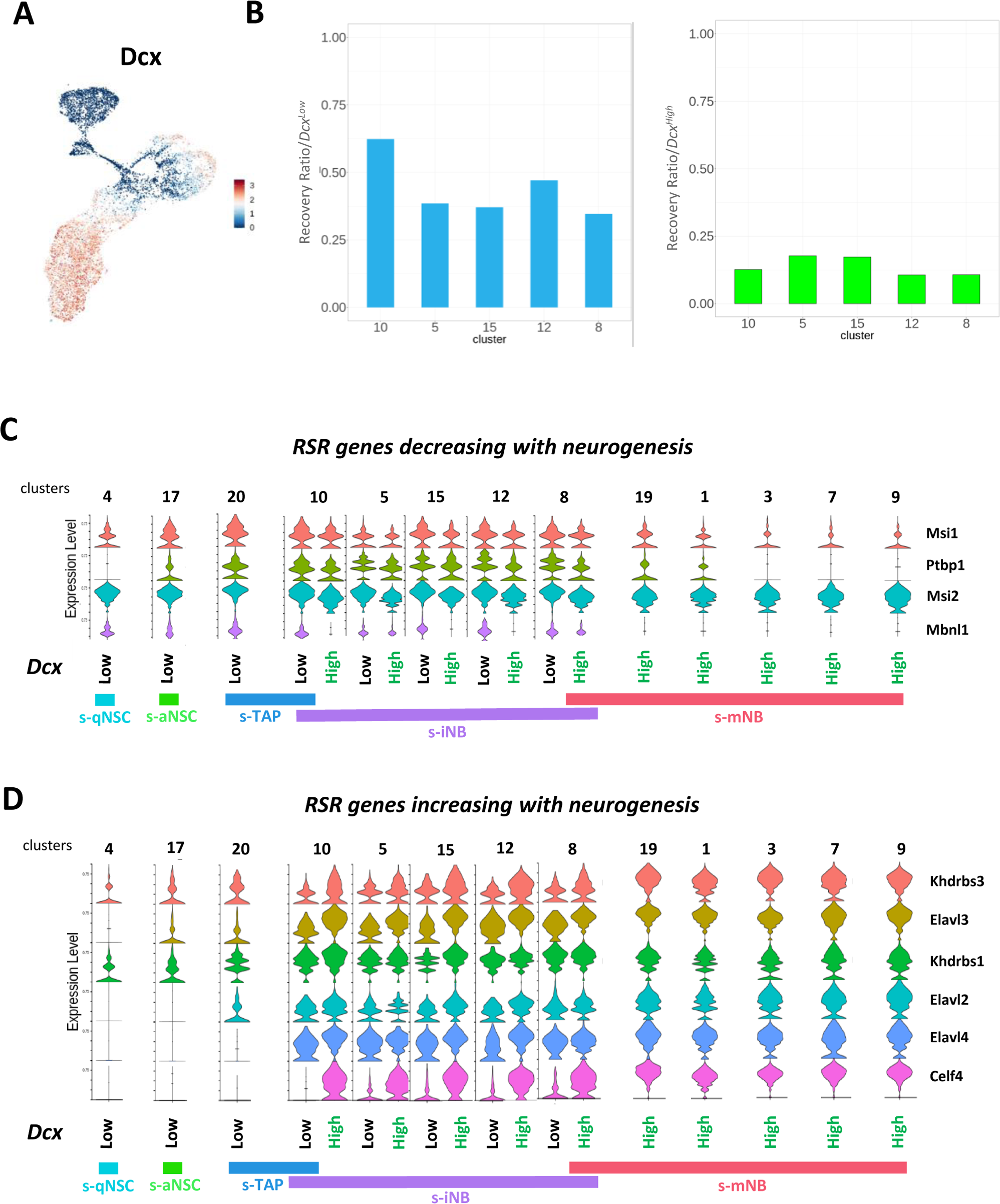
*Dcx* expression in the neurogenic cell clusters. **(A)** Feature plot. **(B)** Barplot representations of the recovery ratios of *Dcx^low^* (left) and *Dcx^High^* (right) in the iNB clusters of the irradiated brains, calculated as in Figure 3K. Violin diagrams illustrating the expression level of RSR genes decreasing **(C)** and increasing **(D)** with neurogenesis in *Dcx^low^* and *Dcx^High^* in neurogenic cell clusters.

Characterization of differentially expressed genes according to the *Dcx* status in these clusters showed 588 unique genes expressed in the *Dcx^low^* cells and 98 genes specifically expressed in the *Dcx^high^* cells. Functional enrichment profiling using the g:Profiler software illustrates biological differences between both populations : genes related to Metabolism of RNA were overrepresented in the *Dcx^low^* cells whereas genes linked to GABAergic synapse, Neuronal System were rather overexpressed in the *Dcx^high^* cells (Supp_data_6). Notably, 55% of *Dcx^high^* cell specific genes were s-mNB genes (*e.g. Robo2, Gap43, Nav3*). Further analysis of RSR genes expression in function of the *Dcx* status, clearly confirms that progressive transcriptomic changes occurred between *Dcx^low^* and *Dcx^high^* cells (Fig. 5C and D). However, RSR gene profiles of *Dcx^low^* cells in clusters related to s-iNB were obviously different from that related to both s-TAP and s-NSC, and that of *Dcx^high^* cells from that of s-mNB. Similar results were obtained with cells from irradiated brains (Fig. S9).

These data clearly show that the clusters related to s-iNB contained cells engaged in neuronal differentiation that are molecularly distinct from the other types of SVZ neurogenic cells already characterized This suggest that s-iNB correspond to a new stage of cycling neurogenic progenitors progressively engaged in neuronal differentiation.

We next took advantage of the DCX-CreERT2::CAG-floxed-eGFP mice model, allowing the expression of eGFP in cells expressing *Dcx* after the administration of Tamoxifen to sort eGFP^+^CD24^+^EGFR^+^ cells corresponding to *Dcx^high^* iNB and eGFP^−^ CD24^+^EGFR^+^ cells containing the *Dcx^low^* iNB to distinguish further these two populations at the functional level. As shown in Fig. 6A and B, recombinant eGFP^+^ iNB exhibited a clonogenic potential and a rate of population doublings quite similar to eGFP^−^ iNB and total iNB, contrary to eGFP^+^ mNB, which did not show clonogenic potential and did not proliferate at long-term *in vitro*. Moreover, we showed that both eGFP^+^ iNB and eGFP^−^ iNB have the capacity to generate the three neural lineages, neurons, astrocytes and oligodendrocytes, when plated for 5–7 day in the appropriate differentiation media (Fig. 6C), similarly as s-iNB (Fig. 1C).

**Figure 6:**
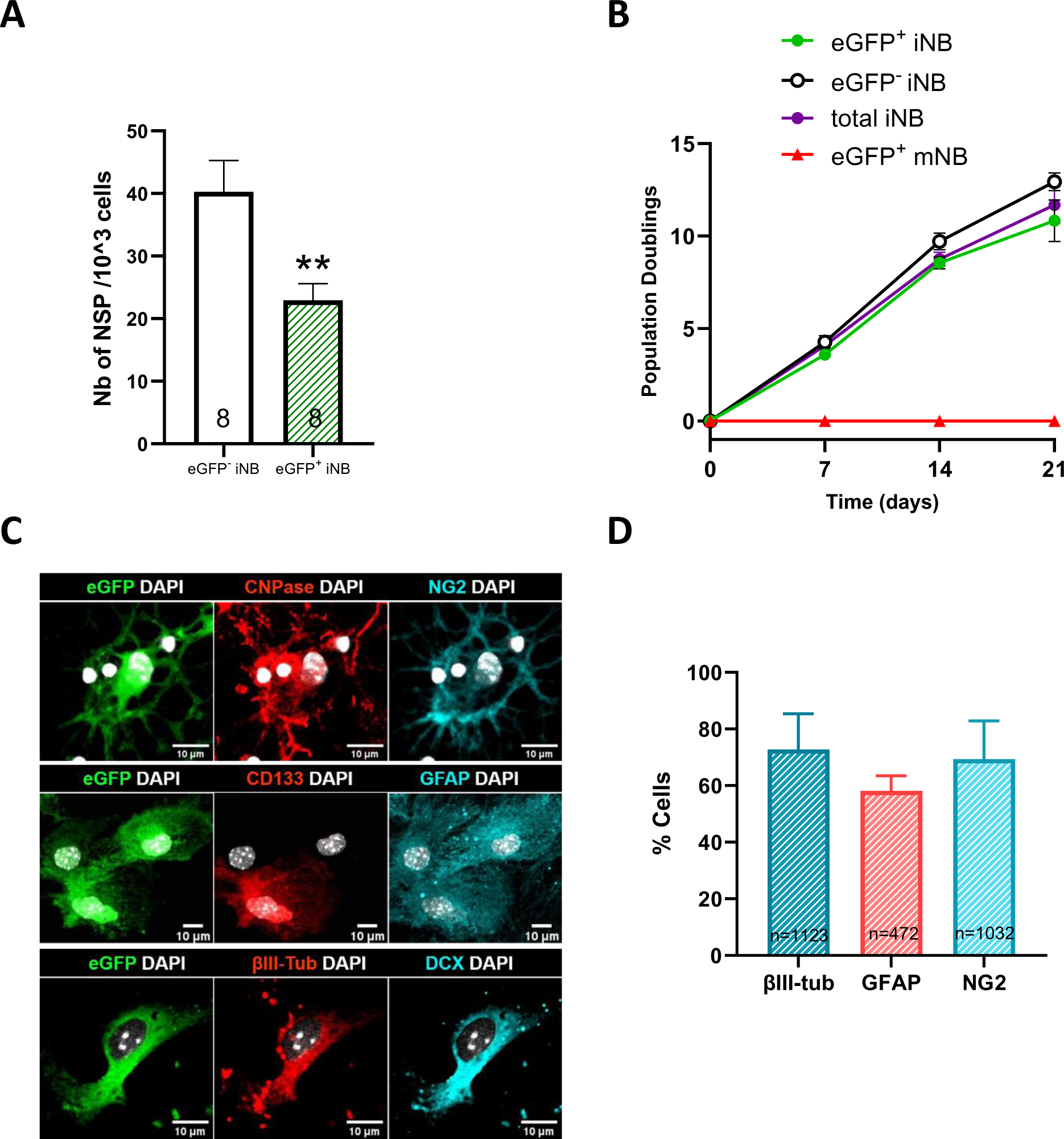
Plasticity *of Dcx^High^* iNB *in vitro*. **(A)** Clonogenic potential of FACS-isolated eGFP^+^iNB and eGFP^−^iNB, and recombined eGFP^+^mNB isolated from DCX-CreERT2::CAG-floxed-eGFP mice, 7 days after the first injection of tamoxifen. **(B)** Population doublings (PD) of FACS-isolated eGFP^+^iNB and eGFP^−^iNB, eGFP^+^mNB cells and total iNB cells (regardless of eGFP status). No statistical difference was found in the growth rate of the different isolated populations. (n = 8) **(C)** Representative images of immunofluorescence of freshly sorted eGFP^+^iNB cells cultured in oligodendrocyte,astrocyte, or neuronal differentiation medium, and stained for NG2 and CNPase, GFAP and CD133, βIII-Tubulin and DCX, respectively. Scale: 10µm. **(D)** Quantification of the percentage of eGFP^+^ iNB among DAPI cells cultured in differentiation media that differentiate in either neurons (βIII-tubulin), oligodendrocytes (NG2) or astrocytes (GFAP).

By contrast, when eGFP^+^iNB were transplanted into the brain of C57Bl/6J mice, no eGFP^+^ cells persisted at long-term (5 weeks) (Fig. S10), suggesting that the *in vivo* plasticity and regenerative properties of s-iNB was rather associated to *Dcx^low^* iNB.

## Discussion

This study bridges the cellular and molecular characterizations of SVZ neural progenitor populations. We combined the transcriptional profiling using DNA microarrays of SVZ that were sorted by an efficient FACS strategy^19^ and large-scale scRNAseq to decipher the progressive molecular and cellular changes involved in SVZ neurogenesis. Our model of regeneration of the neurogenic niches after brain irradiation^20^ allowed a unique way to determinate the sequential regeneration of the different types of neurogenic cell clusters identified by scRNAseq that were precisely characterized using the respective molecular signatures of sorted-SVZ cells, providing new insights onto the progression of SVZ neurogenesis. We show that iNB is an abundant population of cycling progenitors, which is more advanced towards neuronal differentiation than TAP, while retaining unexpected stem cell properties unlike mNB. We suggest that major splicing regulations in iNB might be critical for the final commitment to the neuronal fate.

Previous studies of the different sub-populations of SVZ progenitors were carried out using transcriptomic approaches based on the expression of various more or less specific markers. These approaches enabled the identification of quiescent and activated neural stem cells as well as mature neuroblasts, but were confronted with the strong influence of the cell cycle on cell clustering. Indeed, neural progenitors in these studies cycling have been gathered in either “mitotic” clusters ^14,17,32^ or “neural progenitor cells” clusters ^15^ that lacked clear biological significance and prevented the identification of subtypes of SVZ cycling progenitors. Our study, combining for the first time characterization of FAC-sorted cells and an irradiation-based model of sequential regeneration, clearly distinguished the molecular profiles of TAP and iNB among cycling progenitors reflecting differences in their respective potentials *in vitro* and *in vivo*.

Our results constitute thus a valuable resource for delineating the molecular transitions occurring during neurogenesis that are orchestrated by numerous differentially expressed genes and alternative splicing variants. The transition from qNSC to aNSC has been abundantly investigated in the literature^15,32^, but the transcriptional shift leading to the differentiation of neural progenitors into neuroblasts has not been thoroughly analysed to date. We showed that it occurs in immature neuroblasts, which form a relatively abundant subpopulation of SVZ cells, which exhibit specific transcriptional features distinguishing them from TAP and mNB.

We formerly described the isolation of s-iNB as a subset of cycling SVZ cells that expressed both CD24, a neuroblast marker, and EGFR, a marker of cycling progenitors^19–21^. This was consistent with prior findings describing a subset of neuroblasts co-expressing Ascl1 and Dcx or CD24^low^ expressing low levels of EGFR in the adult SVZ^13,39,43^. Here, we evidenced that s-iNB comprise a molecularly distinct population of cycling cells at the transition between TAP and mNB, and exhibiting molecular characteristics of both these cell types.

The capacity of neuroblasts to reorient towards the glial lineage in pathological contexts have been reported previously^44–46^. Consistent with the molecular findings showing that s-iNB kept transcriptional features of NSC and TAP, we showed that s-iNB also kept *in vitro* and *in vivo* a regenerative potential similar to that of s-aNSC and s-TAP. Importantly, this may also include the capacity to dedifferentiate *in vivo* and generate NSC as suggested by the detection of GFAP-positive engrafted cells persisting in the SVZ.

Interestingly, s-iNB corresponded to several scRNAseq clusters, which were not clearly characterized in the literature. Our model of SVZ regeneration established the hierarchical ordering of these clusters, in which we showed progressive molecular changes leading to the differentiation into migrating neuroblasts. This includes the increase in *Dcx* expression. *Dcx*^high^ cells in these clusters have transcriptional features confirming they are more engaged in the differentiation than their *Dcx*^low^ counterparts. Importantly, we showed that *Dcx*^high^ and *Dcx*^low^ cells in these clusters remained molecularly distinct from both TAP and mNB. We thus propose a revision of the current model of SVZ neurogenesis by introducing iNB cells, as highly cycling progenitors that undergo progressive neuronal differentiation but kept multipotency (Fig. 7). iNB cells could be further divided in two subclasses of cells depending on their progression into the neuronal differentiation process: iNB1 cells, corresponding to *Dcx*^low^ iNB, immediately succeeding the TAP and iNB2 cells, corresponding to *Dcx*^high^ iNB and preceding the stage mNB. Importantly, we showed that both iNB1 and iNB2 cells kept transcriptomic features that differentiated them from both TAP and mNB, whereas iNB1 cells kept a regenerative potential both *in vitro* and *in vivo*.

**Figure 7:**
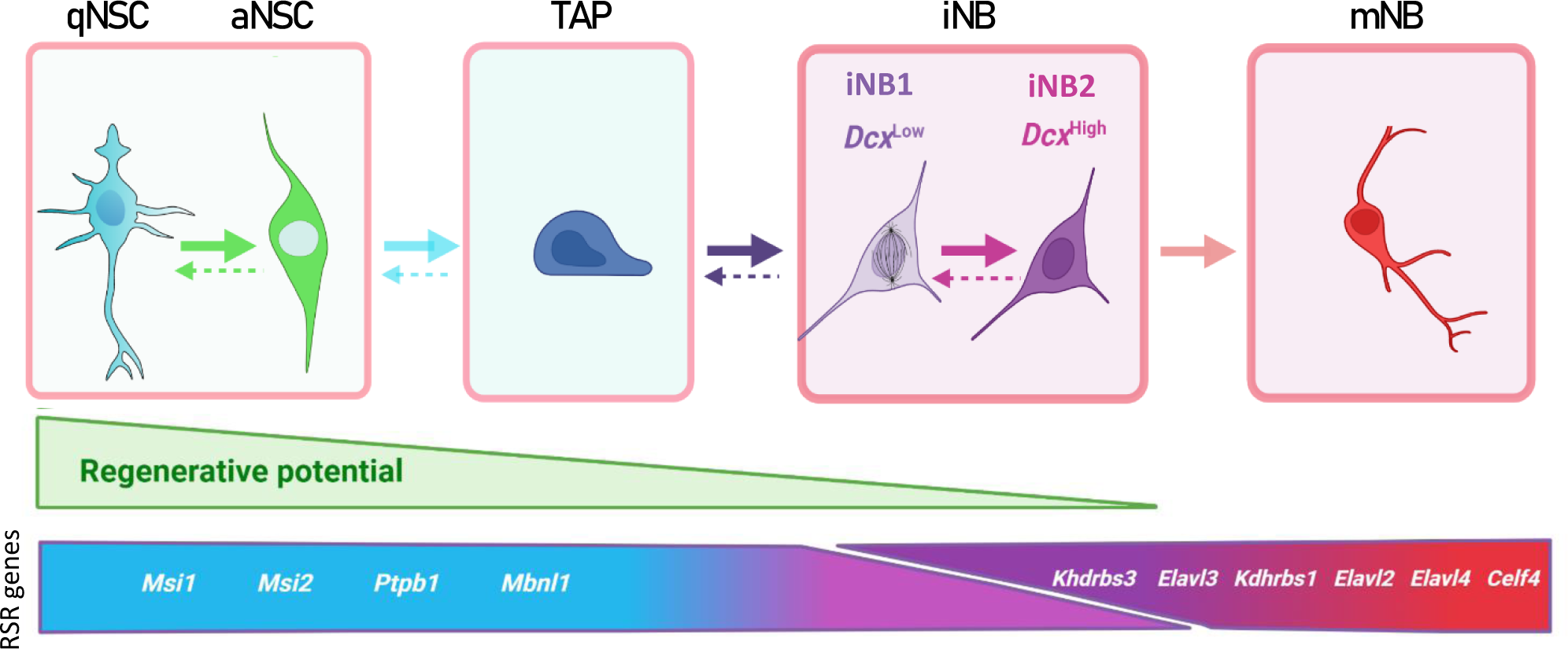
Proposed model of adult mouse SVZ neurogenesis involving major changes in RSR genes expression. Possible interconversion of iNB states would require further experimental confirmation.

One of the most important findings of this study is the first characterization of AS events and RSR expression variations occurring along with neurogenesis. AS events are known to play important roles in neural cell differentiation, neuronal migration, synapse formation, and brain development^27,30,47–49^. Neurons are characterized by a pervasive use of alternative exons^50,51^. Consistently, we showed a dramatic increase in DSG at the transition between s-iNB and s-mNB. These alternative splicing events are clearly related to major changes in RNA splicing regulators (RSR) expressions occurring in s-iNB. Indeed, we identified RSR genes whose expression increased with neurogenesis and others whose expression decreased with neurogenesis. Consistently, some genes upregulated in s-aNSC, s-TAP, such as *Msi1* and *Msi2,* have a known role in maintenance and renewal of stem cells^52–54^, while some others upregulated in s-mNB, such as *Elavl3 and 4,* have known roles in many neuronal processes^54,55^. Interestingly, genes of both groups of genes were overexpressed in scRNAseq clusters related to s-iNB, which thus undoubtedly distinguished these cells from both s-TAP and s-mNB and confirmed that important molecular switches for neuronal differentiation occurred in D cells.

Further analysis of scRNAseq clusters showed that these transcriptomic switches occurred sequentially from iNB1 cells to iNB2 cells. Indeed, the expression of *Elavl4* progressively increased from the *Dcx^low^*iNB1 cells to the *Dcx^high^* iNB2 cells fractions, confirming their progressive commitment into neuroblasts, and overall, the interest of discriminating these two cell types in the model of SVZ neurogenesis we propose (Fig.7). Strikingly, the splicing repressor, *Ptbp1*, was expressed in iNB1 cells at levels similar to those in s-aNSC and s-TAP, but decreased in iNB2 cells, and was undetectable in migrating neuroblasts. This suggests that expression of *Ptbp1* may play an important role in maintaining the regenerative properties in iNB1 cells, which should be further investigated.

Our study has been performed on cells collected from the SVZ the lateral walls of the ventricles of young adult mice. The transcriptional signatures we provide here may thus be useful for further characterization of neurogenic cells present in the septal wall of the SVZ^18^.

To conclude, we have characterized the molecular features of iNB cells, a relatively abundant population in the adult SVZ that kept unexpected plasticity and multipotency that persist in the aged brain^21^. They may thus represent a new target in regenerative medicine. We have shown that these cells undergo major transcriptomic changes involving AS and associated to switches in RSR genes expression. Dysregulations of many of these RSR genes have been involved in cancer and neurodegenerative diseases^56^. Further characterization of iNB cells using the tools we developed may be useful to determine the potential therapeutic targeting of these cells in various brain pathologies.

## Supporting information

script_R_6

script_R_1

script_R_2

script_R_3

script_R_4

Script_R_5

## Acknowledgments

We thank C. Devanand and C. Duwat for their contribution for animal experiments and G. Piton for 10x single cell library preparations. We are grateful to Marie-Justine Guerquin and Pierre Fouchet, and the ARTBIO platform (Université Paris 6) for helpful discussions and assistance in bioinformatics analyses.

B.C has a fellowship from ARSEP (fondation pour l’Aide à la Recherche sur la Sclérose En Plaques). This work was supported by grants of IRBIO (Commissariat à l’Energie Atomique et aux Energies Alternatives, CEA) and Electricité de France (EDF). The funders have no role in the study design, data collection and analysis, decision to publish, or preparation of the manuscript.

## Author contributions

<u>Corentin Bernou:</u> Collection and assembly of data, data analysis and interpretation, manuscript writing.

<u>Marc-André Mouthon:</u> Collection and/or assembly of data.

<u>Mathieu Daynac:</u> Collection and/or assembly of data.

<u>Thierry Kortulewski:</u> Collection and/or assembly of data.

<u>Benjamin Demaille:</u> Analysis of Clariom datasets.

<u>Vilma Barroca:</u> Intracerebral neural progenitor grafting in mouse models.

<u>Sébastien Couillard-Despres:</u> for DCX-CreERT2:CAG-CAT-eGFP mice.

<u>Nathalie Déchamps:</u> Cell sorting by flow cytometry.

<u>Véronique Ménard:</u> Irradiation of animals.

<u>Léa Bellenger:</u> Analysis of scRNAseq datasets.

<u>Christophe Antoniewski:</u> Analysis of scRNAseq datasets.

<u>Alexandra Chicheportiche:</u> Conception and design, collection and assembly of data, data analysis and interpretation, manuscript writing.

<u>François D. Boussin:</u> Supervision of the study, conception and design, financial support, data analysis and interpretation, manuscript writing.

## STAR Methods

### KEY RESSOURCES TABLE

### EXPERIMENTAL MODEL AND SUBJECT DETAILS

#### Mice

All experimental procedures complied with European Directive 2010/63/EU and the French regulations. The protocols were approved by our institutional ethical committee (CEtEA 44) and authorized by the « Research, Innovation and Education Ministry » under registration number APAFIS#25978-2020061110358856 v1. Adult male C57BL/6J were obtained from Janvier Laboratories. Adult male β−actin:eGFP (Okabe et al., 1997) and DCX-CreERT2::CAG-CAT-eGFP mice kindly given by Pr Couillard-Despres Sebastien ^57^. Experiments were performed with male animals aged 8 and 10 weeks. Tamoxifen (TAM, T-5648, Sigma-Aldrich) was dissolved in corn oil (C-8267, Sigma-Aldrich) at 40 mg/ml. To analyse the expression of the *Dcx* reporter gene, 20 mg TAM/g body weight was administered twice daily by gavage over a period of five consecutive days.

#### Cell sorting

Lateral ventricle walls of the SVZ were microdissected and dissociated as previously described ^19–21^ with the modification that cells were centrifuged at 250g for 20 min at 4°C without brake on a 22% Percoll gradient (GE Healthcare) to remove myelin prior to single cell suspension labelling. The antibodies to distinct cell populations were anti-CD24 phycoerythrin [PE]-conjugated (Rat IGg2b; BD Biosciences, 1:50, RRID:AB_2034001), CD15/LeX flurorescein isothiocyanate-[FITC] conjugated (clone MMA, mouse IgM; BD Biosciences, 1:50, RRID:AB_400103), CD15/LeX-Brilliant violet-421-[BV421] (Mouse IgG1, κ; BioLegend, 1:50, RRID:AB_2566519) and Alexa647-conjugated epidermal growth factor (EGF) ligand (Clone 30H45L48, Thermo Fisher Scientific, 1:200, RRID:AB_2662334). Cells were gated following the fluorescence minus one (FMO) control. Immediately prior to FACs sorting, propidium iodide (PI) or Hoechst 33258 was added to a final concentration of 1µg/ml to label the dead cells. Cells were sorted on an INFLUX cell sorter equipped with an 86 µm nozzle at 40 psi (BD Biosciences). All the data were analysed using FlowJo software (Tree Star, Ashland, OR).

#### Cell culture expansion and differentiation

Clonogenicity assay and population doublings: FACs-purified populations were collected in NeuroCult medium complemented with the proliferation supplement (STEMCELL Technologies), 2 µg/mL heparin, 20 ng/mL EGF and 10 ng/mL FGF-2, at a density of 1×10^3^ cells/well in 96-well tissue culture plates coated with poly-D-lysine (Merck Millipore). Five days after plating, the neurospheres were counted to determine the clonogenicity of each cell population.

Neurospheres were then mechanically dissociated and sub-cultured in 24-well plates over 3 weeks to measure the population doublings at each passage (1 passage/week).

Differentiation assays: just after sorting, cells were placed on Poly-L-ornithine (P4957, Merck Millipore) coated 24-well culture plates at 40,000 cells/ml in DMEM:F12 with several supplementation according to the lineage: astroglial : 2% B27 MAO (minus antioxidant, Thermo Fisher Scientific) and 2% fetal bovine serum (10082139, Thermo Fisher Scientific), oligodendroglial : 10 ng/mL FGF-2, a defined hormone mix including Glucose (G-7021, Sigma-Aldrich 50gm/l) /NaHCO_3_ (S-5761, Sigma-Aldrich, 10mg/l) / HEPES (H-3784, Sigma-Aldrich, 1.3mM), insulin (I-1882, Sigma-Aldrich, 22.5mg/l), Apo transferrin (T-2252, Sigma-Aldrich, 0.09g/l), progesterone (P-6149, Sigma-Aldrich, 18µM), putrescine (P-7505, Sigma-Aldrich, 8.7mg/l), and sodium selenite (S-9133, Sigma-Aldrich, 4.7µM), and neuronal : 2% B27 MAO. Cultures in hypoxic conditions (4%O2) at 37°C for 5 (oligodendroglial or astrocytic differentiations) or 7 days (neuronal differentiation).

#### Transplants

Three injections of 1µl of freshly eGFP^+^ cells from β−actin:eGFP or DCX-CreERT2:CAG-CAT-eGFP mice were performed in the striatum at proximity of SVZ of wild-type C57Bl6/J recipient mice using the following coordinates: (1) AP, 0.0; L, 1.4; V, −2.1; (2) AP, 0.5; L,1.1; V, −2.2; (3) AP, 1.0; L, 1.0; V, −2.5mm relative to bregma, as described before by Codega et al. 2014^2^. For CD1 mouse transplants with freshly eGFP^+^ cells from DCX-CreERT2:CAG-CAT-eGFP mice the injection coordinates were redefined from the ratio of the distance bregma to lambda for the C57Bl6/J model over the distance bregma to lambda for the CD1 model.

The transplantations were performed using a small animal stereotaxic apparatus (Kopf model 900) with a 2.5µl Hamilton syringe (Hamilton, Bonaduz, Switzerland). Recipient mice were sacrificed 5 weeks after transplantation.

#### Tissue processing, immunolabeling and microscopy

Deeply anaesthetized animals received a transcardial perfusion of 4% paraformaldehyde. Brains were post-fixed overnight in 4% PFA and cryoprotected in 30% sucrose/PBS. Serial coronal cryostat sections were made at 14µm of thickness (Leica CM3050S). For each brain, all the sections from the hippocampus to olfactive bulb were deposited on a set of 14 slides with a 150µm step. Sections underwent permeabilisation in phosphate buffered saline (PBS) containing 0.3% Triton X100 and 1% of Bovine Serum Albumin 1h at room temperature. Sections were incubated with primary antibodies PBS containing 0.1% Triton X100 overnight at 4°C. Secondary antibodies used were Alexa 594, 488 and 647 and were applied at 1:400 (Thermo Fisher Scientific) for 2h at room temperature. Each staining was replicated in at least 3 slides from different mice.

Cells were fixed with 4% PFA for 15 minutes at room temperature and rinsed three times with PBS. Staining was performed as above except that antibodies were applied for 2h at room temperature.

The following antibodies used were: rabbit anti-eGFP (Abcam, ab290, 1:300), goat anti-GFP (Abcam, ab6673, 1:300), mouse anti-O4 (Millipore, MAB345, 1:100), rabbit anti-NG2 (Millipore, AB5320, 1:100), goat anti-Olig2 (RαD, AF2418, 1:200), mouse anti-NeuN (Millipore, MAB377, 1:100), rabbit anti-Dcx (Cell Signaling, 4604, 1:400), goat anti-Dcx (Santa Cruz, sc-8066, 1:100), mouse anti-GFAP (Millipore, MAB3402, 1:400), rabbit anti-Gfap (Sigma-Aldrich, G9269, 1:400), mouse anti-CNPase (Millipore, MAB326, 1:100), rabbit anti-β-III tubulin (Covance, PRB-435P, 1:200), rat anti-CD133 (Millipore, MAB4310, 1:100).

Brightfield and fluorescent images were captured through Plan Apo X20/ Numerical Aperture (NA):1.3 oil objective and Plan Apo X40/ NA:0.75 dry objective using hybrid detection technology on a laser scanning confocal (Leica Microsystems SP8), Nikon A1R confocal laser scanning microscope system attached to an inverted ECLIPSE Ti (Nikon Corp., Tokyo, Japan)

#### SVZ irradiation

Animals were anesthetized with isoflurane (3% induction for 5 minutes) and 1.5-2% when the mice were placed in Small Animal Radiation Research Platform (SARRP, XSTRAHL, LTD Company). First, a Cone Beam Computer Scanning (CBCT) was performed to target the SVZ and to set up the treatment planning system. Mice were placed on their stomach and two fields (−45° and + 45°) were used to be homogeneous on the irradiation target. The size of the irradiation field was adapted with a multivariable collimator 10*6 mm. The SVZ received a 4Gy dose distributed 50% by each beam and the dose output was around 3.64 Gy/min depending of the size of the field. The X-ray configuration was 220 kV, 13 mA and 0.15 mm of Copper in these conditions.

#### Single cell library preparation for single cell RNA sequencing

SVZ lateral ventricle walls were dissected from two non-irradiated mice brains (2 month old) and two 5 days post-irradiation 4Gy-irradiated mice brains (2-month old).

Protocol of SVZ digestion with Papain (Worthington, LK003150) to obtain single cell suspensions were strictly identical to those performed in the whole genome microarray experiment described above, including the Percoll gradient step to remove myelin. Cells were resuspended in 0.04% BSA in PBS passed through a 70µm cell strainer (ThermoFisher Scientific).

Blood cells were lysed using the Red blood Cell lysis solution (Milteny Biotec, 130-094-183) and dead cell Removal using the Dead Cell Removal Kit (Miteny Biotec, 130-090-101) were performed following the manufacter’s instructions.

Single cell suspensions of two replicates of each sample (unirradiated, 4Gy) were adjusted to 2000 cells/µl and 10,000 non-irradiated cells and 6,000 irradiated cells were loaded per channel onto Chromium Next GEM Chip Single Cell kit (ref: 1000127). Library preparation was performed using the Dual Index Kit TT (ref: 1000215) according to manufacturer’s recommendations (10x Genomics, Pleasanton, CA). Quality and quantification of libraries were done using the High Sensitivity DNA LabChip kit (ref: 5067-4626). Sequencing NGS Standard Illumina was carried out with the NovaSeq6000-S1 flow cell (ICM platform, Paris).

### METHOD DETAILS

#### Whole-genome Microarrays

mRNA were isolated with the RNeasy Micro Kit with DNase treatment (Qiagen) and sample quality was controlled using the Agilent Bioanalyser. MTA-1.0 arrays (Clariom D Affymetrix technology) were performed according to the manufacturer’s protocol (ThermoFisher). PCA and pseudotime ordering was performed using TSCAN online user interface (Ji and Ji, 2016).The data were analysed with the freeware software Transcriptome Analysis Console (TAC) applying the RMA algorithm normalization. Functional profiling were generated using REACTOME and KEGG datasets obtained from the g:Profiler interface (https://biit.cs.ut.ee/gprofiler/gost).

Signatures and unique mRNA splicing isorforms of cell sorted populations were delineated using Venn diagrams comparing the five populations of neural progenitors (s-qNSC, s-aNSC, s-TAP, s-iNB and s-mNB) (https://bioinformatics.psb.ugent.be/cgi-bin/liste/Venn/calculate_venn.htpl).

##### sc-RNAseq and microarray dataset comparisons

Published genelists were matched with microarray data raw data using aggregated Z-scores. For each gene in a genelist, gene expression amongst all samples was normalized between 0 and 1, then mean expression among samples was calculated for each population. Plots represent the distribution of mean normalized gene expression for all genes in the genelist.

#### Single-cell RNA control quality and analysis

ScRNA-seq data were aligned to the GRCm38 - mm10 reference genome (cellranger-6.1.2). Data were analysed in R (version 4.1.0) using Seurat (v4.1.1). Quality control removed outlier cells with fewer than 1200 to 1500 genes per cell (nFeatures_RNA), 2350 to 2700 UMI (nCount_RNA) and cells less than 10% mitochondrial content. Gene expression was normalized using the regularized negative binomial regression to normalize UMI count data (SCTransform) using 4000 variable features and the different samples (controls and irradiated). The features to use in the downstream integration procedure were determined using SelectIntegrationFeatures. After a pre-computed AnchorSet (FindIntergrationAnchors), the Seurat object were integrated using IntegrateData with the normalization method SCT. CellCycleScoring was applied to calculate the S.Score, G2M.Score and Phase. RunUMAP runs the Uniform Manifold Approximation and Projection (UMAP) dimensional reduction technique using 50 PCs. FindNeighbors computes the k.param nearest neighbors using dims = 1:50. Clusters of cells were identified by a shared nearest neighbor (SNN) modularity optimization bases clustering algorithm using FindClusters with resolutions from 0.2 to 2. The resolution 1.2 that segregates cells into 33 clusters was chosen based on Clustree graph showing the relashionship between clusterings at the different resolutions. UMAP plots were generated with UMAPPlot with default parameters.

The PrepSCTFindMarkers function was used to normalize gene expression for differential gene expression analysis among clusters, which was performed using FindAllMarkers on the SCT assay, with significance determined using a Wilcoxon rank sum test (p_val_adj < 0.05) and log(Fold-Change) threshold of 0.1. A histogram representation of *Dcx* (ENSMUSG00000031285) expression in the clusters corresponding to astrocytes and neural progenitors allowed us to place a threshold expression at 1. Cells with an expression lower than or equal to 1 correspond to *Dcx^Low^* cells and cells with an expression strictly higher than 1 to *Dcx^High^* cells. Then, we reapplied PrepSCTFindMarkers and FindAllMarkers to find differentially expressed genes between *Dcx^Low^* and *Dcx^High^* iNB.

##### Gene expression score

We used AddModuleScore_Ucell (Package *UCell* version 1.3.1) to score cells based on the expression of predefined specific gene lists of sorted s-qNSC, s-aNSC, s-TAP, s-iNB and s-mNB determined by microarrays using the Mann-Whitney U statistic ^40^. Each cell was then assigned an identity corresponding to the highest score amongst the signatures tested. Repartition of classes among clusters was visualized using the Circlize package ^58^.

For astrocyte and B-cell classification, we used the lists of Astrocyte and B-cell markers provided by Cebrian-Silla et al. (2021) ^14^ to score cells from the clusters 4 and 13 between 0 (min) and 1 (max). Cells were assigned to the highest score amongst both signatures.

##### Gene enrichment analysis

Comparisons of the specific gene lists of *Dcx^High^* s-iNB and *Dcx^Low^*s-iNB were performed with Biological pathways using the g:Profiler interface (https://biit.cs.ut.ee/gprofiler/gost).

### QUANTIFICATION AND STATISTICAL ANALYSIS

Non-parametric Mann-Whitney test was conducted using Prism 8.1.2 (GraphPad Software Inc., La Jolla, CA, RRID:SCR_002798). The statistical significance was set at p<0.05. The data are expressed as the mean ± SEM.

### DATA AVAILABILITY

The datasets generated for this article were deposited online on the ANNOTARE database (E-MTAB-12265, E-MTAB-12495). Scripts used for analysis are available on https://github.com/AChicheportiche/Bernou2023.

### Supplemental informations

**Figure S1:**
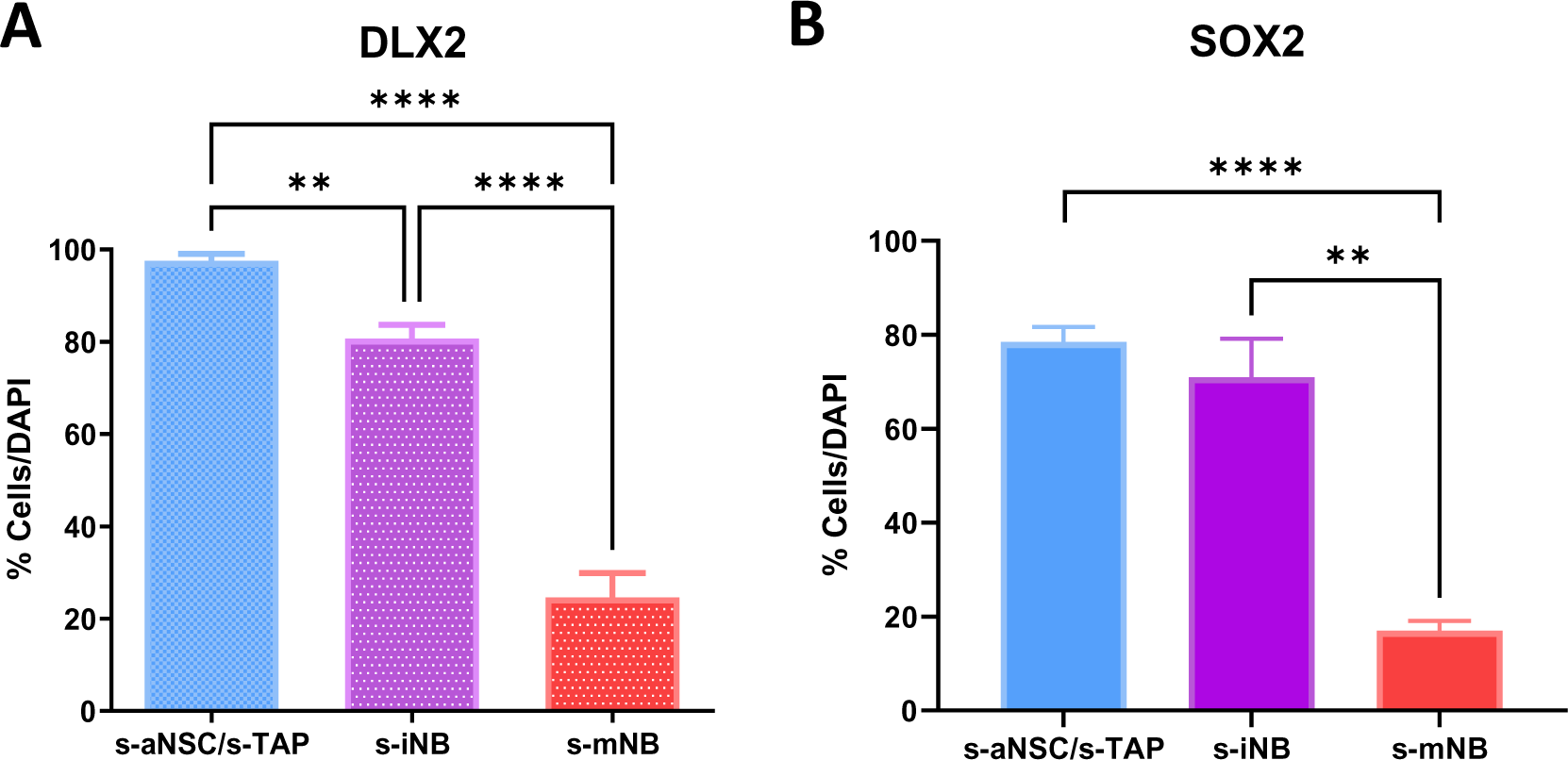
Expression of neural progenitor markers by SVZ cell-sorted populations. s-aNSC, s-TAP, s-iNB, and s-mNB cells were sorted from the SVZ of 2-month old C57Bl/6 mice, fixed and stained with antibodies against DLX2, a marker of proliferating progenitors (Doetsch et al. 2002) and SOX2 that plays an essential role in the maintenance of neural progenitors (Zhang et al. 2014) Quantifications of sorted SVZ cells expressing DLX2 **(A)** and SOX2 **(B)**. *p<0.05, **p<0.01, ***p<0.001.

**Figure S2:**
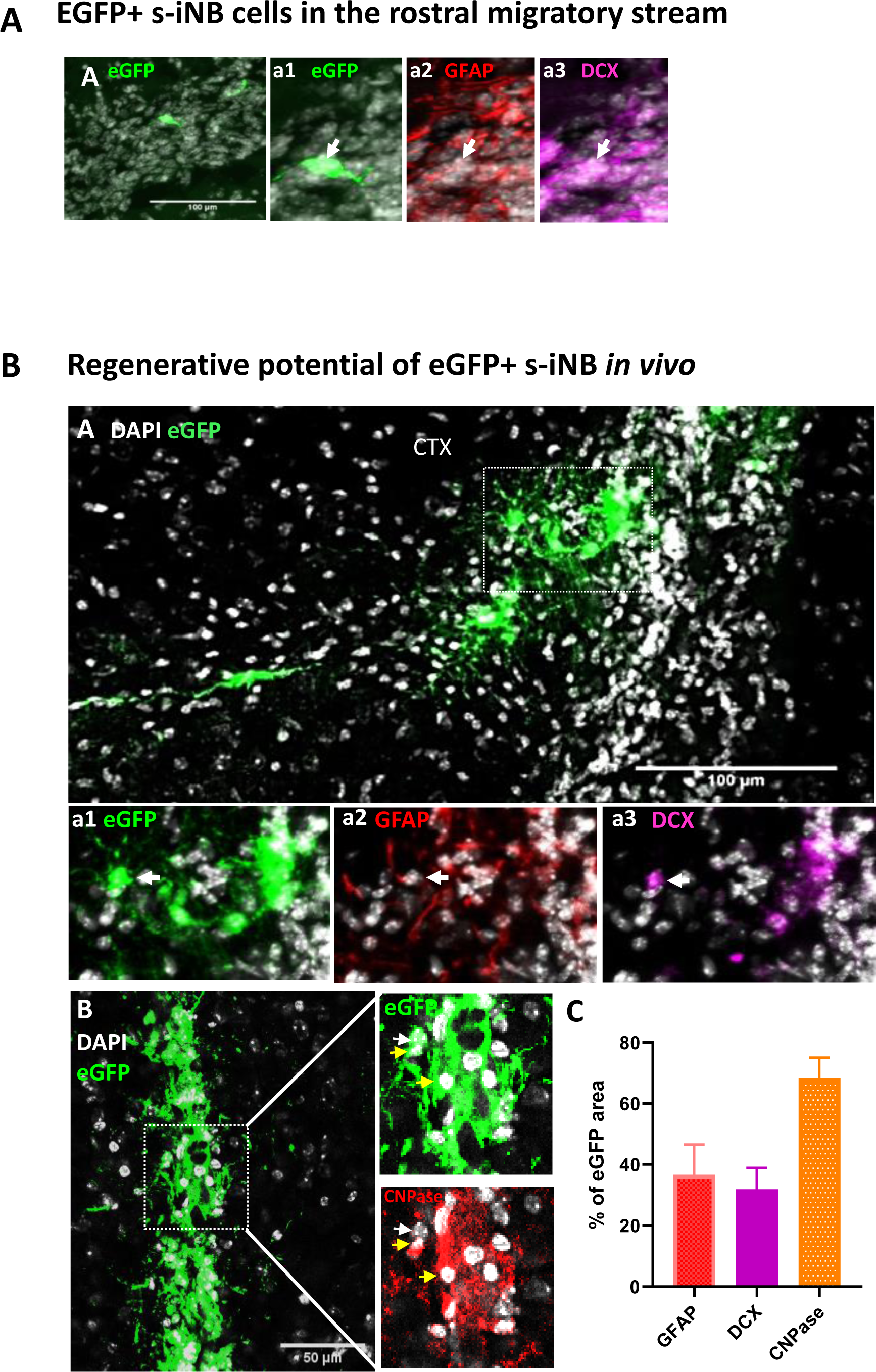
eGFP^+^ s-iNB cells in the rostral migratory stream and regenerative potential of eGFP+ s-iNB *in vivo*. Immunofluorescence of mouse brains transplanted unilaterally in 3 injection points near the dSVZ/RMS with eGFP^+^iNB cells isolated from β-actin:eGFP mice. Transplanted brains were analysed five weeks later. **(A)** Example of EGFP^+^iNB cell (white arrow) that has reached the rostral migratory stream and expressed DCX (inset a3), but not GFAP (inset a2). Insets are high magnification panels. **(B)** eGFP^+^ cells persisting at IS expressing astrocyte or oligodendrocyte markers. High magnifications of the inset1 (dotted line in A) showing eGFP^+^ GFAP^+^ cells (white arrow) **(a2)** and eGFP^+^DCX^+^ cells (white arrow) **(a3)**. High magnifications of the inset2 (dotted line in B) showing eGFP^+^ CNPase^+^ (yellow arrow) and eGFP^+^CNPase^−^ (white arrow). Scale bars= 100 µm. (C) Quantitative analysis of GFAP, CNPase or DCX antibodies staining of GFP-positive cells persisting at IS (where high number of grafted cells were found) in Figure S2. This was performed by using the NIS software measuring eGFP-, GFAP-, CNPase- and DCX-positive areas. The intersections between each marker and eGFP areas were determined as a percentage of staining. The results showed that approximately one third of eGFP+ cells expressed GFAP or DCX. The quantitative analysis of CNPase expression specific to eGFP+ cells was complicated by host cells, but an overexpression in eGFP-positive areas clearly revealed the expression of CNPase by a significant faction of these cells.

**Figure S3:**
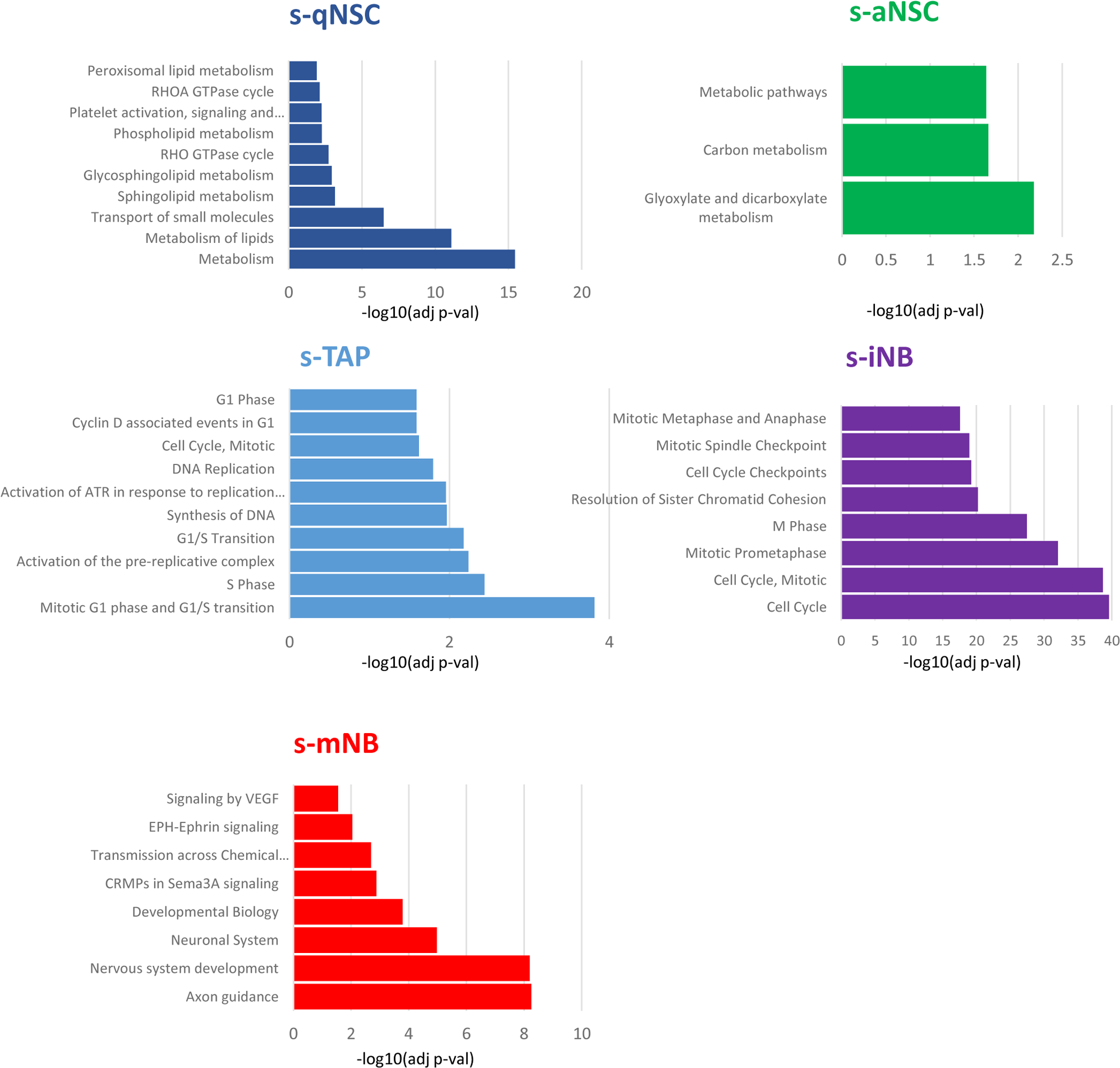
Functional enrichment analyses of the individual transcriptome signature of neurogenic populations. Comparisons of individual transcriptome signatures with various Gene Ontology gene sets and Biological pathways using the g:Profiler software highlight significant stage-dependant changes. S-qNSCs expressed genes related to Metabolism of lipids, Transport of small molecules. S-aNSC expressed genes involved in Glyoxylate and decarboxylate metabolism. S-TAP significantly displayed genes involved in Mitotic G1 phase and G1/S transition and S Phase. S-iNB specifically expressed Cell Cycle and Mitotic genes in contrast to s-mNB which are characterised by the expression of Axon guidance and Nervous system development genes.

**Figure S4:**
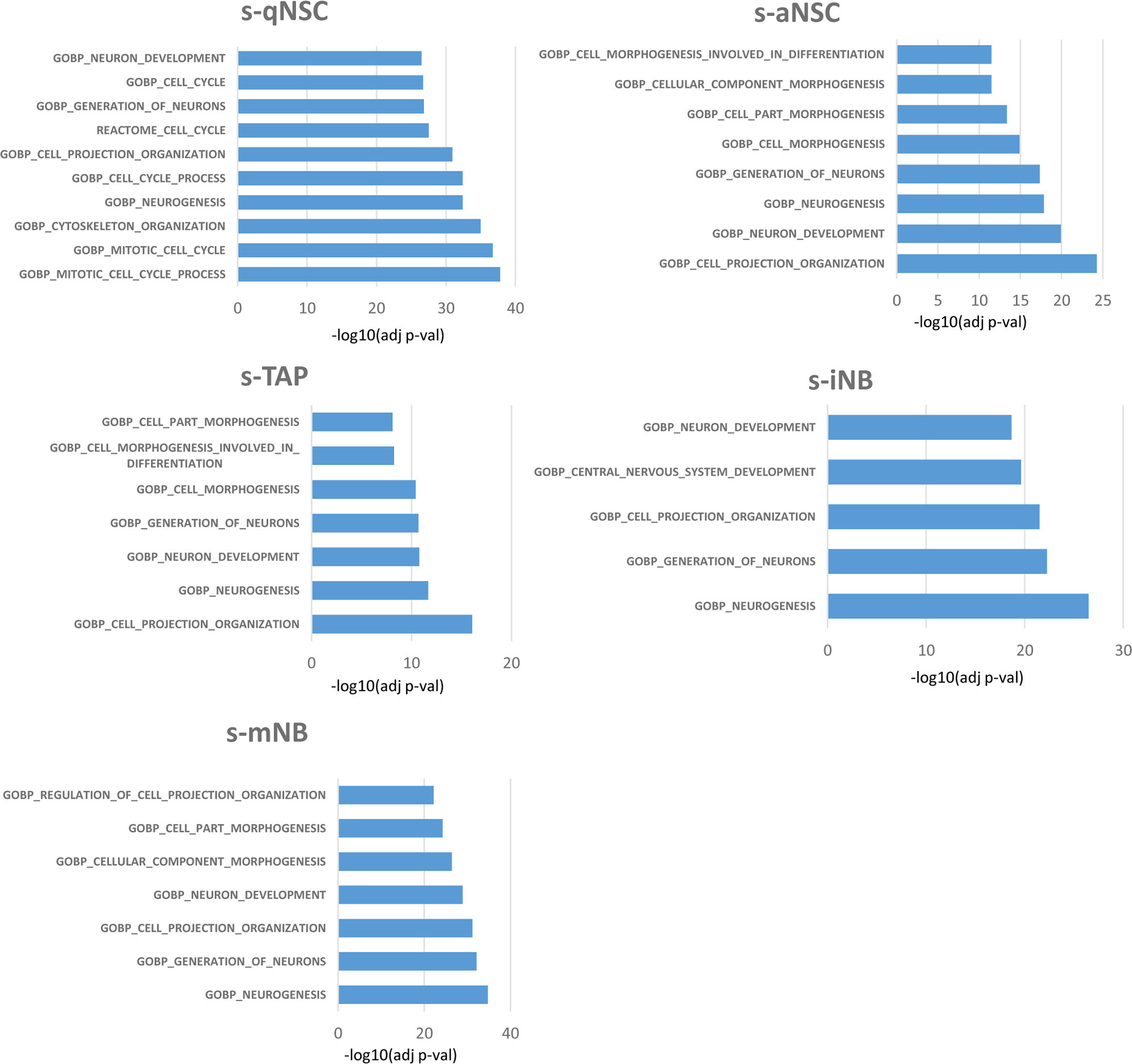
Enrichment of GO terms corresponding to mRNA splicing isoforms in the different types of sorted SVZ cells. Comparisons of individual mRNA splicing isoforms with the Gene Ontology Biological pathway sets using the Gene Set Enrichment Analysis (GSEA) interface. This analysis clearly revealed splicing in genes involved in neuron development and neurogenesis in all SVZ cells. Interestingly, this also showed that s-qNSC logically differed from the other cell types by splicing in genes involved in mitosis and cell cycle. More importantly, GO annotations in differentially spliced isoforms emphasized the close proximity of s-aNSC and s-TAP on one hand and s-iNB and s-mNB on the other hand confirming than s-iNB are closer to s-mNB than s-TAP.

**Figure S5:**
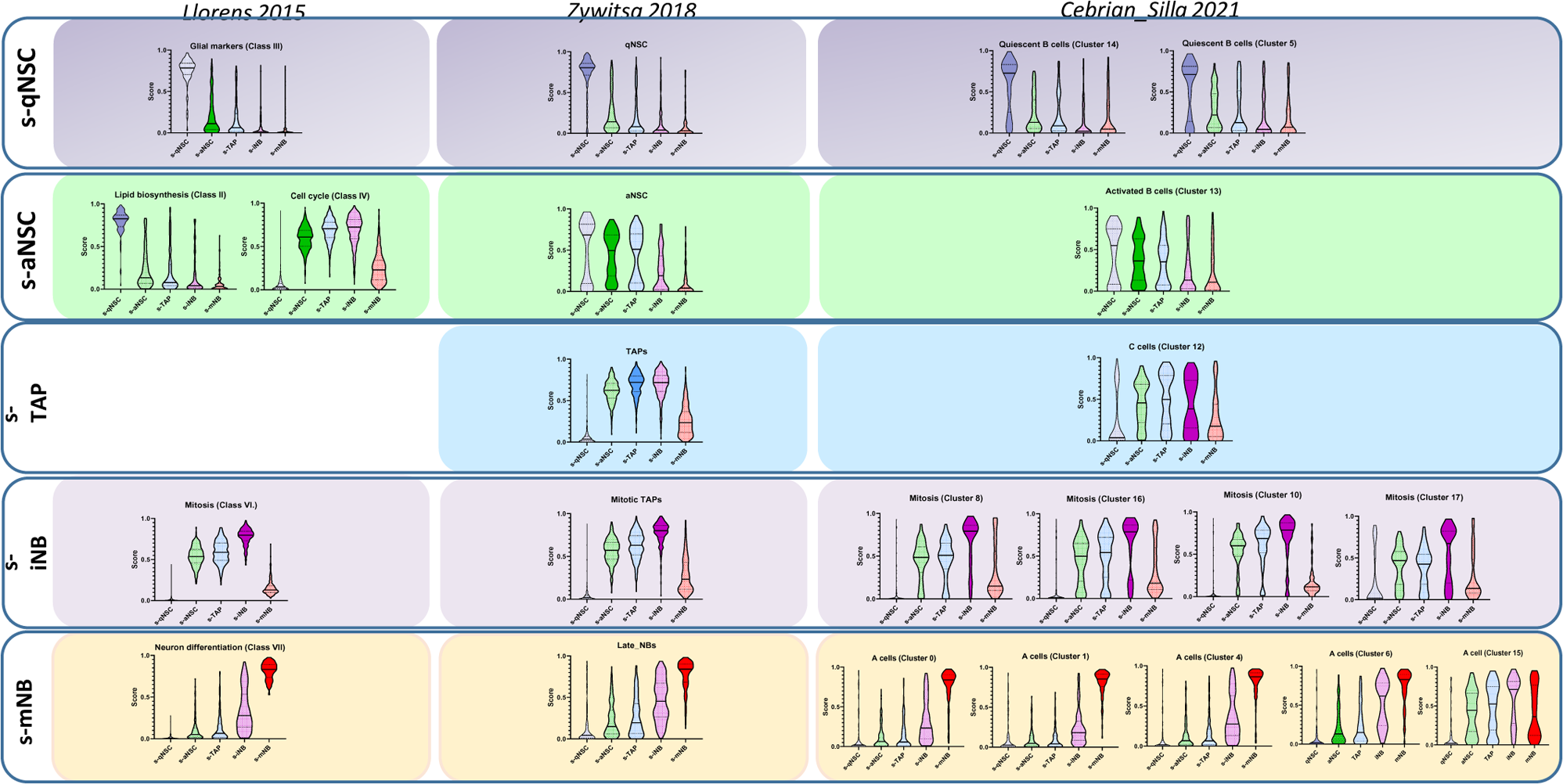
Correspondence between the molecular signatures of sorted neural progenitor populations in our study and available sc-RNA-seq datasets. Violin diagrams representing the expression of selected clusters from published sc-RNAseq analysis in our Clariom microarray^14,17,32^. S-qNSC perfectly matched the expression of Glial markers (Class III)^32^, qNSC^17^ and Quiescent B cells (clusters 5 and 14)^14^. In contrast, s-aNSC poorly matched with Lipid biosynthesis (Class III) and Cell cycle (Class IV)^32^, aNSC^17^ and Activated B cells (Cluster 13)^14^. Similarly, s-TAP poorly matched with TAP^17^ and C cells (Cluster 12)^14^. S-iNB perfectly fit with Mitosis (Class VI), mitotic TAP (mTAP) (Zywitza et al., 2018) and Mitosis (Cluster 8, 10,16 and 17)^14^. s-mNB overlapped genes related to Neuron differentiation (Class VII), Late NBs ^17^ and A cells (clusters 0, 1 and 4).

**Figure S6:**
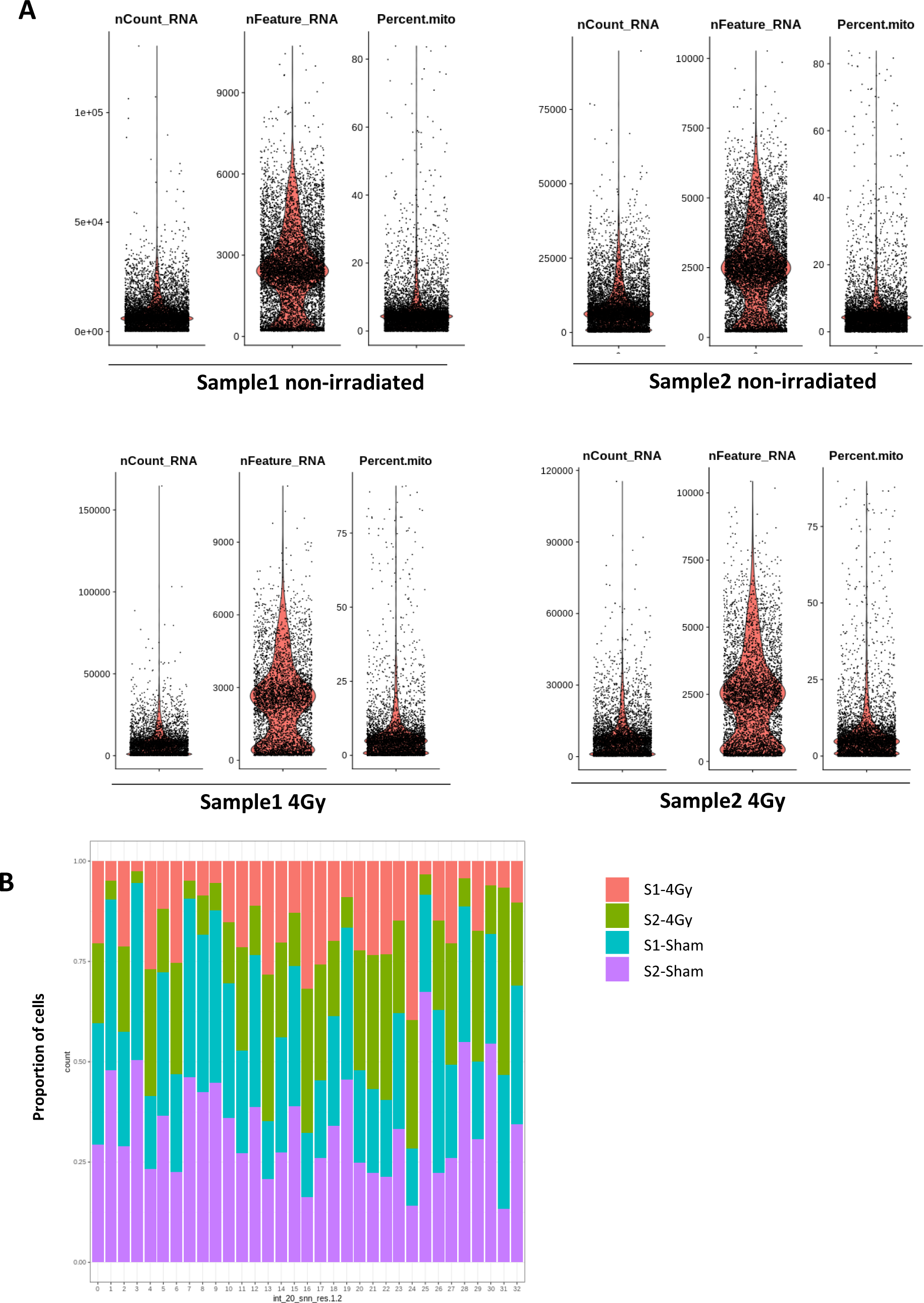
Quality controls of scRNA-seq on whole SVZ tissues. **(A)** Violin diagrams of the nCount_RNA, the nFeature_RNA and the mitochondrial percentage (Percent.mito) before applying cut-offs adapted to each non-irradiated and irradiated samples. **(B)** Proportion of cells from the 4 samples in the 33 clusters.

**Figure S7:**
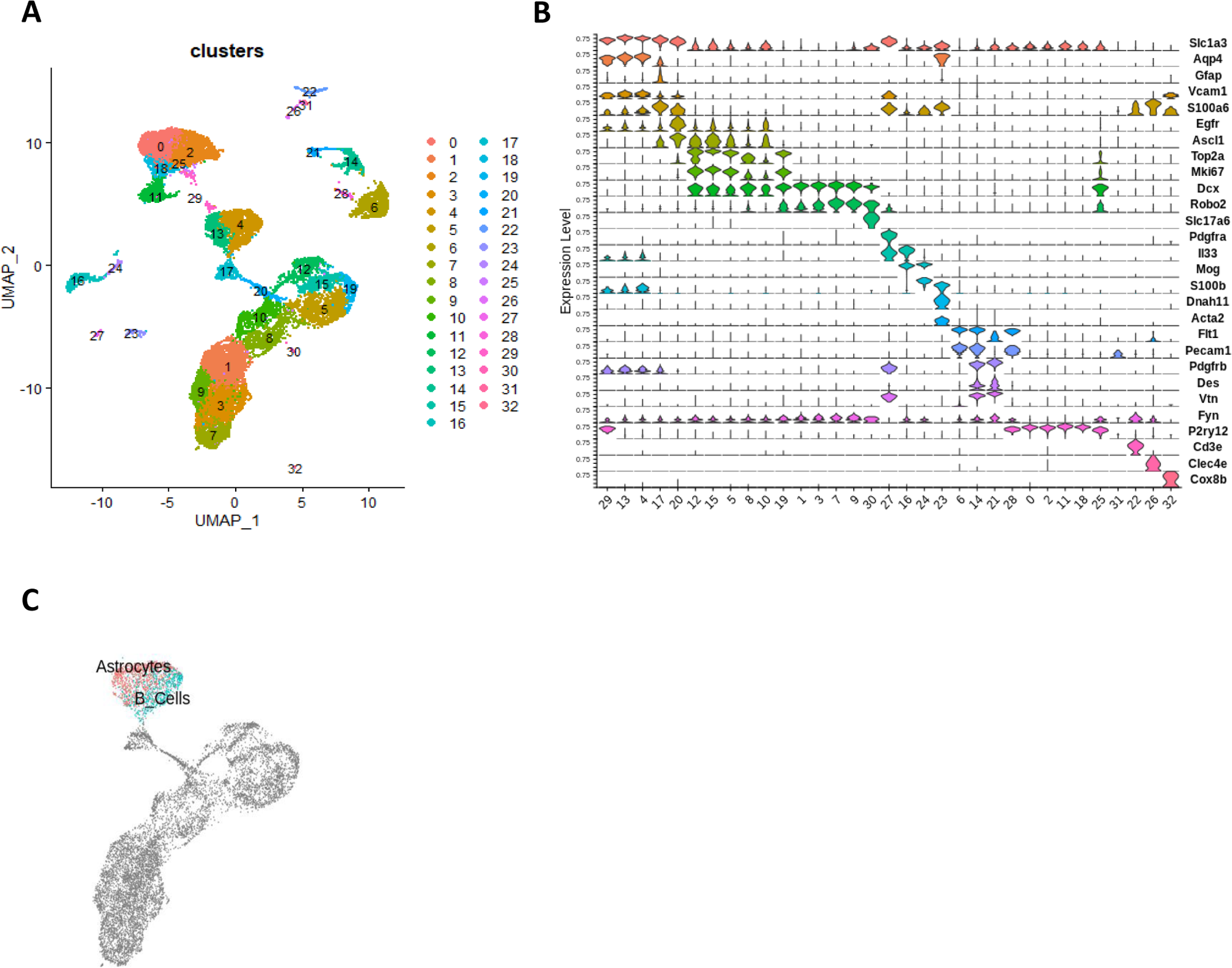
Annotation of sc-RNAseq clusters. **(A)** UMAP at the resolution 1.2 combining scRNA-seq datasets of unirradiated and 4Gy irradiated cells segregated into 33 clusters. **(B**) Violin diagrams illustrating the expression level of known cell markers in the 33 clusters according to the literature. **(C)** UMAP of top normalized UCellScore Astrocyte- and the B-cell specific genes described by Cebrian-Silla et al ^14^ for cells from clusters 4 and 13.

**Figure S8:**
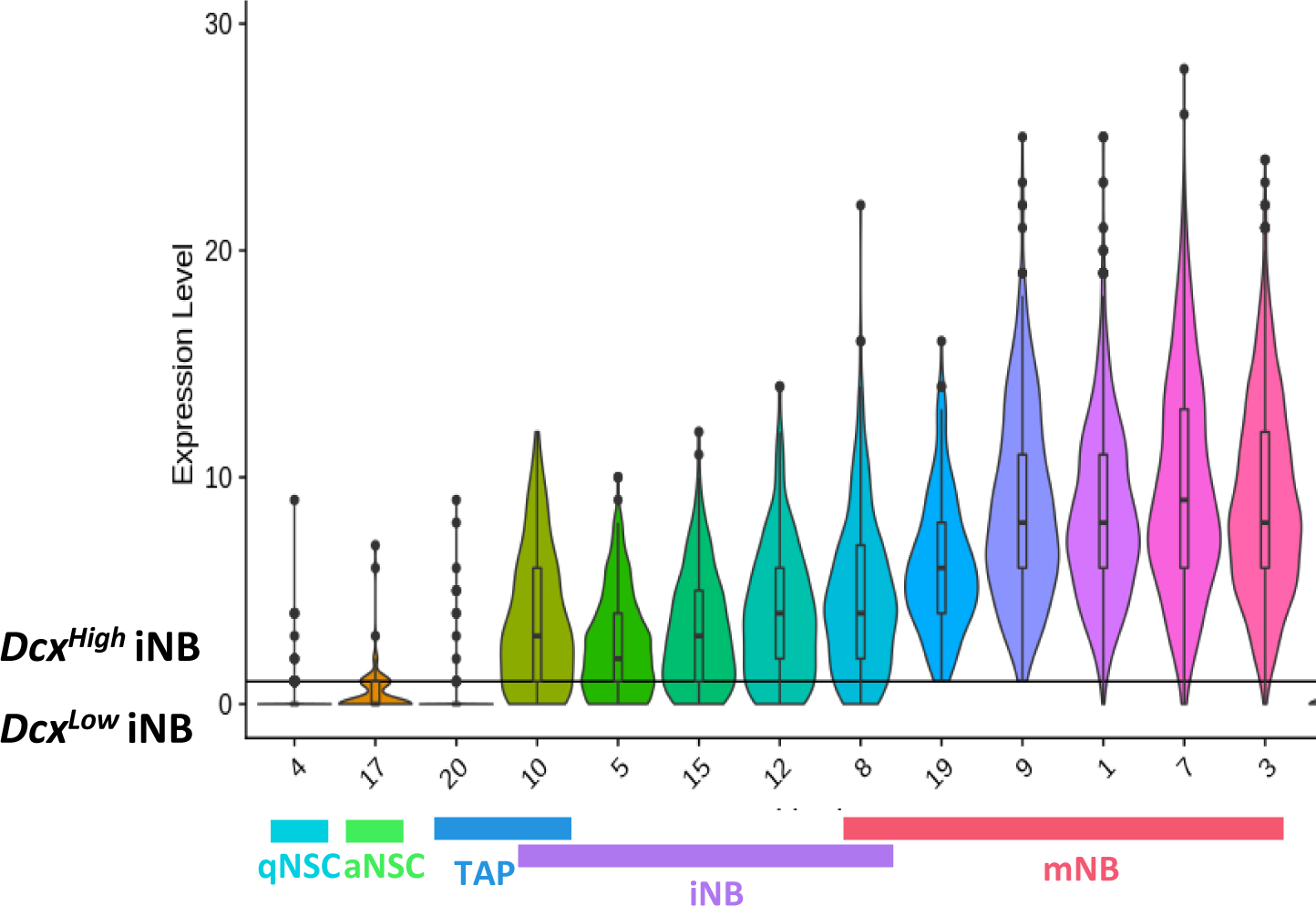
Expression of *Dcx* in scRNA-seq neurogenic clusters. Violin diagram of the expression of *Dcx* in cells from neurogenic clusters illustrating two populations of iNB: iNB *Dc^Hiigh^*and iNB *Dcx^Low^* cells.

**Figure S9:**
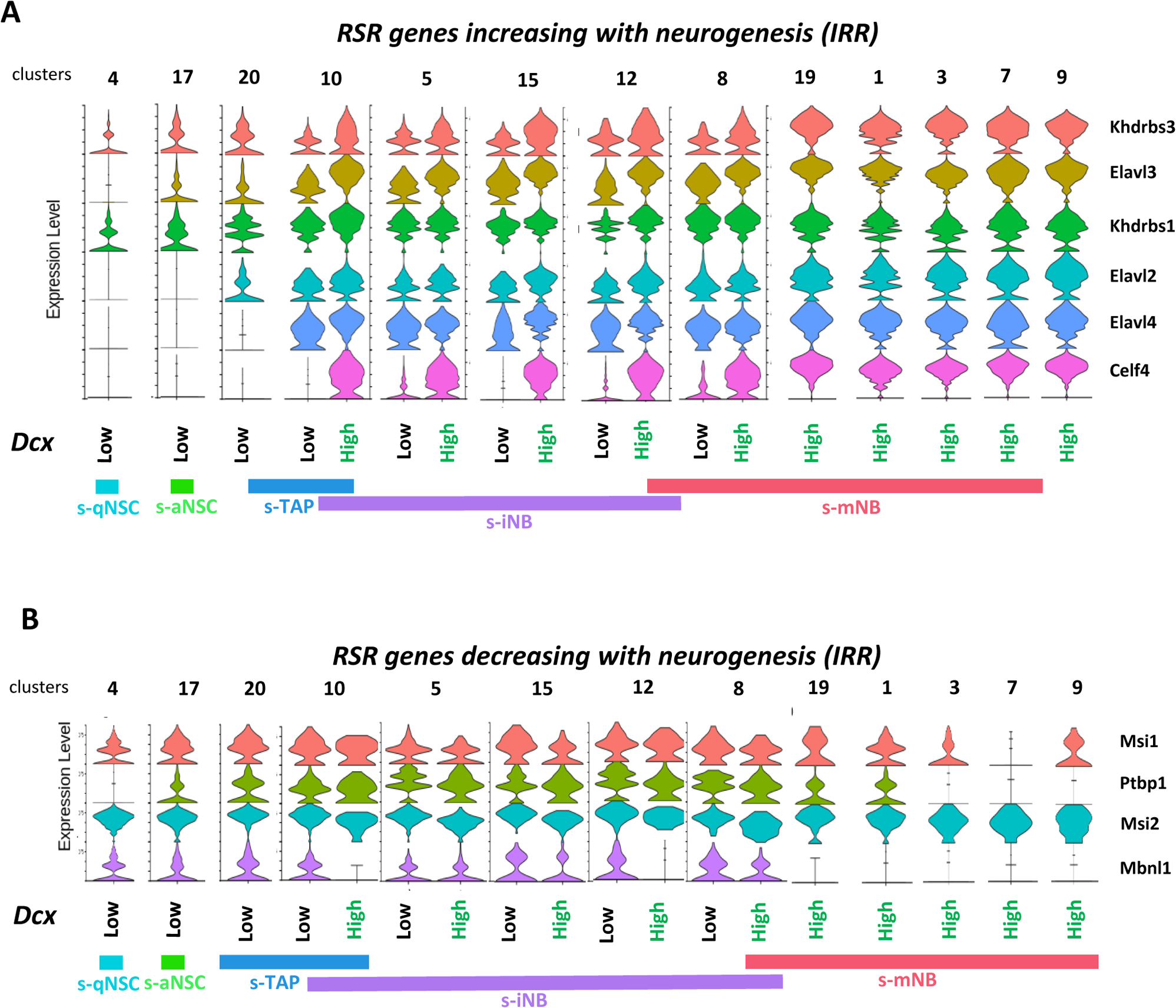
Expression of RSR genes in scRNA-seq neurogenic clusters in irradiated brains. Violin diagrams illustrating the expression of the RSR genes increasing **(A)** or decreasing **(B)** with neurogenesis.

**Figure S10:**
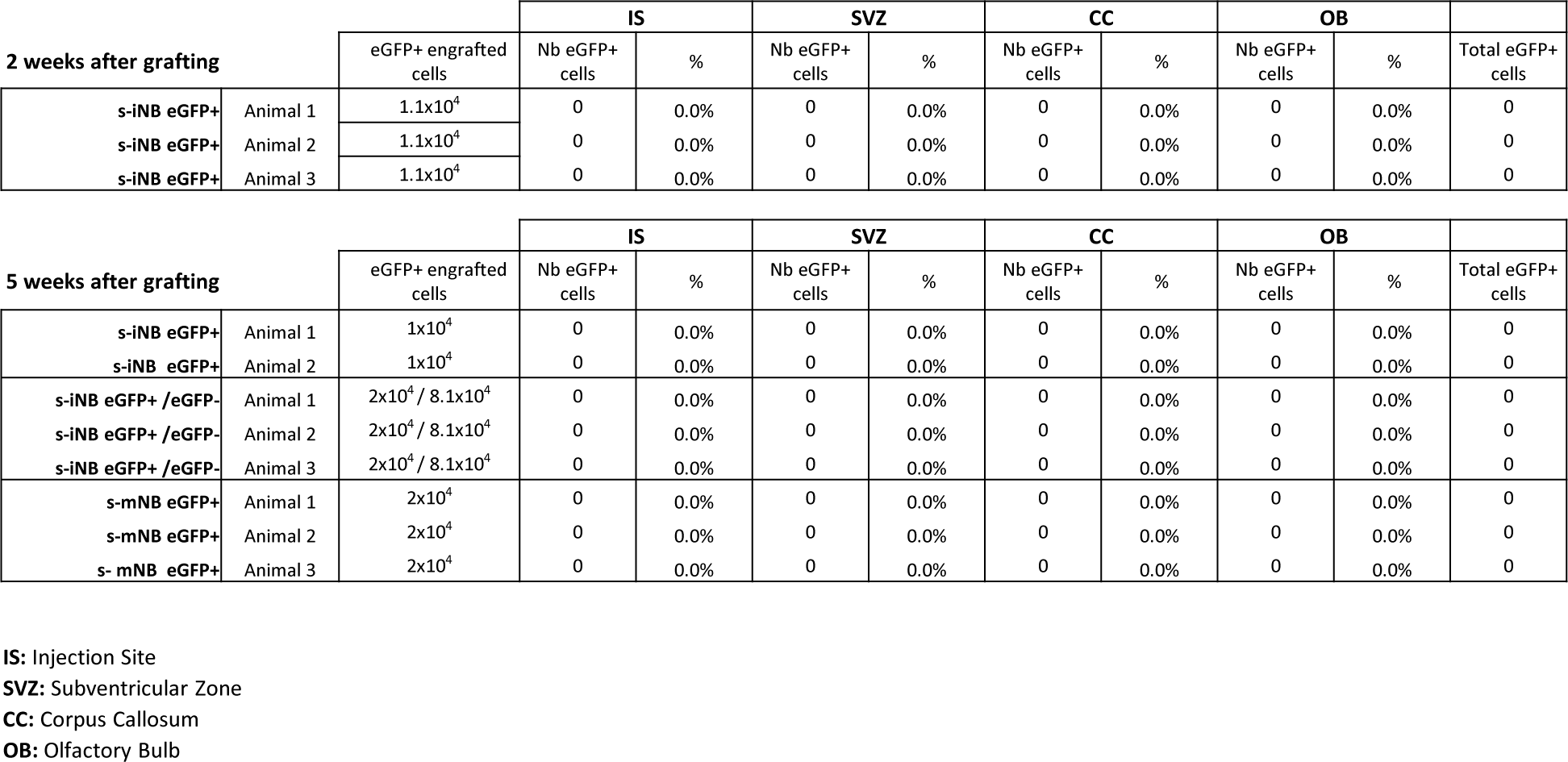
Transplantations of eGFP^+^s-iNB, eGFP^−^s-iNB and eGFP^+s-^mNB freshly isolated from DCX-CreERT2::CAG-floxed-eGFP mice model. Table recapitulating the immunohistological analyses of the numbers of eGFP^+^ cells in different regions in recipient C57Bl6/J mice brains, 2 and 5 weeks after transplantation.

**Supp_data_1:** Transcriptomic signatures of s-qNSC, s-aNSC, s-TAP, s-iNB and s-mNB (Clariom assay) and corresponding g:Profiler Gene Ontology analyses.

**Supp_data_2:** Differentially spliced genes (DSG) unique to s-qNSC, s-aNSC, s-TAP, s-iNB and s-mNB and corresponding Gene Enrichment Analyses (GSEA).

**Supp_data_3:** Differentially spliced genes in each transition along SVZ neurogenesis and corresponding g.Profiler Gene Ontology analyses.

**Supp_data_4:** Differentially expressed genes (DEG) among RNA splicing regulators (RSR) genes in each transition along SVZ neurogenesis.

**Supp_data_5:** Identification of astrocytes and qNSC among clusters 4 and 13 from our scRNAseq dataset using the transcriptomic signatures of Astrocytes and B_cells described by Cebrian et al. 2021.

**Supp_data_6:** Differentially expressed genes of s-INB *Dcx*^High^ and s-iNB *Dcx*^Low^ cells and functional profiling using g:Profiler interface.

