## Supplementary material for "Switching of RNA splicing regulators in immature neuroblasts: a key step in adult neurogenesis": METHOD

| REAGENT or RESSOURCE | SOURCE | IDENTIFIER |
| --- | --- | --- |
| Antibodies |  |  |
| PE-rat Anti-mouse CD24 (clone 1/69) | BD Biosciences | Cat#307567;RRID:AB_2034001 |
| FITC-rat Anti-mouse CD15 (clone MMA) | BD Biosciences | Cat#340703; RRID:AB_400103 |
| BV421-mouse Anti-human CD15 (Clone W6D3) | BioLegend | Cat#323039; RRID:AB_2566519 |
| Alexa647-conjugated epidermal growth factor (EGF) ligand (clone 30H45L48) | ThermoFisher Scientific | Cat#MA7-00308-A647 |
| Goat Anti-GFP | Abcam | Cat#ab6673 |
| Rabbit Anti-GFP | Abcam | Cat#ab290 |
| Mouse Anti-mouse O4 (clone 81) | Merck Millipore | Cat#MAB345 |
| Rabbit Anti-mouse NG2 | Merck Millipore | Cat#AB5320 |
| Goat Anti-mouse Dcx C-18 | Santa Cruz | Cat#sc-8066 |
| Rabbit Anti-mouse Dcx | CellSignaling | Cat#4604 |
| Rabbit Anti-mouse Tubulin β3 (TUBB3) | Covance | Cat#PRB-435P |
| Mouse Anti-mouse Gfap (clone GA5) | Merck Millipore | Cat#MAB3402 |
| Rabbit Anti-mouse Gfap | Merck Millipore | Cat#G9269 |
| Rat Anti-CD133 (clone 13A4) | Merck Millipore | Cat#MAB4310 |
| Mouse Anti-mouse CNPase (clone 11-5B) | Merck Millipore | Cat#MAB326 |
| Goat Anti-mouse Olig2 | Biotechne | Cat#AF2418 |
| Mouse Anti-mouse NeuN (clone A60) | Merck Millipore | Cat#MAB377 |
| Alexa Fluor 594_ Donkey anti-Mouse IgG (H+L) Secondary Antibody | ThermoFisher Scientific | Cat#A21203 |
| Alexa Fluor 488_Donkey anti-Mouse IgG (H+L) Secondary Antibody | ThermoFisher Scientific | Cat#A21202 |
| Alexa Fluor 647_ Donkey anti-Mouse IgG (H+L) Secondary Antibody | ThermoFisher Scientific | Cat#A32787 |
| Chemical, peptides, and recombinant proteins |  |  |
| Albumin Bovine Serum 10% | Merck Millipore | Cat#9048-46-8 |
| Percoll | Merck Millipore | Cat#GE17-0891-01 |
| DPBS w/o calcium w/o magnesium | ThermoFisher Scientific | Cat#12559069 |
| NeuroCult Basal medium | STEMCELL Technologies | Cat#05700 |
| Heparin Solution | STEMCELL Technologies | Cat#07980 |
| Epidermal Growth Factor Protein, human recombinant | Merck Millipore | Cat#GF144 |
| Fibroblast growth Factor basic, human recombinant | Merck Millipore | Cat#GF003AF |
| Poly-L-ornithine solution | Merck Millipore | P4957 |
| Poly-D-lysine solution | Merck Millipore | 25988-63-0 |
| DMEM:F12 medium | ThermoFisher Scientific | 31331028 |
| B-27-Supplement Minus Antioyxidants (AO) | ThermoFisher Scientific | Cat#10889038 |
| Fetal Bovine serum | ThermoFisher Scientific | Cat#10082139 |
| D-(+)-Glucose | Merck Millipore | Cat#G7021 |
| Sodium bicarbonate | Merck Millipore | Cat#S5761 |
| HEPES sodium salt | Merck Millipore | Cat#H3784 |
| Insulin from bovine pancreas | Merck Millipore | Cat#I1882 |
| Apo-Transferrin human | Merck Millipore | Cat#T2252 |
| Progesterone | Merck Millipore | Cat#P6149 |
| Putrescine | Merck Millipore | Cat#P7505 |
| Sodium selenite | Merck Millipore | Cat#S9133 |
| Papain | Worthington Biochemical | Cat#LK003150 |
| Tamoxifen | Sigma Aldrich | Cat#T5648 |
| Corn oil | Sigma Aldrich | Cat#C8267 |
| Solution de paraformaldéhyde, 4 % en PBS | ThermoFisher Scientific | Cat#15670799 |
| Triton X-100 | Sigma Aldrich | Cat# 11332481001 |
| Propidium iodide | Sigma Aldrich | Cat#P4170 |
| Hoechst 33258 | ThermoFisher Scientific | Cat#H3569 |
| Critical Commercial Assays |  |  |
| Red blood Cell lysis solution | Milteny Biotec | Cat#130-094-183 |
| Dead Cell Removal Kit | Milteny Biotec | Cat#130-090-101 |
| RNeasy Micro Kit | Qiagen | Cat#74004 |
| RNase-Free DNase Set | Qiagen | Cat#79254 |
| AgilentHigh Sensitivity DNA Kit | LabChip | Cat#5067-4626 |
| Chromium Next GEM Chip G Single Cell Kit | 10X GENOMICS | Cat#PN-1000127 |
| Dual Index Kit Plate TT Set A | 10X GENOMICS | Cat#PN-1000215 |
| Deposited data |  |  |
| List of Significantly Upregulated Genes in Subclusters, Related to Figures 3 and S4. | Zywitsa et al., 2018 | <https://doi.org/10.1016/j.celrep.2018.11.003>  TableS3  File:1-s2.0-S2211124718317327-mmc4.xlsx |
| B cells vs. astrocytes: DE genes and GO terms | Cebrian et al. 2021 | <https://doi.org/10.7554/eLife.67436>  Supplementary file 1 |
| scRNAseq datasets from the University of California Santa Cruz Cell Browser | Cebrian et al. 2021 | <https://svzneurogeniclineage.cells.ucsc.edu.> |
| List of cluster gene sets and GO terms | Llorens et al. 2015 | <https://doi.org/10.1016/j.stem.2015.07.002>  File:1-s2.0-S193459091500301X-mmc3.xls |
| Softwares |  |  |
| RStudio | RStudio | <https://www.rstudio.com/> |
| Transcriptome Analysis Console (TAC 4.0) | ThermoFisher Scientific | -Ritchie ME et al. (2015) limma powers differential expression analyses for RNAsequencing and microarray studies. Nucleic Acids Research 43(7):e47.  -Romero JP et al. (2016) EventPointer: an effective identification of alternative splicing  events using junction arrays. BMC Genomics 17:467. |
| Other |  |  |
| 70µm cell strainer | ThermoFisher Scientific | Cat#10788201 |
| 2.5µl, Model 62 RN SYR, Small Removable NDL, 22s ga, 2 in, point style 3 | Hamilton | Cat#87942 |
| Small Hub 33G | Hamilton | Cat#074750 |
