## Supplementary material for "Switching of RNA splicing regulators in immature neuroblasts: a key step in adult neurogenesis": script_R_6: 6_Dcx_analysis.html

Dcx classes creation and irradiation ratios


### Dcx classes creation and irradiation ratios

Here we define DcxHigh and DcxLow cells amongst s-iNB and examine their relative abundance

```
library(tidyverse)
```

```
## ── Attaching packages ─────────────────────────────────────── tidyverse 1.3.2 ──
## ✔ ggplot2 3.4.0      ✔ purrr   0.3.5 
## ✔ tibble  3.1.8      ✔ dplyr   1.0.10
## ✔ tidyr   1.2.1      ✔ stringr 1.4.1 
## ✔ readr   2.1.2      ✔ forcats 0.5.2 
## ── Conflicts ────────────────────────────────────────── tidyverse_conflicts() ──
## ✖ dplyr::filter() masks stats::filter()
## ✖ dplyr::lag()    masks stats::lag()
```

```
library(dplyr)
library(patchwork)
library(Seurat, lib.loc = "/opt/rstudio-server_conda/conda/envs/rstudio-server_4.1.0/lib/R/library/")
```

```
## Attaching SeuratObject
## Attaching sp
```

Loading required objects This analysis is focused on NPC populations, therefore we will use a subset of the base object

```
SCT_obj <- readRDS("Objects/SCT_obj.rds")
Biomart_annotated <- read.csv("Analysis_tools/Biomart_annotated.csv")
geneIDs <- # For easy calling of genes through external gene name
  Biomart_annotated[!duplicated(Biomart_annotated$external_gene_name),1]
names(geneIDs) <- Biomart_annotated[!duplicated(Biomart_annotated$external_gene_name),2]
NPC_obj <- subset(SCT_obj, NPC == "NPC")
```

```
dcx_expression <- FetchData( #FetchData sur le slot data
    NPC_obj,
    vars = c(geneIDs["Dcx"], "ident", "Status"),
    assay = "SCT",
    slot = "counts"
  ) %>%
    mutate(dcx = ifelse(ENSMUSG00000031285 > 1,"DcxHigh","DcxLow"))
dcx_lvl <- dcx_expression$dcx
names(dcx_lvl) <- rownames(dcx_expression)
SCT_obj <- AddMetaData(SCT_obj, metadata = dcx_lvl, col.name = "dcx_lvl")

colnames(dcx_expression)[1:4] <- c("DCX_val", "cluster", "Status", "dcx")  

write.csv(table(dcx_expression$cluster, 
                dcx_expression$Status,
                dcx_expression$dcx),
          file = "Outputs/DCX_cluster_status.csv")
```

Saving object with all metadata

```
saveRDS(object = SCT_obj, file = "Objects/Ber23_SVZ_IRR.rds")
sessionInfo()
```

```
## R version 4.1.0 (2021-05-18)
## Platform: x86_64-conda-linux-gnu (64-bit)
## Running under: Ubuntu 20.04.4 LTS
## 
## Matrix products: default
## BLAS/LAPACK: /opt/rstudio-server_conda/conda/envs/rstudio-server_4.1.0/lib/libopenblasp-r0.3.15.so
## 
## locale:
##  [1] LC_CTYPE=en_US.UTF-8       LC_NUMERIC=C              
##  [3] LC_TIME=en_US.UTF-8        LC_COLLATE=en_US.UTF-8    
##  [5] LC_MONETARY=en_US.UTF-8    LC_MESSAGES=en_US.UTF-8   
##  [7] LC_PAPER=en_US.UTF-8       LC_NAME=C                 
##  [9] LC_ADDRESS=C               LC_TELEPHONE=C            
## [11] LC_MEASUREMENT=en_US.UTF-8 LC_IDENTIFICATION=C       
## 
## attached base packages:
## [1] stats     graphics  grDevices utils     datasets  methods   base     
## 
## other attached packages:
##  [1] sp_1.4-7           SeuratObject_4.1.0 Seurat_4.1.1       patchwork_1.1.2   
##  [5] forcats_0.5.2      stringr_1.4.1      dplyr_1.0.10       purrr_0.3.5       
##  [9] readr_2.1.2        tidyr_1.2.1        tibble_3.1.8       ggplot2_3.4.0     
## [13] tidyverse_1.3.2   
## 
## loaded via a namespace (and not attached):
##   [1] readxl_1.4.1          backports_1.4.1       plyr_1.8.7           
##   [4] igraph_1.3.1          lazyeval_0.2.2        splines_4.1.0        
##   [7] listenv_0.8.0         scattermore_0.8       digest_0.6.31        
##  [10] htmltools_0.5.4       fansi_1.0.3           magrittr_2.0.3       
##  [13] tensor_1.5            googlesheets4_1.0.0   cluster_2.1.3        
##  [16] ROCR_1.0-11           tzdb_0.3.0            globals_0.15.0       
##  [19] modelr_0.1.8          matrixStats_0.62.0    spatstat.sparse_2.1-1
##  [22] colorspace_2.0-3      rvest_1.0.2           ggrepel_0.9.1        
##  [25] haven_2.5.0           xfun_0.31             crayon_1.5.2         
##  [28] jsonlite_1.8.4        progressr_0.10.0      spatstat.data_2.2-0  
##  [31] survival_3.3-1        zoo_1.8-10            glue_1.6.2           
##  [34] polyclip_1.10-0       gtable_0.3.1          gargle_1.2.0         
##  [37] leiden_0.4.2          future.apply_1.9.0    abind_1.4-5          
##  [40] scales_1.2.1          DBI_1.1.2             spatstat.random_2.2-0
##  [43] miniUI_0.1.1.1        Rcpp_1.0.8.3          viridisLite_0.4.0    
##  [46] xtable_1.8-4          reticulate_1.22       spatstat.core_2.4-4  
##  [49] htmlwidgets_1.5.4     httr_1.4.4            RColorBrewer_1.1-3   
##  [52] ellipsis_0.3.2        ica_1.0-2             pkgconfig_2.0.3      
##  [55] uwot_0.1.11           sass_0.4.4            dbplyr_2.1.1         
##  [58] deldir_1.0-6          utf8_1.2.2            tidyselect_1.2.0     
##  [61] rlang_1.0.6           reshape2_1.4.4        later_1.3.0          
##  [64] munsell_0.5.0         cellranger_1.1.0      tools_4.1.0          
##  [67] cachem_1.0.6          cli_3.4.1             generics_0.1.3       
##  [70] broom_0.8.0           ggridges_0.5.3        evaluate_0.15        
##  [73] fastmap_1.1.0         yaml_2.3.6            goftest_1.2-3        
##  [76] knitr_1.39            fs_1.5.2              fitdistrplus_1.1-8   
##  [79] RANN_2.6.1            pbapply_1.5-0         future_1.25.0        
##  [82] nlme_3.1-157          mime_0.12             xml2_1.3.3           
##  [85] compiler_4.1.0        rstudioapi_0.13       plotly_4.10.0        
##  [88] png_0.1-7             spatstat.utils_2.3-1  reprex_2.0.1         
##  [91] bslib_0.4.2           stringi_1.7.8         rgeos_0.5-9          
##  [94] lattice_0.20-45       Matrix_1.4-1          vctrs_0.5.1          
##  [97] pillar_1.8.1          lifecycle_1.0.3       spatstat.geom_2.4-0  
## [100] lmtest_0.9-40         jquerylib_0.1.4       RcppAnnoy_0.0.19     
## [103] data.table_1.14.2     cowplot_1.1.1         irlba_2.3.5          
## [106] httpuv_1.6.5          R6_2.5.1              promises_1.2.0.1     
## [109] KernSmooth_2.23-20    gridExtra_2.3         parallelly_1.31.1    
## [112] codetools_0.2-18      MASS_7.3-57           assertthat_0.2.1     
## [115] withr_2.5.0           sctransform_0.3.3     mgcv_1.8-40          
## [118] parallel_4.1.0        hms_1.1.1             grid_4.1.0           
## [121] rpart_4.1.16          rmarkdown_2.14        googledrive_2.0.0    
## [124] Rtsne_0.16            shiny_1.7.1           lubridate_1.8.0
```
