## Supplementary material for "Switching of RNA splicing regulators in immature neuroblasts: a key step in adult neurogenesis": script_R_1: 1_Libraries_installation.html

Version and necessary libraries


Code 

- Show All Code
- Hide All Code
- Download Rmd

### Version and necessary libraries


```
```r
R.Version()
```

```
<!-- rnb-source-end -->

<!-- rnb-chunk-end -->


<!-- rnb-text-begin -->


Necessary librairies


<!-- rnb-text-end -->


<!-- rnb-chunk-begin -->


<!-- rnb-source-begin eyJkYXRhIjoiYGBgclxuaW5zdGFsbC5wYWNrYWdlcyhuZWNlc3NhcnlfbGlicmFpcmllcylcblxuYGBgIn0= -->

```r
install.packages(necessary_librairies)
```


```
Error in install.packages : Updating loaded packages
```


BiocManager::install(“satijalab/Seurat@v4.1.0”)
