## Supplementary material for "Switching of RNA splicing regulators in immature neuroblasts: a key step in adult neurogenesis": script_R_2: 2_Fig_S6A_Quality_Sample_Integration.html

Sample loading and quality control


### Sample loading and quality control

##### Loading Samples and set up

Loading libraries Note: please make sure to use both Seurat v4.1.1 and SeuratObject v4.1.0 to reproduce this analysis, as later versions of either packages change the UMAP at the RunUMAP step.

```
library(Seurat)
```

```
## Attaching SeuratObject
```

```
## Attaching sp
```

```
library(remotes)
library(tidyverse)
```

```
## ── Attaching packages
## ───────────────────────────────────────
## tidyverse 1.3.2 ──
```

```
## ✔ ggplot2 3.4.0      ✔ purrr   1.0.1 
## ✔ tibble  3.1.7      ✔ dplyr   1.0.10
## ✔ tidyr   1.2.0      ✔ stringr 1.5.0 
## ✔ readr   2.1.2      ✔ forcats 0.5.2 
## ── Conflicts ────────────────────────────────────────── tidyverse_conflicts() ──
## ✖ dplyr::filter() masks stats::filter()
## ✖ dplyr::lag()    masks stats::lag()
```

```
library(dplyr)
library(biomaRt)
library(ggplot2)
library(patchwork)

Ensembl_gene <- useEnsembl(biomart = "genes", dataset = "mmusculus_gene_ensembl")
```

Loading all samples and creating list of seurat objects; teh list will be used until integration

```
Sample_names <- c("CTRL_1", "CTRL_2", "4Gy_1", "4Gy_2")

Sample_list <- 
  lapply(X = Sample_names, FUN = function(sample){
    Read10X(data.dir = paste0("All_inputs/Samples/SVZ_", sample),
            gene.column = 1,
            cell.column = 1,
            unique.features = TRUE,
            strip.suffix = TRUE) %>%
      CreateSeuratObject(project = sample,
                         assay = "RNA",
                         min.cells = 0,
                         min.features = 1)
  })
```

```
## as(<dgTMatrix>, "dgCMatrix") is deprecated since Matrix 1.5-0; do as(., "CsparseMatrix") instead
```

Use BioMart as a reference for all features (features are shared amongst samples)

```
Biomart_annotated <- getBM(attributes = c("ensembl_gene_id", 
                                          "external_gene_name",
                                          "description",
                                          "chromosome_name"),
                           filters = "ensembl_gene_id", 
                           values = rownames(Sample_list[[1]]),
                           mart = Ensembl_gene) 
write_csv(Biomart_annotated, file = "Analysis_tools/Biomart_annotated.csv")
```

Create variables for easy calling of Ensembl Id from External Gene Name (or vice-versa)

```
geneIDs <- # For easy calling of genes through external gene name
  Biomart_annotated[!duplicated(Biomart_annotated$external_gene_name),1]
names(geneIDs) <- Biomart_annotated[!duplicated(Biomart_annotated$external_gene_name),2]
write.csv(x = geneIDs, file = "Analysis_tools/geneIDs.csv")
```

##### Quality controls

Extract mitochondrial gene list and calculate percentage of mitochondrial genes

```
MT_genes <- subset(Biomart_annotated,
                   chromosome_name == "MT")$ensembl_gene_id

Sample_list <- lapply(X= Sample_list, FUN = PercentageFeatureSet, 
                   features = MT_genes,
                   col.name = "Percent.mito")
```

Pre-filter cells containing very few features & counts

```
Sample_list <- lapply(X =  Sample_list, FUN = subset,
                   nCount_RNA > 100 & nFeature_RNA > 200)
```

Optional merging to represent samples together for quality controls

```
Temp_merge <- merge(x = Sample_list[[1]],
                    y = c(Sample_list[[2]], Sample_list[[3]], Sample_list[[4]]),
                    add.cell.ids = Sample_names)
png("Quality_Control/Vln_Counts.png")
VlnPlot(Temp_merge, 
        features = "nCount_RNA") #  ,y.max = 40000 to see low count cells
dev.off()
```

```
## png 
##   2
```

```
png("Quality_Control/Vln_nFeature.png")
VlnPlot(Temp_merge, 
        features = "nFeature_RNA")
dev.off()
```

```
## png 
##   2
```

```
png("Quality_Control/Vln_Percent_mito.png")
VlnPlot(Temp_merge, 
        features = "Percent.mito") #  ,y.max = 50 to see low %mito cells
dev.off()
```

```
## png 
##   2
```

```
rm(Temp_merge)
```

Choose cutoff for features, count minimmums & counts maximum

```
nFeatures_cutoff <- c(1500, 1500, 1800, 1500)
nCounts_cutoff <- c(2350, 2600, 2700, 2700)
nCounts_upper <- c(48000, 54000, 48000, 54000)
```

Fig. S3-A Violin Plots of Counts, Features and %mito for sample quality controls and optional additional representations to visualize thresholds

```
for (i in 1:4) {
  png(paste0("Quality_Control/FigS4A_Vln_CTM_",Sample_names[i], ".png"), 
      height = 500, width = 500)
  print(VlnPlot(Sample_list[[i]], 
                features = c("nCount_RNA", "nFeature_RNA", "Percent.mito")))
  dev.off()
  
  # # Optional separate violin plots with thresholds represented
  
  # png(paste0("Quality_Control/Vln_Counts_",Sample_names[i], ".png"), 
  #     height = 500, width = 400 )
  # print(VlnPlot(Sample_list[[i]], 
  #               features = "nCount_RNA") +
  #         geom_hline(yintercept = nCounts_cutoff[i]) +
  #         geom_hline(yintercept = nCounts_upper[i]))
  # dev.off()
  # 
  # png(paste0("Quality_Control/Vln_Features_",Sample_names[i], ".png"), 
  #     height = 500, width = 400 )
  # print(VlnPlot(Sample_list[[i]], 
  #               features = "nFeature_RNA"))+
  #   geom_hline(yintercept = nFeatures_cutoff[i])
  # dev.off()
  # 
  # png(paste0("Quality_Control/Vln_mito_",Sample_names[i], ".png"), 
  #     height = 500, width = 400 )
  # print(VlnPlot(Sample_list[[i]], 
  #               features = "Percent.mito")+
  #         geom_hline(yintercept = 10))
  # dev.off()
  
  # # Optional histogram to help with threshold choice
  # png(paste0("Quality_Control/hist_Counts_",Sample_names[i], ".png"), 
  #     height = 500, width = 400 )
  # hist(Sample_list[[i]]@meta.data[["nCount_RNA"]], breaks = 150, xlim = c(0, nCounts_upper[i]))
  # abline(v = nCounts_cutoff[i], col = "red")
  # dev.off()
  

  # # Optional histogram to help with threshold choice
  # png(paste0("Quality_Control/hist_Features_",Sample_names[i], ".png"), 
  #     height = 500, width = 400 )
  # hist(Sample_list[[i]]@meta.data[["nFeature_RNA"]], breaks = 150, xlim = c(0, 8000) )
  # abline(v = nFeatures_cutoff[i], col = "red")
  # dev.off()
  
#  #Optional DotPlot to visualize Counts, Features and %mito and cutoffs
#   png(
#     paste0("Quality_Control/Quality_dotplot", Sample_names[i], ".png"),
#     height = 500,
#     width = 400
#   )
#   print(
#     ggplot(
#       Sample_list[[i]]@meta.data,
#       aes(x = nFeature_RNA, y = nCount_RNA, colour = Percent.mito)
#     ) +
#       geom_point() +
#       scale_color_gradient2(
#         low = "green",
#         mid = "yellow",
#         high = "red",
#         midpoint = 20
#       ) +
#       geom_hline(yintercept = nCounts_cutoff[i]) +
#       geom_hline(yintercept = nCounts_upper[i]) +
#       geom_vline(xintercept = nFeatures_cutoff[i]) +
#     scale_y_log10() )
# dev.off()
}
```

Subsetting cells that meet Feature, Genes, and mitochondrial content criterias.

In this analysis, we retained only cells containing <10% of mitochondrial genes.

```
Sample_list <- mapply(
  FUN = function(object, nC, nF, nC_up) {
    return(subset(
      object, (nCount_RNA > nC &
                 nFeature_RNA > nF &
                 Percent.mito < 10 &
                 nCount_RNA < nC_up)))},
  object = Sample_list,
  nC = nCounts_cutoff,
  nF = nFeatures_cutoff,
  nC_up = nCounts_upper)
```

Useful metadata addition & object save

```
Sample_list <- lapply(X = 1:4, FUN = function(Spl_nb){
  Sample_list[[Spl_nb]]$id_sequence <- colnames(Sample_list[[Spl_nb]])
  ID <- paste0(colnames(Sample_list[[Spl_nb]]), "_", Spl_nb)
  names(ID) <- colnames(Sample_list[[Spl_nb]])
  Sample_list[[Spl_nb]]$id <- ID
  return(Sample_list[[Spl_nb]])
})
```

Prepare integration using SCTransform function (For workflow tutorial see https://satijalab.org/seurat/articles/sctransform\_v2\_vignette.html). In this analysis, we used 4000 variable features to normalize UMIs We also used 4000 variables for integration

```
Sample_list <- lapply(X = Sample_list, 
                   FUN = SCTransform,
                   variable.features.n = 4000,
                   return.only.var.genes = F)
```

```
## Calculating cell attributes from input UMI matrix: log_umi
```

```
## Variance stabilizing transformation of count matrix of size 18198 by 5525
```

```
## Model formula is y ~ log_umi
```

```
## Get Negative Binomial regression parameters per gene
```

```
## Using 2000 genes, 5000 cells
```

```
## 
  |                                                                            
  |                                                                      |   0%
  |                                                                            
  |==================                                                    |  25%
  |                                                                            
  |===================================                                   |  50%
  |                                                                            
  |====================================================                  |  75%
  |                                                                            
  |======================================================================| 100%
```

```
## Found 84 outliers - those will be ignored in fitting/regularization step
```

```
## Second step: Get residuals using fitted parameters for 18198 genes
```

```
## 
  |                                                                            
  |                                                                      |   0%
  |                                                                            
  |==                                                                    |   3%
  |                                                                            
  |====                                                                  |   5%
  |                                                                            
  |======                                                                |   8%
  |                                                                            
  |========                                                              |  11%
  |                                                                            
  |=========                                                             |  14%
  |                                                                            
  |===========                                                           |  16%
  |                                                                            
  |=============                                                         |  19%
  |                                                                            
  |===============                                                       |  22%
  |                                                                            
  |=================                                                     |  24%
  |                                                                            
  |===================                                                   |  27%
  |                                                                            
  |=====================                                                 |  30%
  |                                                                            
  |=======================                                               |  32%
  |                                                                            
  |=========================                                             |  35%
  |                                                                            
  |==========================                                            |  38%
  |                                                                            
  |============================                                          |  41%
  |                                                                            
  |==============================                                        |  43%
  |                                                                            
  |================================                                      |  46%
  |                                                                            
  |==================================                                    |  49%
  |                                                                            
  |====================================                                  |  51%
  |                                                                            
  |======================================                                |  54%
  |                                                                            
  |========================================                              |  57%
  |                                                                            
  |==========================================                            |  59%
  |                                                                            
  |============================================                          |  62%
  |                                                                            
  |=============================================                         |  65%
  |                                                                            
  |===============================================                       |  68%
  |                                                                            
  |=================================================                     |  70%
  |                                                                            
  |===================================================                   |  73%
  |                                                                            
  |=====================================================                 |  76%
  |                                                                            
  |=======================================================               |  78%
  |                                                                            
  |=========================================================             |  81%
  |                                                                            
  |===========================================================           |  84%
  |                                                                            
  |=============================================================         |  86%
  |                                                                            
  |==============================================================        |  89%
  |                                                                            
  |================================================================      |  92%
  |                                                                            
  |==================================================================    |  95%
  |                                                                            
  |====================================================================  |  97%
  |                                                                            
  |======================================================================| 100%
```

```
## Computing corrected count matrix for 18198 genes
```

```
## 
  |                                                                            
  |                                                                      |   0%
  |                                                                            
  |==                                                                    |   3%
  |                                                                            
  |====                                                                  |   5%
  |                                                                            
  |======                                                                |   8%
  |                                                                            
  |========                                                              |  11%
  |                                                                            
  |=========                                                             |  14%
  |                                                                            
  |===========                                                           |  16%
  |                                                                            
  |=============                                                         |  19%
  |                                                                            
  |===============                                                       |  22%
  |                                                                            
  |=================                                                     |  24%
  |                                                                            
  |===================                                                   |  27%
  |                                                                            
  |=====================                                                 |  30%
  |                                                                            
  |=======================                                               |  32%
  |                                                                            
  |=========================                                             |  35%
  |                                                                            
  |==========================                                            |  38%
  |                                                                            
  |============================                                          |  41%
  |                                                                            
  |==============================                                        |  43%
  |                                                                            
  |================================                                      |  46%
  |                                                                            
  |==================================                                    |  49%
  |                                                                            
  |====================================                                  |  51%
  |                                                                            
  |======================================                                |  54%
  |                                                                            
  |========================================                              |  57%
  |                                                                            
  |==========================================                            |  59%
  |                                                                            
  |============================================                          |  62%
  |                                                                            
  |=============================================                         |  65%
  |                                                                            
  |===============================================                       |  68%
  |                                                                            
  |=================================================                     |  70%
  |                                                                            
  |===================================================                   |  73%
  |                                                                            
  |=====================================================                 |  76%
  |                                                                            
  |=======================================================               |  78%
  |                                                                            
  |=========================================================             |  81%
  |                                                                            
  |===========================================================           |  84%
  |                                                                            
  |=============================================================         |  86%
  |                                                                            
  |==============================================================        |  89%
  |                                                                            
  |================================================================      |  92%
  |                                                                            
  |==================================================================    |  95%
  |                                                                            
  |====================================================================  |  97%
  |                                                                            
  |======================================================================| 100%
```

```
## Calculating gene attributes
```

```
## Wall clock passed: Time difference of 4.006198 mins
```

```
## Determine variable features
```

```
## Place corrected count matrix in counts slot
```

```
## Centering data matrix
```

```
## Set default assay to SCT
```

```
## Calculating cell attributes from input UMI matrix: log_umi
```

```
## Variance stabilizing transformation of count matrix of size 18399 by 6004
```

```
## Model formula is y ~ log_umi
```

```
## Get Negative Binomial regression parameters per gene
```

```
## Using 2000 genes, 5000 cells
```

```
## 
  |                                                                            
  |                                                                      |   0%
  |                                                                            
  |==================                                                    |  25%
  |                                                                            
  |===================================                                   |  50%
  |                                                                            
  |====================================================                  |  75%
  |                                                                            
  |======================================================================| 100%
```

```
## Found 84 outliers - those will be ignored in fitting/regularization step
```

```
## Second step: Get residuals using fitted parameters for 18399 genes
```

```
## 
  |                                                                            
  |                                                                      |   0%
  |                                                                            
  |==                                                                    |   3%
  |                                                                            
  |====                                                                  |   5%
  |                                                                            
  |======                                                                |   8%
  |                                                                            
  |========                                                              |  11%
  |                                                                            
  |=========                                                             |  14%
  |                                                                            
  |===========                                                           |  16%
  |                                                                            
  |=============                                                         |  19%
  |                                                                            
  |===============                                                       |  22%
  |                                                                            
  |=================                                                     |  24%
  |                                                                            
  |===================                                                   |  27%
  |                                                                            
  |=====================                                                 |  30%
  |                                                                            
  |=======================                                               |  32%
  |                                                                            
  |=========================                                             |  35%
  |                                                                            
  |==========================                                            |  38%
  |                                                                            
  |============================                                          |  41%
  |                                                                            
  |==============================                                        |  43%
  |                                                                            
  |================================                                      |  46%
  |                                                                            
  |==================================                                    |  49%
  |                                                                            
  |====================================                                  |  51%
  |                                                                            
  |======================================                                |  54%
  |                                                                            
  |========================================                              |  57%
  |                                                                            
  |==========================================                            |  59%
  |                                                                            
  |============================================                          |  62%
  |                                                                            
  |=============================================                         |  65%
  |                                                                            
  |===============================================                       |  68%
  |                                                                            
  |=================================================                     |  70%
  |                                                                            
  |===================================================                   |  73%
  |                                                                            
  |=====================================================                 |  76%
  |                                                                            
  |=======================================================               |  78%
  |                                                                            
  |=========================================================             |  81%
  |                                                                            
  |===========================================================           |  84%
  |                                                                            
  |=============================================================         |  86%
  |                                                                            
  |==============================================================        |  89%
  |                                                                            
  |================================================================      |  92%
  |                                                                            
  |==================================================================    |  95%
  |                                                                            
  |====================================================================  |  97%
  |                                                                            
  |======================================================================| 100%
```

```
## Computing corrected count matrix for 18399 genes
```

```
## 
  |                                                                            
  |                                                                      |   0%
  |                                                                            
  |==                                                                    |   3%
  |                                                                            
  |====                                                                  |   5%
  |                                                                            
  |======                                                                |   8%
  |                                                                            
  |========                                                              |  11%
  |                                                                            
  |=========                                                             |  14%
  |                                                                            
  |===========                                                           |  16%
  |                                                                            
  |=============                                                         |  19%
  |                                                                            
  |===============                                                       |  22%
  |                                                                            
  |=================                                                     |  24%
  |                                                                            
  |===================                                                   |  27%
  |                                                                            
  |=====================                                                 |  30%
  |                                                                            
  |=======================                                               |  32%
  |                                                                            
  |=========================                                             |  35%
  |                                                                            
  |==========================                                            |  38%
  |                                                                            
  |============================                                          |  41%
  |                                                                            
  |==============================                                        |  43%
  |                                                                            
  |================================                                      |  46%
  |                                                                            
  |==================================                                    |  49%
  |                                                                            
  |====================================                                  |  51%
  |                                                                            
  |======================================                                |  54%
  |                                                                            
  |========================================                              |  57%
  |                                                                            
  |==========================================                            |  59%
  |                                                                            
  |============================================                          |  62%
  |                                                                            
  |=============================================                         |  65%
  |                                                                            
  |===============================================                       |  68%
  |                                                                            
  |=================================================                     |  70%
  |                                                                            
  |===================================================                   |  73%
  |                                                                            
  |=====================================================                 |  76%
  |                                                                            
  |=======================================================               |  78%
  |                                                                            
  |=========================================================             |  81%
  |                                                                            
  |===========================================================           |  84%
  |                                                                            
  |=============================================================         |  86%
  |                                                                            
  |==============================================================        |  89%
  |                                                                            
  |================================================================      |  92%
  |                                                                            
  |==================================================================    |  95%
  |                                                                            
  |====================================================================  |  97%
  |                                                                            
  |======================================================================| 100%
```

```
## Calculating gene attributes
```

```
## Wall clock passed: Time difference of 2.900929 mins
```

```
## Determine variable features
```

```
## Place corrected count matrix in counts slot
```

```
## Centering data matrix
```

```
## Set default assay to SCT
```

```
## Calculating cell attributes from input UMI matrix: log_umi
```

```
## Variance stabilizing transformation of count matrix of size 17534 by 2739
```

```
## Model formula is y ~ log_umi
```

```
## Get Negative Binomial regression parameters per gene
```

```
## Using 2000 genes, 2739 cells
```

```
## 
  |                                                                            
  |                                                                      |   0%
  |                                                                            
  |==================                                                    |  25%
  |                                                                            
  |===================================                                   |  50%
  |                                                                            
  |====================================================                  |  75%
  |                                                                            
  |======================================================================| 100%
```

```
## Found 70 outliers - those will be ignored in fitting/regularization step
```

```
## Second step: Get residuals using fitted parameters for 17534 genes
```

```
## 
  |                                                                            
  |                                                                      |   0%
  |                                                                            
  |==                                                                    |   3%
  |                                                                            
  |====                                                                  |   6%
  |                                                                            
  |======                                                                |   8%
  |                                                                            
  |========                                                              |  11%
  |                                                                            
  |==========                                                            |  14%
  |                                                                            
  |============                                                          |  17%
  |                                                                            
  |==============                                                        |  19%
  |                                                                            
  |================                                                      |  22%
  |                                                                            
  |==================                                                    |  25%
  |                                                                            
  |===================                                                   |  28%
  |                                                                            
  |=====================                                                 |  31%
  |                                                                            
  |=======================                                               |  33%
  |                                                                            
  |=========================                                             |  36%
  |                                                                            
  |===========================                                           |  39%
  |                                                                            
  |=============================                                         |  42%
  |                                                                            
  |===============================                                       |  44%
  |                                                                            
  |=================================                                     |  47%
  |                                                                            
  |===================================                                   |  50%
  |                                                                            
  |=====================================                                 |  53%
  |                                                                            
  |=======================================                               |  56%
  |                                                                            
  |=========================================                             |  58%
  |                                                                            
  |===========================================                           |  61%
  |                                                                            
  |=============================================                         |  64%
  |                                                                            
  |===============================================                       |  67%
  |                                                                            
  |=================================================                     |  69%
  |                                                                            
  |===================================================                   |  72%
  |                                                                            
  |====================================================                  |  75%
  |                                                                            
  |======================================================                |  78%
  |                                                                            
  |========================================================              |  81%
  |                                                                            
  |==========================================================            |  83%
  |                                                                            
  |============================================================          |  86%
  |                                                                            
  |==============================================================        |  89%
  |                                                                            
  |================================================================      |  92%
  |                                                                            
  |==================================================================    |  94%
  |                                                                            
  |====================================================================  |  97%
  |                                                                            
  |======================================================================| 100%
```

```
## Computing corrected count matrix for 17534 genes
```

```
## 
  |                                                                            
  |                                                                      |   0%
  |                                                                            
  |==                                                                    |   3%
  |                                                                            
  |====                                                                  |   6%
  |                                                                            
  |======                                                                |   8%
  |                                                                            
  |========                                                              |  11%
  |                                                                            
  |==========                                                            |  14%
  |                                                                            
  |============                                                          |  17%
  |                                                                            
  |==============                                                        |  19%
  |                                                                            
  |================                                                      |  22%
  |                                                                            
  |==================                                                    |  25%
  |                                                                            
  |===================                                                   |  28%
  |                                                                            
  |=====================                                                 |  31%
  |                                                                            
  |=======================                                               |  33%
  |                                                                            
  |=========================                                             |  36%
  |                                                                            
  |===========================                                           |  39%
  |                                                                            
  |=============================                                         |  42%
  |                                                                            
  |===============================                                       |  44%
  |                                                                            
  |=================================                                     |  47%
  |                                                                            
  |===================================                                   |  50%
  |                                                                            
  |=====================================                                 |  53%
  |                                                                            
  |=======================================                               |  56%
  |                                                                            
  |=========================================                             |  58%
  |                                                                            
  |===========================================                           |  61%
  |                                                                            
  |=============================================                         |  64%
  |                                                                            
  |===============================================                       |  67%
  |                                                                            
  |=================================================                     |  69%
  |                                                                            
  |===================================================                   |  72%
  |                                                                            
  |====================================================                  |  75%
  |                                                                            
  |======================================================                |  78%
  |                                                                            
  |========================================================              |  81%
  |                                                                            
  |==========================================================            |  83%
  |                                                                            
  |============================================================          |  86%
  |                                                                            
  |==============================================================        |  89%
  |                                                                            
  |================================================================      |  92%
  |                                                                            
  |==================================================================    |  94%
  |                                                                            
  |====================================================================  |  97%
  |                                                                            
  |======================================================================| 100%
```

```
## Calculating gene attributes
```

```
## Wall clock passed: Time difference of 43.99547 secs
```

```
## Determine variable features
```

```
## Place corrected count matrix in counts slot
```

```
## Centering data matrix
```

```
## Set default assay to SCT
```

```
## Calculating cell attributes from input UMI matrix: log_umi
```

```
## Variance stabilizing transformation of count matrix of size 17811 by 3075
```

```
## Model formula is y ~ log_umi
```

```
## Get Negative Binomial regression parameters per gene
```

```
## Using 2000 genes, 3075 cells
```

```
## 
  |                                                                            
  |                                                                      |   0%
  |                                                                            
  |==================                                                    |  25%
  |                                                                            
  |===================================                                   |  50%
  |                                                                            
  |====================================================                  |  75%
  |                                                                            
  |======================================================================| 100%
```

```
## Found 72 outliers - those will be ignored in fitting/regularization step
```

```
## Second step: Get residuals using fitted parameters for 17811 genes
```

```
## 
  |                                                                            
  |                                                                      |   0%
  |                                                                            
  |==                                                                    |   3%
  |                                                                            
  |====                                                                  |   6%
  |                                                                            
  |======                                                                |   8%
  |                                                                            
  |========                                                              |  11%
  |                                                                            
  |==========                                                            |  14%
  |                                                                            
  |============                                                          |  17%
  |                                                                            
  |==============                                                        |  19%
  |                                                                            
  |================                                                      |  22%
  |                                                                            
  |==================                                                    |  25%
  |                                                                            
  |===================                                                   |  28%
  |                                                                            
  |=====================                                                 |  31%
  |                                                                            
  |=======================                                               |  33%
  |                                                                            
  |=========================                                             |  36%
  |                                                                            
  |===========================                                           |  39%
  |                                                                            
  |=============================                                         |  42%
  |                                                                            
  |===============================                                       |  44%
  |                                                                            
  |=================================                                     |  47%
  |                                                                            
  |===================================                                   |  50%
  |                                                                            
  |=====================================                                 |  53%
  |                                                                            
  |=======================================                               |  56%
  |                                                                            
  |=========================================                             |  58%
  |                                                                            
  |===========================================                           |  61%
  |                                                                            
  |=============================================                         |  64%
  |                                                                            
  |===============================================                       |  67%
  |                                                                            
  |=================================================                     |  69%
  |                                                                            
  |===================================================                   |  72%
  |                                                                            
  |====================================================                  |  75%
  |                                                                            
  |======================================================                |  78%
  |                                                                            
  |========================================================              |  81%
  |                                                                            
  |==========================================================            |  83%
  |                                                                            
  |============================================================          |  86%
  |                                                                            
  |==============================================================        |  89%
  |                                                                            
  |================================================================      |  92%
  |                                                                            
  |==================================================================    |  94%
  |                                                                            
  |====================================================================  |  97%
  |                                                                            
  |======================================================================| 100%
```

```
## Computing corrected count matrix for 17811 genes
```

```
## 
  |                                                                            
  |                                                                      |   0%
  |                                                                            
  |==                                                                    |   3%
  |                                                                            
  |====                                                                  |   6%
  |                                                                            
  |======                                                                |   8%
  |                                                                            
  |========                                                              |  11%
  |                                                                            
  |==========                                                            |  14%
  |                                                                            
  |============                                                          |  17%
  |                                                                            
  |==============                                                        |  19%
  |                                                                            
  |================                                                      |  22%
  |                                                                            
  |==================                                                    |  25%
  |                                                                            
  |===================                                                   |  28%
  |                                                                            
  |=====================                                                 |  31%
  |                                                                            
  |=======================                                               |  33%
  |                                                                            
  |=========================                                             |  36%
  |                                                                            
  |===========================                                           |  39%
  |                                                                            
  |=============================                                         |  42%
  |                                                                            
  |===============================                                       |  44%
  |                                                                            
  |=================================                                     |  47%
  |                                                                            
  |===================================                                   |  50%
  |                                                                            
  |=====================================                                 |  53%
  |                                                                            
  |=======================================                               |  56%
  |                                                                            
  |=========================================                             |  58%
  |                                                                            
  |===========================================                           |  61%
  |                                                                            
  |=============================================                         |  64%
  |                                                                            
  |===============================================                       |  67%
  |                                                                            
  |=================================================                     |  69%
  |                                                                            
  |===================================================                   |  72%
  |                                                                            
  |====================================================                  |  75%
  |                                                                            
  |======================================================                |  78%
  |                                                                            
  |========================================================              |  81%
  |                                                                            
  |==========================================================            |  83%
  |                                                                            
  |============================================================          |  86%
  |                                                                            
  |==============================================================        |  89%
  |                                                                            
  |================================================================      |  92%
  |                                                                            
  |==================================================================    |  94%
  |                                                                            
  |====================================================================  |  97%
  |                                                                            
  |======================================================================| 100%
```

```
## Calculating gene attributes
```

```
## Wall clock passed: Time difference of 48.77429 secs
```

```
## Determine variable features
```

```
## Place corrected count matrix in counts slot
```

```
## Centering data matrix
```

```
## Set default assay to SCT
```

We can then proceed to finding integration features based on the 4000 variable features, and carry on with the standard SCT integration workflow Note that this results in an integrated assay, but that differential expression analysis will require an additional normalization step, using the PrepSCTIntegration function.

```
Integration_Features <- SelectIntegrationFeatures(object.list = Sample_list, 
                                                  nfeatures = 4000,
                                                  assay = rep("SCT",4))

Sample_list <- PrepSCTIntegration(object.list = Sample_list,
                               anchor.features = Integration_Features)

Sample_list <- 
  FindIntegrationAnchors(Sample_list, normalization.method = "SCT",
                         anchor.features = Integration_Features)
```

```
## Warning in CheckDuplicateCellNames(object.list = object.list): Some cell names
## are duplicated across objects provided. Renaming to enforce unique cell names.
```

```
## Finding all pairwise anchors
```

```
## Running CCA
```

```
## Merging objects
```

```
## Finding neighborhoods
```

```
## Finding anchors
```

```
##  Found 16075 anchors
```

```
## Filtering anchors
```

```
##  Retained 13939 anchors
```

```
## Running CCA
```

```
## Merging objects
```

```
## Finding neighborhoods
```

```
## Finding anchors
```

```
##  Found 8265 anchors
```

```
## Filtering anchors
```

```
##  Retained 7568 anchors
```

```
## Running CCA
```

```
## Merging objects
```

```
## Finding neighborhoods
```

```
## Finding anchors
```

```
##  Found 8654 anchors
```

```
## Filtering anchors
```

```
##  Retained 7992 anchors
```

```
## Running CCA
```

```
## Merging objects
```

```
## Finding neighborhoods
```

```
## Finding anchors
```

```
##  Found 8516 anchors
```

```
## Filtering anchors
```

```
##  Retained 7743 anchors
```

```
## Running CCA
```

```
## Merging objects
```

```
## Finding neighborhoods
```

```
## Finding anchors
```

```
##  Found 8844 anchors
```

```
## Filtering anchors
```

```
##  Retained 7919 anchors
```

```
## Running CCA
```

```
## Merging objects
```

```
## Finding neighborhoods
```

```
## Finding anchors
```

```
##  Found 8791 anchors
```

```
## Filtering anchors
```

```
##  Retained 8231 anchors
```

```
SCT_obj <- IntegrateData(anchorset = Sample_list, normalization.method = "SCT")
```

```
## Merging dataset 3 into 4
```

```
## Extracting anchors for merged samples
```

```
## Finding integration vectors
```

```
## Finding integration vector weights
```

```
## Integrating data
```

```
## Merging dataset 4 3 into 2
```

```
## Warning: Attempting to merge an SCTAssay with another Assay type 
## Converting all to standard Assay objects.
```

```
## Extracting anchors for merged samples
```

```
## Warning: Attempting to merge an SCTAssay with another Assay type 
## Converting all to standard Assay objects.
```

```
## Finding integration vectors
```

```
## Finding integration vector weights
```

```
## Integrating data
```

```
## Merging dataset 1 into 2 4 3
```

```
## Extracting anchors for merged samples
```

```
## Finding integration vectors
```

```
## Finding integration vector weights
```

```
## Integrating data
```

```
saveRDS(object = SCT_obj, file = "Objects/SCT_obj.rds")
```

```
sessionInfo()
```

```
## R version 4.1.0 (2021-05-18)
## Platform: x86_64-conda-linux-gnu (64-bit)
## Running under: Ubuntu 20.04.4 LTS
## 
## Matrix products: default
## BLAS/LAPACK: /opt/rstudio-server_conda/conda/envs/rstudio-server_4.1.0/lib/libopenblasp-r0.3.15.so
## 
## locale:
##  [1] LC_CTYPE=en_US.UTF-8       LC_NUMERIC=C              
##  [3] LC_TIME=en_US.UTF-8        LC_COLLATE=en_US.UTF-8    
##  [5] LC_MONETARY=en_US.UTF-8    LC_MESSAGES=en_US.UTF-8   
##  [7] LC_PAPER=en_US.UTF-8       LC_NAME=C                 
##  [9] LC_ADDRESS=C               LC_TELEPHONE=C            
## [11] LC_MEASUREMENT=en_US.UTF-8 LC_IDENTIFICATION=C       
## 
## attached base packages:
## [1] stats     graphics  grDevices utils     datasets  methods   base     
## 
## other attached packages:
##  [1] patchwork_1.1.2    biomaRt_2.48.3     forcats_0.5.2      stringr_1.5.0     
##  [5] dplyr_1.0.10       purrr_1.0.1        readr_2.1.2        tidyr_1.2.0       
##  [9] tibble_3.1.7       ggplot2_3.4.0      tidyverse_1.3.2    remotes_2.4.2     
## [13] sp_1.6-0           SeuratObject_4.1.0 Seurat_4.1.1      
## 
## loaded via a namespace (and not attached):
##   [1] utf8_1.2.2             reticulate_1.27        tidyselect_1.2.0      
##   [4] RSQLite_2.2.14         AnnotationDbi_1.54.1   htmlwidgets_1.5.4     
##   [7] grid_4.1.0             Rtsne_0.16             munsell_0.5.0         
##  [10] codetools_0.2-18       ica_1.0-2              future_1.30.0         
##  [13] miniUI_0.1.1.1         withr_2.5.0            spatstat.random_3.0-1 
##  [16] colorspace_2.0-3       progressr_0.13.0       Biobase_2.52.0        
##  [19] filelock_1.0.2         knitr_1.39             rstudioapi_0.13       
##  [22] stats4_4.1.0           ROCR_1.0-11            tensor_1.5            
##  [25] listenv_0.8.0          labeling_0.4.2         GenomeInfoDbData_1.2.6
##  [28] polyclip_1.10-0        farver_2.1.0           bit64_4.0.5           
##  [31] parallelly_1.34.0      vctrs_0.5.1            generics_0.1.3        
##  [34] xfun_0.31              BiocFileCache_2.0.0    R6_2.5.1              
##  [37] GenomeInfoDb_1.28.4    bitops_1.0-7           spatstat.utils_3.0-1  
##  [40] cachem_1.0.6           assertthat_0.2.1       promises_1.2.0.1      
##  [43] scales_1.2.0           vroom_1.5.7            googlesheets4_1.0.0   
##  [46] rgeos_0.5-9            gtable_0.3.0           globals_0.16.2        
##  [49] goftest_1.2-3          rlang_1.0.6            splines_4.1.0         
##  [52] lazyeval_0.2.2         gargle_1.2.0           spatstat.geom_3.0-3   
##  [55] broom_1.0.2            yaml_2.3.5             reshape2_1.4.4        
##  [58] abind_1.4-5            modelr_0.1.8           backports_1.4.1       
##  [61] httpuv_1.6.5           tools_4.1.0            ellipsis_0.3.2        
##  [64] spatstat.core_2.4-4    jquerylib_0.1.4        RColorBrewer_1.1-3    
##  [67] BiocGenerics_0.38.0    ggridges_0.5.3         Rcpp_1.0.8.3          
##  [70] plyr_1.8.7             progress_1.2.2         zlibbioc_1.38.0       
##  [73] RCurl_1.98-1.6         prettyunits_1.1.1      rpart_4.1.16          
##  [76] deldir_1.0-6           pbapply_1.5-0          cowplot_1.1.1         
##  [79] S4Vectors_0.30.2       zoo_1.8-10             haven_2.5.0           
##  [82] ggrepel_0.9.1          cluster_2.1.3          fs_1.5.2              
##  [85] magrittr_2.0.3         data.table_1.14.2      scattermore_0.8       
##  [88] lmtest_0.9-40          reprex_2.0.1           RANN_2.6.1            
##  [91] googledrive_2.0.0      fitdistrplus_1.1-8     matrixStats_0.62.0    
##  [94] hms_1.1.1              mime_0.12              evaluate_0.15         
##  [97] xtable_1.8-4           XML_3.99-0.9           readxl_1.4.1          
## [100] IRanges_2.26.0         gridExtra_2.3          compiler_4.1.0        
## [103] KernSmooth_2.23-20     crayon_1.5.1           htmltools_0.5.2       
## [106] mgcv_1.8-40            later_1.3.0            tzdb_0.3.0            
## [109] lubridate_1.8.0        DBI_1.1.2              dbplyr_2.3.0          
## [112] MASS_7.3-57            rappdirs_0.3.3         Matrix_1.5-3          
## [115] cli_3.6.0              parallel_4.1.0         igraph_1.3.5          
## [118] pkgconfig_2.0.3        plotly_4.10.1          spatstat.sparse_3.0-0 
## [121] xml2_1.3.3             bslib_0.3.1            XVector_0.32.0        
## [124] rvest_1.0.2            digest_0.6.29          sctransform_0.3.5     
## [127] RcppAnnoy_0.0.19       spatstat.data_3.0-0    Biostrings_2.60.2     
## [130] rmarkdown_2.14         cellranger_1.1.0       leiden_0.4.2          
## [133] uwot_0.1.14            curl_4.3.2             shiny_1.7.1           
## [136] lifecycle_1.0.3        nlme_3.1-157           jsonlite_1.8.0        
## [139] viridisLite_0.4.0      fansi_1.0.3            pillar_1.7.0          
## [142] lattice_0.20-45        KEGGREST_1.32.0        fastmap_1.1.0         
## [145] httr_1.4.3             survival_3.3-1         glue_1.6.2            
## [148] png_0.1-7              bit_4.0.4              stringi_1.7.6         
## [151] sass_0.4.1             blob_1.2.3             memoise_2.0.1         
## [154] irlba_2.3.5            future.apply_1.9.0
```
