## Supplementary material for "Switching of RNA splicing regulators in immature neuroblasts: a key step in adult neurogenesis": script_R_3: 3_UMAP_clustering_resolution_choice.html

Cell cycle scoring and UMAP generation


### Cell cycle scoring and UMAP generation

In this script, we use the CellCycle Scoring function of Seurat, generate a UMAP and cluster cells together.

```
library(tidyverse)
```

```
## ── Attaching packages ─────────────────────────────────────── tidyverse 1.3.2 ──
## ✔ ggplot2 3.4.0      ✔ purrr   0.3.5 
## ✔ tibble  3.1.8      ✔ dplyr   1.0.10
## ✔ tidyr   1.2.1      ✔ stringr 1.4.1 
## ✔ readr   2.1.2      ✔ forcats 0.5.2 
## ── Conflicts ────────────────────────────────────────── tidyverse_conflicts() ──
## ✖ dplyr::filter() masks stats::filter()
## ✖ dplyr::lag()    masks stats::lag()
```

```
library(dplyr)
library(clustree)
```

```
## Loading required package: ggraph
```

```
library(patchwork)
library(Seurat, lib.loc = "/opt/rstudio-server_conda/conda/envs/rstudio-server_4.1.0/lib/R/library/")
```

```
## Attaching SeuratObject
## Attaching sp
```

Import objects

```
SCT_obj <- readRDS("Objects/SCT_obj.rds")
Biomart_annotated <- read.csv("Analysis_tools/Biomart_annotated.csv")
```

The cell cycle scoring feature of seurat affects a S and G2/M score to each cell. For basic classification, cell cycle status can be inferred from the difference between S and G2M score

```
DefaultAssay(SCT_obj) <- "SCT"

#Fetch genes associated to the cell cycle phases, returns a 2-items list
cc.genes.mouse <- lapply(X = cc.genes.updated.2019, FUN = str_to_sentence) 

#Select genes present in assay
cc.mouse.ok <- lapply(X = cc.genes.mouse, function(gene_vector){ 
  return(Biomart_annotated[Biomart_annotated$external_gene_name %in% gene_vector,1])})

SCT_obj <- CellCycleScoring(SCT_obj, 
                            s.features = cc.mouse.ok$s.genes,
                            g2m.features = cc.mouse.ok$g2m.genes)

# A Diff score can be used to regress the cell cycle
SCT_obj$diff_score <- SCT_obj$S.Score - SCT_obj$G2M.Score
```

Run PCA and assess explained variance using ElbowPlot

```
DefaultAssay(SCT_obj) <- "integrated"
SCT_obj <- RunPCA(object = SCT_obj)
```

```
## PC_ 1 
## Positive:  ENSMUSG00000028832, ENSMUSG00000072235, ENSMUSG00000000184, ENSMUSG00000029838, ENSMUSG00000079523, ENSMUSG00000045136, ENSMUSG00000063632, ENSMUSG00000027500, ENSMUSG00000001525, ENSMUSG00000005089 
##     ENSMUSG00000003038, ENSMUSG00000007097, ENSMUSG00000068748, ENSMUSG00000054717, ENSMUSG00000028517, ENSMUSG00000062380, ENSMUSG00000021268, ENSMUSG00000027210, ENSMUSG00000076431, ENSMUSG00000029309 
##     ENSMUSG00000060743, ENSMUSG00000031342, ENSMUSG00000020914, ENSMUSG00000040204, ENSMUSG00000090063, ENSMUSG00000008575, ENSMUSG00000027805, ENSMUSG00000032482, ENSMUSG00000079037, ENSMUSG00000037852 
## Negative:  ENSMUSG00000036887, ENSMUSG00000021665, ENSMUSG00000036905, ENSMUSG00000036896, ENSMUSG00000052336, ENSMUSG00000038642, ENSMUSG00000007891, ENSMUSG00000027447, ENSMUSG00000024621, ENSMUSG00000021190 
##     ENSMUSG00000023992, ENSMUSG00000036353, ENSMUSG00000018593, ENSMUSG00000026628, ENSMUSG00000048163, ENSMUSG00000052837, ENSMUSG00000040229, ENSMUSG00000052684, ENSMUSG00000044786, ENSMUSG00000058715 
##     ENSMUSG00000030579, ENSMUSG00000028581, ENSMUSG00000015852, ENSMUSG00000054675, ENSMUSG00000021250, ENSMUSG00000020423, ENSMUSG00000021423, ENSMUSG00000051504, ENSMUSG00000056708, ENSMUSG00000027848 
## PC_ 2 
## Positive:  ENSMUSG00000028832, ENSMUSG00000072235, ENSMUSG00000000184, ENSMUSG00000054717, ENSMUSG00000063632, ENSMUSG00000027500, ENSMUSG00000001525, ENSMUSG00000079523, ENSMUSG00000003038, ENSMUSG00000076431 
##     ENSMUSG00000045136, ENSMUSG00000020914, ENSMUSG00000062380, ENSMUSG00000040204, ENSMUSG00000090063, ENSMUSG00000058773, ENSMUSG00000094777, ENSMUSG00000037894, ENSMUSG00000069272, ENSMUSG00000052727 
##     ENSMUSG00000060743, ENSMUSG00000026238, ENSMUSG00000031004, ENSMUSG00000027805, ENSMUSG00000026605, ENSMUSG00000066392, ENSMUSG00000027210, ENSMUSG00000021087, ENSMUSG00000035551, ENSMUSG00000001403 
## Negative:  ENSMUSG00000029309, ENSMUSG00000029838, ENSMUSG00000007097, ENSMUSG00000007872, ENSMUSG00000030235, ENSMUSG00000002985, ENSMUSG00000023175, ENSMUSG00000031765, ENSMUSG00000017754, ENSMUSG00000032766 
##     ENSMUSG00000036256, ENSMUSG00000029648, ENSMUSG00000028645, ENSMUSG00000004951, ENSMUSG00000020315, ENSMUSG00000024140, ENSMUSG00000041378, ENSMUSG00000058297, ENSMUSG00000031239, ENSMUSG00000079018 
##     ENSMUSG00000025492, ENSMUSG00000045092, ENSMUSG00000075602, ENSMUSG00000030237, ENSMUSG00000042116, ENSMUSG00000042745, ENSMUSG00000020154, ENSMUSG00000026921, ENSMUSG00000020044, ENSMUSG00000061353 
## PC_ 3 
## Positive:  ENSMUSG00000036256, ENSMUSG00000029648, ENSMUSG00000041378, ENSMUSG00000079018, ENSMUSG00000004951, ENSMUSG00000031239, ENSMUSG00000075602, ENSMUSG00000030237, ENSMUSG00000017754, ENSMUSG00000026921 
##     ENSMUSG00000020154, ENSMUSG00000040584, ENSMUSG00000058297, ENSMUSG00000001946, ENSMUSG00000061353, ENSMUSG00000056492, ENSMUSG00000042116, ENSMUSG00000002504, ENSMUSG00000039167, ENSMUSG00000030413 
##     ENSMUSG00000038370, ENSMUSG00000032766, ENSMUSG00000054690, ENSMUSG00000000805, ENSMUSG00000029802, ENSMUSG00000030096, ENSMUSG00000025085, ENSMUSG00000001240, ENSMUSG00000025492, ENSMUSG00000026814 
## Negative:  ENSMUSG00000027447, ENSMUSG00000028517, ENSMUSG00000032482, ENSMUSG00000005360, ENSMUSG00000005089, ENSMUSG00000022037, ENSMUSG00000031760, ENSMUSG00000002985, ENSMUSG00000050953, ENSMUSG00000026424 
##     ENSMUSG00000017390, ENSMUSG00000030428, ENSMUSG00000004892, ENSMUSG00000032883, ENSMUSG00000022132, ENSMUSG00000020591, ENSMUSG00000031765, ENSMUSG00000028128, ENSMUSG00000060961, ENSMUSG00000026701 
##     ENSMUSG00000006205, ENSMUSG00000024411, ENSMUSG00000055254, ENSMUSG00000007097, ENSMUSG00000059974, ENSMUSG00000001270, ENSMUSG00000024302, ENSMUSG00000033998, ENSMUSG00000075012, ENSMUSG00000021508 
## PC_ 4 
## Positive:  ENSMUSG00000027447, ENSMUSG00000029309, ENSMUSG00000054717, ENSMUSG00000007097, ENSMUSG00000005089, ENSMUSG00000045092, ENSMUSG00000030605, ENSMUSG00000020914, ENSMUSG00000032482, ENSMUSG00000068748 
##     ENSMUSG00000050953, ENSMUSG00000005360, ENSMUSG00000022037, ENSMUSG00000030235, ENSMUSG00000028517, ENSMUSG00000040204, ENSMUSG00000031760, ENSMUSG00000031762, ENSMUSG00000017390, ENSMUSG00000007872 
##     ENSMUSG00000026424, ENSMUSG00000004892, ENSMUSG00000058773, ENSMUSG00000020591, ENSMUSG00000022132, ENSMUSG00000031004, ENSMUSG00000048960, ENSMUSG00000041329, ENSMUSG00000026605, ENSMUSG00000060961 
## Negative:  ENSMUSG00000032554, ENSMUSG00000062591, ENSMUSG00000036634, ENSMUSG00000041607, ENSMUSG00000024661, ENSMUSG00000031425, ENSMUSG00000037625, ENSMUSG00000076439, ENSMUSG00000032517, ENSMUSG00000032060 
##     ENSMUSG00000006782, ENSMUSG00000022425, ENSMUSG00000026830, ENSMUSG00000032854, ENSMUSG00000027199, ENSMUSG00000022548, ENSMUSG00000056966, ENSMUSG00000027858, ENSMUSG00000037166, ENSMUSG00000036745 
##     ENSMUSG00000020486, ENSMUSG00000006651, ENSMUSG00000020774, ENSMUSG00000015806, ENSMUSG00000073680, ENSMUSG00000046160, ENSMUSG00000027674, ENSMUSG00000033579, ENSMUSG00000028399, ENSMUSG00000034488 
## PC_ 5 
## Positive:  ENSMUSG00000054717, ENSMUSG00000020914, ENSMUSG00000040204, ENSMUSG00000031004, ENSMUSG00000058773, ENSMUSG00000026605, ENSMUSG00000038943, ENSMUSG00000001403, ENSMUSG00000094777, ENSMUSG00000049932 
##     ENSMUSG00000069272, ENSMUSG00000028873, ENSMUSG00000027469, ENSMUSG00000027306, ENSMUSG00000017716, ENSMUSG00000062248, ENSMUSG00000034349, ENSMUSG00000022033, ENSMUSG00000029177, ENSMUSG00000037894 
##     ENSMUSG00000005233, ENSMUSG00000019942, ENSMUSG00000020649, ENSMUSG00000045328, ENSMUSG00000028312, ENSMUSG00000020330, ENSMUSG00000012443, ENSMUSG00000023505, ENSMUSG00000041431, ENSMUSG00000074476 
## Negative:  ENSMUSG00000021268, ENSMUSG00000027500, ENSMUSG00000045136, ENSMUSG00000072235, ENSMUSG00000090063, ENSMUSG00000076431, ENSMUSG00000027210, ENSMUSG00000038718, ENSMUSG00000062380, ENSMUSG00000040209 
##     ENSMUSG00000052727, ENSMUSG00000079523, ENSMUSG00000052551, ENSMUSG00000021087, ENSMUSG00000031285, ENSMUSG00000023800, ENSMUSG00000051243, ENSMUSG00000052516, ENSMUSG00000024268, ENSMUSG00000066392 
##     ENSMUSG00000052534, ENSMUSG00000035551, ENSMUSG00000042834, ENSMUSG00000029563, ENSMUSG00000016386, ENSMUSG00000008575, ENSMUSG00000067786, ENSMUSG00000056895, ENSMUSG00000061911, ENSMUSG00000041362
```

```
((PCAPlot(SCT_obj, group.by = "orig.ident")+
    ElbowPlot(SCT_obj, ndims = 50, reduction = "pca"))+
    plot_layout(guides = 'collect'))
```

UMAP is created using the first 50 PCs and clusterization is performed based on that graph.

```
DefaultAssay(SCT_obj) <- "integrated"
SCT_obj <- RunUMAP(SCT_obj,
                   dims = 1:50,
                   n.components = 2L, # 3L for downstream 3D UMAP representation, will affect 2D UMAP
                   assay = "integrated",
                   return.model = T) # Useful for downstream pseudotime applications (e.g. Monocle)
```

```
## Warning: The default method for RunUMAP has changed from calling Python UMAP via reticulate to the R-native UWOT using the cosine metric
## To use Python UMAP via reticulate, set umap.method to 'umap-learn' and metric to 'correlation'
## This message will be shown once per session
```

```
## UMAP will return its model
```

```
## 12:04:51 UMAP embedding parameters a = 0.9922 b = 1.112
```

```
## 12:04:51 Read 17343 rows and found 50 numeric columns
```

```
## 12:04:51 Using Annoy for neighbor search, n_neighbors = 30
```

```
## 12:04:51 Building Annoy index with metric = cosine, n_trees = 50
```

```
## 0%   10   20   30   40   50   60   70   80   90   100%
```

```
## [----|----|----|----|----|----|----|----|----|----|
```

```
## **************************************************|
## 12:04:54 Writing NN index file to temp file /tmp/RtmpZtkB5A/filedeafa19914492
## 12:04:54 Searching Annoy index using 1 thread, search_k = 3000
## 12:04:59 Annoy recall = 100%
## 12:05:00 Commencing smooth kNN distance calibration using 1 thread
## 12:05:01 Initializing from normalized Laplacian + noise
## 12:05:03 Commencing optimization for 200 epochs, with 713598 positive edges
## 12:05:11 Optimization finished
```

```
UMAPPlot(object = SCT_obj, group.by = "orig.ident")
```

```
SCT_obj <- FindNeighbors(SCT_obj, k.param = 20, dims = c(1:50), reduction = "pca",
                         graph.name = c("int_20_nn", "int_20_snn"))
```

```
## Computing nearest neighbor graph
## Computing SNN
```

```
resolutions <- seq(0.2,1.6, 0.2)
SCT_obj <- FindClusters(SCT_obj, 
                        graph.name = "int_20_snn", 
                        resolution = resolutions)
```

```
## Modularity Optimizer version 1.3.0 by Ludo Waltman and Nees Jan van Eck
## 
## Number of nodes: 17343
## Number of edges: 615265
## 
## Running Louvain algorithm...
## Maximum modularity in 10 random starts: 0.9641
## Number of communities: 17
## Elapsed time: 1 seconds
## Modularity Optimizer version 1.3.0 by Ludo Waltman and Nees Jan van Eck
## 
## Number of nodes: 17343
## Number of edges: 615265
## 
## Running Louvain algorithm...
## Maximum modularity in 10 random starts: 0.9404
## Number of communities: 20
## Elapsed time: 2 seconds
## Modularity Optimizer version 1.3.0 by Ludo Waltman and Nees Jan van Eck
## 
## Number of nodes: 17343
## Number of edges: 615265
## 
## Running Louvain algorithm...
## Maximum modularity in 10 random starts: 0.9210
## Number of communities: 24
## Elapsed time: 1 seconds
## Modularity Optimizer version 1.3.0 by Ludo Waltman and Nees Jan van Eck
## 
## Number of nodes: 17343
## Number of edges: 615265
## 
## Running Louvain algorithm...
## Maximum modularity in 10 random starts: 0.9058
## Number of communities: 29
## Elapsed time: 1 seconds
## Modularity Optimizer version 1.3.0 by Ludo Waltman and Nees Jan van Eck
## 
## Number of nodes: 17343
## Number of edges: 615265
## 
## Running Louvain algorithm...
## Maximum modularity in 10 random starts: 0.8937
## Number of communities: 30
## Elapsed time: 1 seconds
## Modularity Optimizer version 1.3.0 by Ludo Waltman and Nees Jan van Eck
## 
## Number of nodes: 17343
## Number of edges: 615265
## 
## Running Louvain algorithm...
## Maximum modularity in 10 random starts: 0.8820
## Number of communities: 33
## Elapsed time: 1 seconds
## Modularity Optimizer version 1.3.0 by Ludo Waltman and Nees Jan van Eck
## 
## Number of nodes: 17343
## Number of edges: 615265
## 
## Running Louvain algorithm...
## Maximum modularity in 10 random starts: 0.8734
## Number of communities: 34
## Elapsed time: 2 seconds
## Modularity Optimizer version 1.3.0 by Ludo Waltman and Nees Jan van Eck
## 
## Number of nodes: 17343
## Number of edges: 615265
## 
## Running Louvain algorithm...
## Maximum modularity in 10 random starts: 0.8644
## Number of communities: 36
## Elapsed time: 2 seconds
```

Based on these data, we found resolution 1.2 to be optimal to distinguish between NPC populations

```
SCT_obj$clusters <- SCT_obj$int_20_snn_res.1.2
UMAPPlot(SCT_obj, group.by = "clusters")
```

Adding irradiation status metadata on samples as a “Status” metadata column

```
# Creating named vector with status as values and associated sample names as names
Status_info <- c("0Gy","0Gy", "4Gy","4Gy")
Sample_names <- as.factor(c("CTRL_1", "CTRL_2", "4Gy_1", "4Gy_2"))
names(Status_info) <- Sample_names

# Add Status as metadata
Idents(SCT_obj) <- "orig.ident"
SCT_obj <- RenameIdents(SCT_obj, Status_info)
SCT_obj$Status <- Idents(SCT_obj)

saveRDS(object = SCT_obj, file = "Objects/SCT_obj.rds")
```

```
sessionInfo()
```

```
## R version 4.1.0 (2021-05-18)
## Platform: x86_64-conda-linux-gnu (64-bit)
## Running under: Ubuntu 20.04.4 LTS
## 
## Matrix products: default
## BLAS/LAPACK: /opt/rstudio-server_conda/conda/envs/rstudio-server_4.1.0/lib/libopenblasp-r0.3.15.so
## 
## locale:
##  [1] LC_CTYPE=en_US.UTF-8       LC_NUMERIC=C              
##  [3] LC_TIME=en_US.UTF-8        LC_COLLATE=en_US.UTF-8    
##  [5] LC_MONETARY=en_US.UTF-8    LC_MESSAGES=en_US.UTF-8   
##  [7] LC_PAPER=en_US.UTF-8       LC_NAME=C                 
##  [9] LC_ADDRESS=C               LC_TELEPHONE=C            
## [11] LC_MEASUREMENT=en_US.UTF-8 LC_IDENTIFICATION=C       
## 
## attached base packages:
## [1] stats     graphics  grDevices utils     datasets  methods   base     
## 
## other attached packages:
##  [1] sp_1.4-7           SeuratObject_4.1.0 Seurat_4.1.1       patchwork_1.1.2   
##  [5] clustree_0.5.0     ggraph_2.0.5       forcats_0.5.2      stringr_1.4.1     
##  [9] dplyr_1.0.10       purrr_0.3.5        readr_2.1.2        tidyr_1.2.1       
## [13] tibble_3.1.8       ggplot2_3.4.0      tidyverse_1.3.2   
## 
## loaded via a namespace (and not attached):
##   [1] readxl_1.4.1          backports_1.4.1       plyr_1.8.7           
##   [4] igraph_1.3.5          lazyeval_0.2.2        splines_4.1.0        
##   [7] listenv_0.8.0         scattermore_0.8       digest_0.6.31        
##  [10] htmltools_0.5.4       viridis_0.6.2         fansi_1.0.3          
##  [13] magrittr_2.0.3        tensor_1.5            googlesheets4_1.0.0  
##  [16] cluster_2.1.3         ROCR_1.0-11           tzdb_0.3.0           
##  [19] globals_0.15.0        graphlayouts_0.8.0    modelr_0.1.8         
##  [22] matrixStats_0.62.0    spatstat.sparse_2.1-1 colorspace_2.0-3     
##  [25] rvest_1.0.2           ggrepel_0.9.2         haven_2.5.0          
##  [28] xfun_0.31             crayon_1.5.2          jsonlite_1.8.4       
##  [31] progressr_0.10.0      spatstat.data_2.2-0   survival_3.3-1       
##  [34] zoo_1.8-10            glue_1.6.2            polyclip_1.10-4      
##  [37] gtable_0.3.1          gargle_1.2.0          leiden_0.4.2         
##  [40] future.apply_1.9.0    abind_1.4-5           scales_1.2.1         
##  [43] DBI_1.1.2             spatstat.random_2.2-0 miniUI_0.1.1.1       
##  [46] Rcpp_1.0.9            viridisLite_0.4.1     xtable_1.8-4         
##  [49] reticulate_1.22       spatstat.core_2.4-4   htmlwidgets_1.5.4    
##  [52] httr_1.4.4            RColorBrewer_1.1-3    ellipsis_0.3.2       
##  [55] ica_1.0-2             pkgconfig_2.0.3       farver_2.1.1         
##  [58] uwot_0.1.11           deldir_1.0-6          sass_0.4.4           
##  [61] dbplyr_2.1.1          utf8_1.2.2            labeling_0.4.2       
##  [64] tidyselect_1.2.0      rlang_1.0.6           reshape2_1.4.4       
##  [67] later_1.3.0           munsell_0.5.0         cellranger_1.1.0     
##  [70] tools_4.1.0           cachem_1.0.6          cli_3.4.1            
##  [73] generics_0.1.3        broom_0.8.0           ggridges_0.5.3       
##  [76] evaluate_0.15         fastmap_1.1.0         goftest_1.2-3        
##  [79] yaml_2.3.6            knitr_1.39            fs_1.5.2             
##  [82] fitdistrplus_1.1-8    tidygraph_1.2.1       RANN_2.6.1           
##  [85] nlme_3.1-157          pbapply_1.5-0         future_1.25.0        
##  [88] mime_0.12             xml2_1.3.3            compiler_4.1.0       
##  [91] rstudioapi_0.13       plotly_4.10.0         png_0.1-7            
##  [94] spatstat.utils_2.3-1  reprex_2.0.1          tweenr_1.0.2         
##  [97] bslib_0.4.2           stringi_1.7.8         highr_0.9            
## [100] RSpectra_0.16-1       rgeos_0.5-9           lattice_0.20-45      
## [103] Matrix_1.4-1          vctrs_0.5.1           pillar_1.8.1         
## [106] lifecycle_1.0.3       spatstat.geom_2.4-0   lmtest_0.9-40        
## [109] jquerylib_0.1.4       RcppAnnoy_0.0.19      data.table_1.14.2    
## [112] cowplot_1.1.1         irlba_2.3.5           httpuv_1.6.5         
## [115] R6_2.5.1              promises_1.2.0.1      KernSmooth_2.23-20   
## [118] gridExtra_2.3         parallelly_1.31.1     codetools_0.2-18     
## [121] MASS_7.3-57           assertthat_0.2.1      withr_2.5.0          
## [124] sctransform_0.3.3     mgcv_1.8-40           parallel_4.1.0       
## [127] hms_1.1.1             rpart_4.1.16          grid_4.1.0           
## [130] rmarkdown_2.14        googledrive_2.0.0     Rtsne_0.16           
## [133] ggforce_0.3.3         shiny_1.7.1           lubridate_1.8.0
```
