## Supplementary material for "Switching of RNA splicing regulators in immature neuroblasts: a key step in adult neurogenesis": script_R_4: 4_Cell_Type_annotation.html


### Cell type annotation

Libraries

```
library(tidyverse)
```

```
## ── Attaching packages ─────────────────────────────────────── tidyverse 1.3.2 ──
## ✔ ggplot2 3.4.0      ✔ purrr   0.3.5 
## ✔ tibble  3.1.8      ✔ dplyr   1.0.10
## ✔ tidyr   1.2.1      ✔ stringr 1.4.1 
## ✔ readr   2.1.2      ✔ forcats 0.5.2 
## ── Conflicts ────────────────────────────────────────── tidyverse_conflicts() ──
## ✖ dplyr::filter() masks stats::filter()
## ✖ dplyr::lag()    masks stats::lag()
```

```
library(dplyr)
library(clustree)
```

```
## Loading required package: ggraph
```

```
library(patchwork)
library(Seurat, lib.loc = "/opt/rstudio-server_conda/conda/envs/rstudio-server_4.1.0/lib/R/library/")
```

```
## Attaching SeuratObject
## Attaching sp
```

Import objects

```
SCT_obj <- readRDS("Objects/SCT_obj.rds")
Biomart_annotated <- read.csv("Analysis_tools/Biomart_annotated.csv")
```

Clear unused resolutions from metadata

```
metadata_to_clear <- paste0("int_20_snn_res.", seq(0.2,2,0.2))
SCT_obj@meta.data[,metadata_to_clear] <- NULL
```

We used selected genes from the literature to characterize the various clusters

```
Genes_to_show <- c("Slc1a3", "Aqp4", "Gfap", "Vcam1", "S100a6", "Egfr", "Ascl1", "Top2a", "Mki67", "Dcx", "Robo2", "Slc17a6", "Pdgfra", "Il33", "Mog", "S100b", "Dnah11", "Acta2", "Flt1", "Pecam1", "Pdgfrb", "Des", "Vtn", "Fyn", "P2ry12", "Cd14", "Cd3e")

clusters_order <- 
  c("29", "4", "13", "17","20", "10", "5","12", "15", "8",
    "19","1", "3", "7","9", "30","27","16","24","23", "6","14", 
    "21", "28","0", "2","11", "18", "25", "31", "22","26","32")

SCT_obj$clusters <- factor(SCT_obj$clusters, levels = clusters_order)

Idents(SCT_obj) <- "clusters"
```

Based on this gene expression patterns, we assigned cell types manually for every cluster

```
cell_annotations <- c(
  "Astrocytes",
  "Astrocytes",
  "Astrocytes",
  "NSCs",
  "TAPs",
  "Cycling Prog.",
  "Cycling Prog.",
  "Cycling Prog.",
  "Cycling Prog.",
  "Cycling Prog.",
  "Neuroblasts",
  "Neuroblasts",
  "Neuroblasts",
  "Neuroblasts",
  "Neuroblasts",
  "Neurons",
  "OPCs",
  "MFOLs",
  "MFOLs",
  "Ependymal",
  "Endothelial",
  "Pericytes",
  "Pericytes",
  "Microglia",
  "Microglia",
  "Microglia",
  "Microglia",
  "Microglia",
  "Microglia",
  "Immune",
  "Immune",
  "Immune",
  "Immune"
)

names(cell_annotations) <- as.factor(clusters_order)

Idents(SCT_obj) <- "clusters"

SCT_obj <- RenameIdents(SCT_obj, cell_annotations)

SCT_obj <- AddMetaData(SCT_obj, 
                       metadata =  Idents(SCT_obj), 
                       col.name = "Cell_Type")
SCT_obj$Cell_Type <- factor(SCT_obj$Cell_Type, levels = unique(cell_annotations))

UMAPPlot(SCT_obj, group.by = "Cell_Type", label = T) + NoLegend()
```

Selecting NPC cells only Cluster 29 was excluded from NPC-restricted analyses as it shared features from astrocytes and microglia

```
#Creating NPC metadata for easy selection (using subset(object, NPC == "NPC"))
cell_annotations[c(1,16:33)] <- "Micro-environment"
cell_annotations[2:15] <- "NPC"

Idents(SCT_obj) <- "clusters"

SCT_obj <- RenameIdents(SCT_obj, cell_annotations)

SCT_obj <- AddMetaData(SCT_obj, 
                       metadata =  Idents(SCT_obj), 
                       col.name = "NPC")

#Creating NPC-restricted ordered cluster metadata
cell_annotations[c(1,16:33)] <- NA
cell_annotations[2:15] <- clusters_order[2:15]

Idents(SCT_obj) <- "clusters"

SCT_obj <- RenameIdents(SCT_obj, cell_annotations)

SCT_obj <- AddMetaData(SCT_obj, 
                       metadata =  Idents(SCT_obj), 
                       col.name = "clusters_NPC")
SCT_obj$clusters_NPC <- factor(SCT_obj$clusters_NPC, levels = clusters_order[2:15])

UMAPPlot(SCT_obj, group.by = "clusters_NPC", label = T) + NoLegend()
```

```
saveRDS(object = SCT_obj, file = "Objects/SCT_obj.rds")
sessionInfo()
```

```
## R version 4.1.0 (2021-05-18)
## Platform: x86_64-conda-linux-gnu (64-bit)
## Running under: Ubuntu 20.04.4 LTS
## 
## Matrix products: default
## BLAS/LAPACK: /opt/rstudio-server_conda/conda/envs/rstudio-server_4.1.0/lib/libopenblasp-r0.3.15.so
## 
## locale:
##  [1] LC_CTYPE=en_US.UTF-8       LC_NUMERIC=C              
##  [3] LC_TIME=en_US.UTF-8        LC_COLLATE=en_US.UTF-8    
##  [5] LC_MONETARY=en_US.UTF-8    LC_MESSAGES=en_US.UTF-8   
##  [7] LC_PAPER=en_US.UTF-8       LC_NAME=C                 
##  [9] LC_ADDRESS=C               LC_TELEPHONE=C            
## [11] LC_MEASUREMENT=en_US.UTF-8 LC_IDENTIFICATION=C       
## 
## attached base packages:
## [1] stats     graphics  grDevices utils     datasets  methods   base     
## 
## other attached packages:
##  [1] sp_1.4-7           SeuratObject_4.1.0 Seurat_4.1.1       patchwork_1.1.2   
##  [5] clustree_0.5.0     ggraph_2.0.5       forcats_0.5.2      stringr_1.4.1     
##  [9] dplyr_1.0.10       purrr_0.3.5        readr_2.1.2        tidyr_1.2.1       
## [13] tibble_3.1.8       ggplot2_3.4.0      tidyverse_1.3.2   
## 
## loaded via a namespace (and not attached):
##   [1] readxl_1.4.1          backports_1.4.1       plyr_1.8.7           
##   [4] igraph_1.3.5          lazyeval_0.2.2        splines_4.1.0        
##   [7] listenv_0.8.0         scattermore_0.8       digest_0.6.31        
##  [10] htmltools_0.5.4       viridis_0.6.2         fansi_1.0.3          
##  [13] magrittr_2.0.3        tensor_1.5            googlesheets4_1.0.0  
##  [16] cluster_2.1.3         ROCR_1.0-11           tzdb_0.3.0           
##  [19] globals_0.15.0        graphlayouts_0.8.0    modelr_0.1.8         
##  [22] matrixStats_0.62.0    spatstat.sparse_2.1-1 colorspace_2.0-3     
##  [25] rvest_1.0.2           ggrepel_0.9.2         haven_2.5.0          
##  [28] xfun_0.31             crayon_1.5.2          jsonlite_1.8.4       
##  [31] progressr_0.10.0      spatstat.data_2.2-0   survival_3.3-1       
##  [34] zoo_1.8-10            glue_1.6.2            polyclip_1.10-4      
##  [37] gtable_0.3.1          gargle_1.2.0          leiden_0.4.2         
##  [40] future.apply_1.9.0    abind_1.4-5           scales_1.2.1         
##  [43] DBI_1.1.2             spatstat.random_2.2-0 miniUI_0.1.1.1       
##  [46] Rcpp_1.0.9            viridisLite_0.4.1     xtable_1.8-4         
##  [49] reticulate_1.22       spatstat.core_2.4-4   htmlwidgets_1.5.4    
##  [52] httr_1.4.4            RColorBrewer_1.1-3    ellipsis_0.3.2       
##  [55] ica_1.0-2             pkgconfig_2.0.3       farver_2.1.1         
##  [58] uwot_0.1.11           deldir_1.0-6          sass_0.4.4           
##  [61] dbplyr_2.1.1          utf8_1.2.2            labeling_0.4.2       
##  [64] tidyselect_1.2.0      rlang_1.0.6           reshape2_1.4.4       
##  [67] later_1.3.0           munsell_0.5.0         cellranger_1.1.0     
##  [70] tools_4.1.0           cachem_1.0.6          cli_3.4.1            
##  [73] generics_0.1.3        broom_0.8.0           ggridges_0.5.3       
##  [76] evaluate_0.15         fastmap_1.1.0         goftest_1.2-3        
##  [79] yaml_2.3.6            knitr_1.39            fs_1.5.2             
##  [82] fitdistrplus_1.1-8    tidygraph_1.2.1       RANN_2.6.1           
##  [85] nlme_3.1-157          pbapply_1.5-0         future_1.25.0        
##  [88] mime_0.12             xml2_1.3.3            compiler_4.1.0       
##  [91] rstudioapi_0.13       plotly_4.10.0         png_0.1-7            
##  [94] spatstat.utils_2.3-1  reprex_2.0.1          tweenr_1.0.2         
##  [97] bslib_0.4.2           stringi_1.7.8         highr_0.9            
## [100] rgeos_0.5-9           lattice_0.20-45       Matrix_1.4-1         
## [103] vctrs_0.5.1           pillar_1.8.1          lifecycle_1.0.3      
## [106] spatstat.geom_2.4-0   lmtest_0.9-40         jquerylib_0.1.4      
## [109] RcppAnnoy_0.0.19      data.table_1.14.2     cowplot_1.1.1        
## [112] irlba_2.3.5           httpuv_1.6.5          R6_2.5.1             
## [115] promises_1.2.0.1      KernSmooth_2.23-20    gridExtra_2.3        
## [118] parallelly_1.31.1     codetools_0.2-18      MASS_7.3-57          
## [121] assertthat_0.2.1      withr_2.5.0           sctransform_0.3.3    
## [124] mgcv_1.8-40           parallel_4.1.0        hms_1.1.1            
## [127] rpart_4.1.16          grid_4.1.0            rmarkdown_2.14       
## [130] googledrive_2.0.0     Rtsne_0.16            ggforce_0.3.3        
## [133] shiny_1.7.1           lubridate_1.8.0
```
