## Supplementary material for "Switching of RNA splicing regulators in immature neuroblasts: a key step in adult neurogenesis": Script_R_5: 5_Population_scoring_UCell.html

Population identification with microarray signatures matching


### Population identification with microarray signatures matching

In this script, we load the population signatures defined using our microarrays experiments We then select the top genes for each signature, and calculate a UCell score for each signature in each cell. Cells are individually attributed to populations based on the highest score amongst signatures.

Libraries

```
library(tidyverse)
```

```
## ── Attaching packages ─────────────────────────────────────── tidyverse 1.3.2 ──
## ✔ ggplot2 3.4.0      ✔ purrr   0.3.5 
## ✔ tibble  3.1.8      ✔ dplyr   1.0.10
## ✔ tidyr   1.2.1      ✔ stringr 1.4.1 
## ✔ readr   2.1.2      ✔ forcats 0.5.2 
## ── Conflicts ────────────────────────────────────────── tidyverse_conflicts() ──
## ✖ dplyr::filter() masks stats::filter()
## ✖ dplyr::lag()    masks stats::lag()
```

```
library(dplyr)
library(patchwork)
library(Seurat, lib.loc = "/opt/rstudio-server_conda/conda/envs/rstudio-server_4.1.0/lib/R/library/")
```

```
## Attaching SeuratObject
## Attaching sp
```

```
library(readr)
library(UCell)
library(ggplot2)
library(circlize)
```

```
## ========================================
## circlize version 0.4.15
## CRAN page: https://cran.r-project.org/package=circlize
## Github page: https://github.com/jokergoo/circlize
## Documentation: https://jokergoo.github.io/circlize_book/book/
## 
## If you use it in published research, please cite:
## Gu, Z. circlize implements and enhances circular visualization
##   in R. Bioinformatics 2014.
## 
## This message can be suppressed by:
##   suppressPackageStartupMessages(library(circlize))
## ========================================
```

```
library(scales)
```

```
## 
## Attaching package: 'scales'
## 
## The following object is masked from 'package:purrr':
## 
##     discard
## 
## The following object is masked from 'package:readr':
## 
##     col_factor
```

```
library(openxlsx)
```

Loading required objects This analysis is focused on NPC populations, therefore we will use a subset of the base object

```
SCT_obj <- readRDS("Objects/SCT_obj.rds")
Biomart_annotated <- read.csv("Analysis_tools/Biomart_annotated.csv")
geneIDs <- # For easy calling of genes through external gene name
  Biomart_annotated[!duplicated(Biomart_annotated$external_gene_name),1]
names(geneIDs) <- Biomart_annotated[!duplicated(Biomart_annotated$external_gene_name),2]
```

Distinction of astrocytes and qNSC using UCell Score

Loading genesets from Cebrian-Silla et al. 2021, selecting Top 100 genes

```
Astrocytes_markers <- read.xlsx("All_inputs/Cebrian-Silla/elife_67436_supp1_v3.xlsx", sheet = 1)
B_cells_markers <- read.xlsx("All_inputs/Cebrian-Silla/elife_67436_supp1_v3.xlsx", sheet = 3)
Astro_vs_B <- list(geneIDs[Astrocytes_markers$genes][1:100], geneIDs[B_cells_markers$genes][1:100])
names(Astro_vs_B) <- c("Astrocytes", "B_Cells")
```

Selecting astrocytes and B cells

```
clusters_of_Astro <- c("4", "13")

Astro_obj <- subset(SCT_obj, clusters %in% clusters_of_Astro)
```

Running UCell

```
DefaultAssay(Astro_obj) <- "SCT" # Run on normalized counts
method_UC <- "_UC_Score"
Astro_obj <-
  AddModuleScore_UCell(
    obj = Astro_obj,
    features = Astro_vs_B,
    name =  method_UC,  
    assay = "SCT"
  )#suffix for metadata
```

```
## Warning in check_genes(matrix, features): The following genes were not found and will be imputed to exp=0:
## * NA,ENSMUSG00000099869
```

Creating new metadata column containing the name of the highest score

```
Astro_obj$Astro_vs_B <- 
  colnames(Astro_obj@meta.data[,
                             paste0(names(Astro_vs_B), method_UC)])[
                     apply(Astro_obj@meta.data[,
                                             paste0(names(Astro_vs_B),
                                                    method_UC)], 
                           1,
                           function(vec) {which(vec == max(vec))})
                     ]
```

Metadata addition

```
SCT_obj <- AddMetaData(SCT_obj, 
                       metadata = Astro_obj$Astro_vs_B, 
                       col.name = "Astro_vs_B")
UMAPPlot(SCT_obj, group.by = "Astro_vs_B", label = T) + NoLegend()
```

Loading population signatures from bulk transcriptome data (Supp. Material #)

```
names_sig <- c("qNSC", "aNSC", "TAP", "iNB", "mNB")

Population_signatures <- lapply(X = paste0("All_inputs/Microarray_signatures/", 
                                           names_sig,
                                           "_signature.xlsx"),
                                FUN = read.xlsx)
```

Selection of genes present in SCT assay and determination of top 100 highest fold-change

```
Population_signatures <- 
  lapply(X = 1:length(names_sig),
         FUN = function(signature_nb){
           signature_list <- merge(x = Population_signatures[[signature_nb]],
                                   y = Biomart_annotated,
                                   by.x = "gene",
                                   by.y = "external_gene_name",
                                   all.x = TRUE) %>%
             filter(!is.na(description)) %>% # Use subset and select
             arrange(desc(FC))
           signature_list$...1 <- NULL
           signature_list$Population <- names_sig[[signature_nb]]
           signature_list <- 
             filter(signature_list, ensembl_gene_id %in%
                      rownames(SCT_obj@assays[["SCT"]]@data)) %>%
             arrange(desc(FC))
           write.csv(signature_list[1:100,], row.names = FALSE,
                     file = paste0("Outputs/Population_signatures_top100/Top100_", 
                                   names_sig[[signature_nb]], 
                                   "_signature.csv")) #Returns the provided gene lists
           return(signature_list[1:100,"ensembl_gene_id"])
         })

names_sig2 <- paste0("s_", names_sig)
names(Population_signatures) <- names_sig2
```

Calculation of UCell Score for each signature (s-TAP has only 99 genes, producing a NA warning for position 100)

```
NPC_obj <- subset(SCT_obj, NPC == "NPC") 

DefaultAssay(NPC_obj) <- "SCT"

NPC_obj <-
  AddModuleScore_UCell(
    obj = NPC_obj,
    features = Population_signatures,
    name = method_UC,
    assay = "SCT",
    slot = "counts")
```

```
## Warning in check_genes(matrix, features): The following genes were not found and will be imputed to exp=0:
## * NA
```

Retrieve UCell scores and ID from object, then assign cell to population based on highest score

```
NPC_obj$Identity <-  
  colnames(NPC_obj@meta.data[,
                             paste0(names_sig2, method_UC)])[
                               apply(NPC_obj@meta.data[,
                                                       paste0(names_sig2,
                                                              method_UC)],
                                     1,
                                     function(vec) {which(vec == max(vec))})
                               ]


Identity_convert <- paste0("s-", names_sig)
names(Identity_convert) <- paste0(names_sig2, method_UC)

Idents(NPC_obj) <- "Identity"
NPC_obj <- RenameIdents(NPC_obj, Identity_convert)

NPC_obj$Identity <- Idents(NPC_obj)
NPC_obj$Identity <- factor(Idents(NPC_obj), levels = Identity_convert)

SCT_obj <- AddMetaData(SCT_obj,
                       metadata = NPC_obj$Identity, 
                       col.name = "Identity")
UMAPPlot(SCT_obj, group.by = "Identity", label = T) + NoLegend()
```

```
saveRDS(SCT_obj, file = "Objects/SCT_obj.rds")
sessionInfo()
```

```
## R version 4.1.0 (2021-05-18)
## Platform: x86_64-conda-linux-gnu (64-bit)
## Running under: Ubuntu 20.04.4 LTS
## 
## Matrix products: default
## BLAS/LAPACK: /opt/rstudio-server_conda/conda/envs/rstudio-server_4.1.0/lib/libopenblasp-r0.3.15.so
## 
## locale:
##  [1] LC_CTYPE=en_US.UTF-8       LC_NUMERIC=C              
##  [3] LC_TIME=en_US.UTF-8        LC_COLLATE=en_US.UTF-8    
##  [5] LC_MONETARY=en_US.UTF-8    LC_MESSAGES=en_US.UTF-8   
##  [7] LC_PAPER=en_US.UTF-8       LC_NAME=C                 
##  [9] LC_ADDRESS=C               LC_TELEPHONE=C            
## [11] LC_MEASUREMENT=en_US.UTF-8 LC_IDENTIFICATION=C       
## 
## attached base packages:
## [1] stats     graphics  grDevices utils     datasets  methods   base     
## 
## other attached packages:
##  [1] openxlsx_4.2.5.1   scales_1.2.1       circlize_0.4.15    UCell_1.3.1       
##  [5] sp_1.4-7           SeuratObject_4.1.0 Seurat_4.1.1       patchwork_1.1.2   
##  [9] forcats_0.5.2      stringr_1.4.1      dplyr_1.0.10       purrr_0.3.5       
## [13] readr_2.1.2        tidyr_1.2.1        tibble_3.1.8       ggplot2_3.4.0     
## [17] tidyverse_1.3.2   
## 
## loaded via a namespace (and not attached):
##   [1] readxl_1.4.1          backports_1.4.1       plyr_1.8.7           
##   [4] igraph_1.3.1          lazyeval_0.2.2        splines_4.1.0        
##   [7] BiocParallel_1.26.2   listenv_0.8.0         scattermore_0.8      
##  [10] digest_0.6.31         htmltools_0.5.4       fansi_1.0.3          
##  [13] magrittr_2.0.3        tensor_1.5            googlesheets4_1.0.0  
##  [16] cluster_2.1.3         ROCR_1.0-11           tzdb_0.3.0           
##  [19] globals_0.15.0        modelr_0.1.8          matrixStats_0.62.0   
##  [22] spatstat.sparse_2.1-1 colorspace_2.0-3      rvest_1.0.2          
##  [25] ggrepel_0.9.1         haven_2.5.0           xfun_0.31            
##  [28] crayon_1.5.2          jsonlite_1.8.4        progressr_0.10.0     
##  [31] spatstat.data_2.2-0   survival_3.3-1        zoo_1.8-10           
##  [34] glue_1.6.2            polyclip_1.10-0       gtable_0.3.1         
##  [37] gargle_1.2.0          leiden_0.4.2          shape_1.4.6          
##  [40] future.apply_1.9.0    abind_1.4-5           DBI_1.1.2            
##  [43] spatstat.random_2.2-0 miniUI_0.1.1.1        Rcpp_1.0.8.3         
##  [46] viridisLite_0.4.0     xtable_1.8-4          reticulate_1.22      
##  [49] spatstat.core_2.4-4   htmlwidgets_1.5.4     httr_1.4.4           
##  [52] RColorBrewer_1.1-3    ellipsis_0.3.2        ica_1.0-2            
##  [55] farver_2.1.1          pkgconfig_2.0.3       uwot_0.1.11          
##  [58] sass_0.4.4            dbplyr_2.1.1          deldir_1.0-6         
##  [61] utf8_1.2.2            labeling_0.4.2        tidyselect_1.2.0     
##  [64] rlang_1.0.6           reshape2_1.4.4        later_1.3.0          
##  [67] munsell_0.5.0         cellranger_1.1.0      tools_4.1.0          
##  [70] cachem_1.0.6          cli_3.4.1             generics_0.1.3       
##  [73] broom_0.8.0           ggridges_0.5.3        evaluate_0.15        
##  [76] fastmap_1.1.0         yaml_2.3.6            goftest_1.2-3        
##  [79] knitr_1.39            fs_1.5.2              fitdistrplus_1.1-8   
##  [82] zip_2.2.0             RANN_2.6.1            pbapply_1.5-0        
##  [85] future_1.25.0         nlme_3.1-157          mime_0.12            
##  [88] xml2_1.3.3            compiler_4.1.0        rstudioapi_0.13      
##  [91] plotly_4.10.0         png_0.1-7             spatstat.utils_2.3-1 
##  [94] reprex_2.0.1          bslib_0.4.2           stringi_1.7.8        
##  [97] highr_0.9             rgeos_0.5-9           lattice_0.20-45      
## [100] Matrix_1.4-1          vctrs_0.5.1           pillar_1.8.1         
## [103] lifecycle_1.0.3       GlobalOptions_0.1.2   spatstat.geom_2.4-0  
## [106] lmtest_0.9-40         jquerylib_0.1.4       RcppAnnoy_0.0.19     
## [109] data.table_1.14.2     cowplot_1.1.1         irlba_2.3.5          
## [112] httpuv_1.6.5          R6_2.5.1              promises_1.2.0.1     
## [115] KernSmooth_2.23-20    gridExtra_2.3         parallelly_1.31.1    
## [118] codetools_0.2-18      MASS_7.3-57           assertthat_0.2.1     
## [121] withr_2.5.0           sctransform_0.3.3     mgcv_1.8-40          
## [124] parallel_4.1.0        hms_1.1.1             grid_4.1.0           
## [127] rpart_4.1.16          rmarkdown_2.14        googledrive_2.0.0    
## [130] Rtsne_0.16            shiny_1.7.1           lubridate_1.8.0
```
